# Segmented filamentous bacteria undergo a structural transition at their adhesive tip during unicellular to filament development

**DOI:** 10.1101/2025.04.09.647586

**Authors:** Ana Raquel Cruz, Benedikt Wimmer, Teck Hui Teo, Gérard Pehau-Arnaudet, Jean-Marie Winter, Anastasia Gazi, Pierre Lafaye, Sébastien Brier, Agnès Legrand, Anna Dubrovsky-Gaupp, Marion Bérard, Gabriel Aymé, Anna Sartori-Rupp, Ohad Medalia, Pamela Schnupf

## Abstract

Segmented filamentous bacteria (SFB) are intestinal commensals that promote immune system development and pathogen protection through intimate attachment to the ileal epithelium. Attachment occurs via the tip of unicellular teardrop-shaped SFB, called intracellular offsprings (IOs), before outgrowth into filaments. To characterize this critical stage of the SFB life-cycle, we imaged SFB using cryo-electron microscopy and tomography. IOs were surrounded by a repetitive surface (S)-layer that became replaced by disordered hair-like structures uniquely at the tip. Upon outgrowth into filaments, the S-layer was exchanged for a morphologically distinct repetitive hair-like layer. The bacterial structures and morphological transition were conserved across SFB from mouse and rat origin, while growth of mouse-SFB under non-attachment conditions in a heterologous host affected the SFB tip length and relative proportion of the tip stages. Moreover, the filament tip displayed surface exposure of the major Th17 antigen, a ubiquitous cell wall protein, underscoring the unique properties of the adhesive tip. This study identifies a novel IO-specific S-layer and reveals a conserved developmental transition of the SFB tip surface including the transient appearance of structures consistent in location and timing with being involved in host cell attachment.

## Introduction

The intimate interactions between the intestinal microbiota and its host plays a crucial role in shaping host physiology including the maturation of the immune system^1^. Segmented Filamentous Bacteria (SFB) are ubiquitous commensals in vertebrates, but most extensively studied in mice where they strongly stimulate both innate and adaptive immunity^2–7^ and provide protection against intestinal and respiratory pathogens^3,8,9^. However, these *Clostridium*-related monoderm bacteria can also be harmful by exacerbating symptoms severity in a number of disease models^10^. Despite their immunomodulatory roles, SFB remain poorly studied due to technical constraints such as limited *in vitro* culturing^11^ and a lack of genetic tools.

Unlike most intestinal commensals, SFB are part of the more restricted mucosa-associated microbiota^1^. To establish their niche, teardrop-shaped unicellular SFB, called intracellular offsprings (IOs), attach to ileal epithelial cells and cells overlaying Peyer’s patches^12^. IOs attach using their tip, a unique hook-like structure, and elongate to form segmented filaments that can reach over 100 µm in length^11,12^. Filament differentiation leads to the formation of two new IOs per bacterial segment and eventual IO release at the distal end^12,13^.

The host-specific interaction^14,15^ mediated by the SFB tip leads to actin re-organization at the attachment site without evidence of SFB penetration beyond the terminal web or apparent plasma membrane disruption^16–18^. Endocytic vesicle formation near the attachment site results in the uptake of the major SFB antigen of the T helper 17 response, a hallmark host response to SFB colonization^2,3,7,18^. However, genome analyses of mouse- and rat-SFB have provided little insights regarding the composition of the SFB surface^19–22^. Typical cell wall binding motifs and domains are largely absent, as are typical surface layer (S-layer) proteins present in most Clostridia^19–22^. Some genes potentially involved in polysaccharide capsule formation were found^19–22^ but scanning and transmission electron microscopy have not identified bacterial supramolecular structures such as an S-layer or capsule in SFB^17,23^.

To obtain a more detailed view of the SFB surface, we imaged purified IOs and filaments from mouse- and rat-SFB in near native conditions using a combination of cryogenic electron microscopy (cryo-EM) and tomography (cryo-ET). We identified a number of intracellular structures but mainly focused on the SFB surface at the bacterial tip for which we describe a developmental structural transition during the outgrowth of unicellular IOs into filaments. This transition includes the replacement of an IO-specific S-layer by a repetitive hair-like layer, the appearance of tip structures with a potential role in attachment, and a difference in accessibility of the major Th17 antigen, a ubiquitous cell wall protein that is surface-exposed uniquely at the filament tip.

## Results

### SFB IOs are surrounded by an S-layer

We aimed to visualize the cellular structures that compose SFB in their native state and assess their conservation in SFB from different hosts. For this, we used cryo-EM and cryo-ET to image whole cells of SFB isolated from monocolonized mice (mouse-SFB) and rats (rat-SFB)^24,25^. We mainly focused on the SFB tip structure (**Fig. 1a**), which has a diameter of approximately 168.8 ± 28.9 nm (n = 10, **Supplementary Table 1**), and therefore can be studied *in toto*. A total of 130 mouse-SFB (85 IOs and 45 filaments) and 68 rat-SFB (48 IOs and 20 filaments) were imaged **(Supplementary Table 1)**. The SFB surface was characterized at both the bacterium’s tip and opposite end, here designated as back (**Fig. 1a-c**). Bacterial tip imaging included part of the SFB cell body which, together with the separate imaging of the IO back, was considered to represent the overall bacterial surface beyond the tip.

**Fig. 1.**
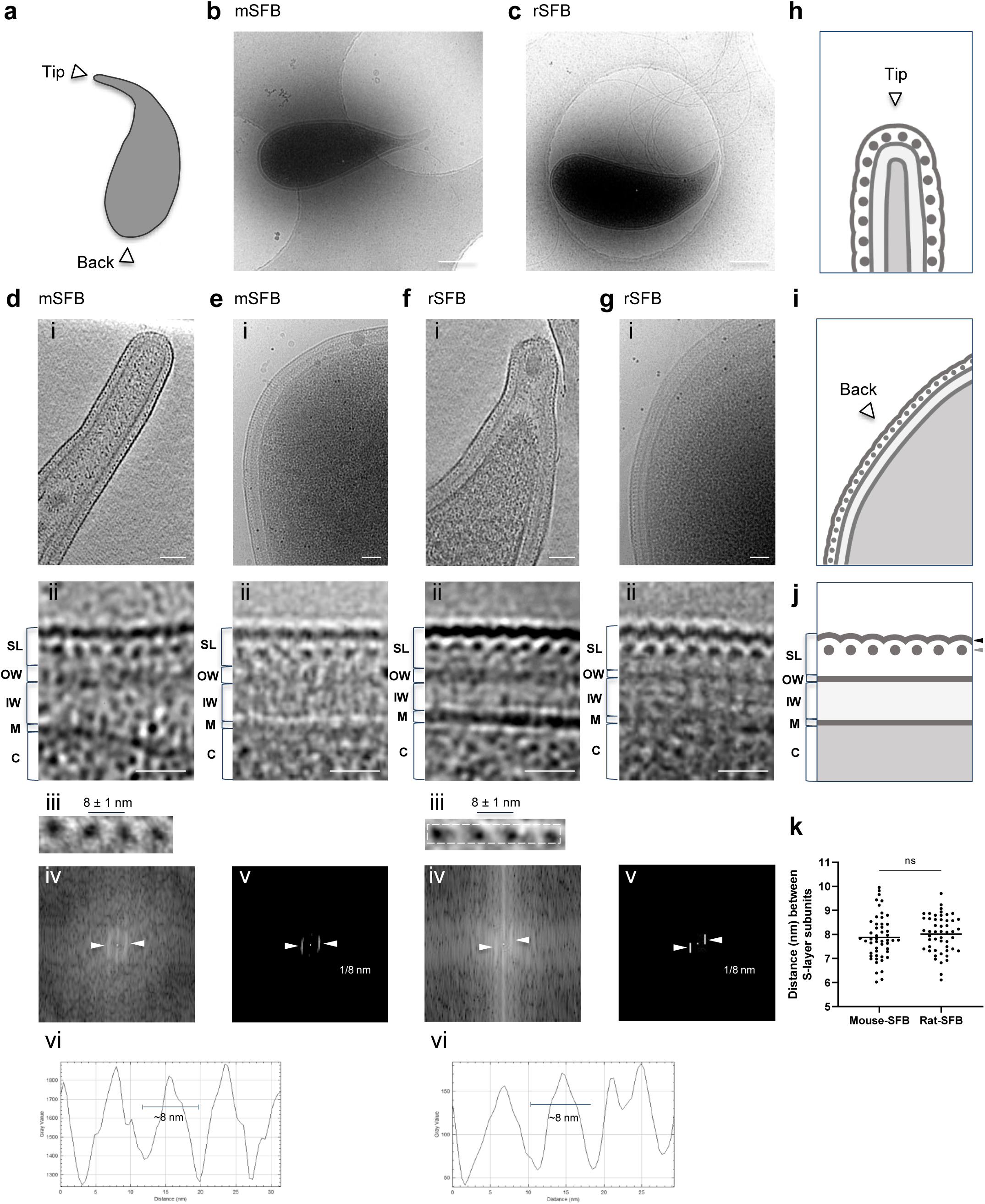
Identification of an S-layer in SFB IOs. **a**, Schematic representation of an SFB IO. The IO tip and back are indicated by white arrows. **b, c,** Projection image of: **(b)** a mouse-SFB IO and **(c)** a rat-SFB IO. **d,f(i)**, Tomographic slice showing the tip of: **d(i)** a mouse-SFB IO (EMD-52655) and **f(i)** a rat-SFB IO (EMD-52684). **e,g(i),** Projection images of the back of: **e(i)** a mouse- SFB IO and **g(i)** a rat-SFB IO. **d-g(ii),** Close ups of the layers present in SFB poles shown in d-g(i), respectively. **d,f(iii)**, Close ups of S-layer subunits from d,f(i). The average spacing and standard deviation between S-layer subunits is shown. The region from f(iii) used for repetitive pattern analysis is delimitated by a dashed line. **d,f(iv),** Fast Fourier transform, **d,f(v),** Maximum intensity peaks (indicated by white arrows) and their corresponding frequency, and **d,f(vi),** Plot profiles are shown for: **d(iii)** mouse-SFB and **f(iii)** rat-SFB S-layer subunits. **h, i, j,** Schematic representation of: **(h)** the tip, **(i)** the back and **(j)** the layers found in S-layer-containing IOs. The regions where crescent-like and globular subunits of the S-layer can be found are indicated by black and grey arrows, respectively. **k**, Distance between S-layer subunits in mouse-SFB and rat-SFB. Ten measurements were performed for each of the 5 SFB selected from 2 independent experiments. The mean is shown and the statistical significance was assessed using the t-test (ns: not significant). SL: Surface layer (S-layer), OW: outer wall zone, IW: inner wall zone, M: membrane, C: cytoplasm. mSFB: mouse-SFB, rSFB: rat-SFB. **Scale bars:** b/c: 500 nm; d-g(i): 50 nm; d-g(ii): 20 nm.

Exterior to the SFB membrane, a region of low electron density was followed by a more electron- dense region although this region was not always well defined (**Fig. 1d-g**). These two layers likely constitute the inner wall (IW) and outer wall (OW) zones of the cell wall (CW), as previously described for other monoderm bacteria such as *Bacillus subtilis*^26^.

Exterior to the cell wall, mouse-SFB IOs (**Fig. 1d/e**) and rat-SFB IOs (**Fig. 1f/g**) exhibited another electron-dense layer at both the tip and the back (**Fig. 1d-g (ii),h-j).** In cross-section view, this outer layer appeared to be composed of repetitive crescent-like elements and regularly-spaced globular densities (**Fig. 1d-g (ii),j)** that showed resemblance to the S-layer of *Caulobacter crescentus*, *Clostridioides difficile* and *Clostridium thermocellum*^27–29^. These data are consistent with the coverage of the entire IO surface by an S-layer, similarly to what is described for other bacteria^30^.

The distance between the repetitive globular subunits was measured manually, by Fourier spectra analysis, and by plot profile tracing on the side view of tomograms from the IO tip (**Fig. 1d /f(iii- vi),k)**. The three methods consistently estimated the globular subunit spacing around 8 nm. Spacing was similar for mouse-SFB (8 ± 1 nm, n = 50) and rat-SFB (8 ± 1 nm, n = 50) (*p*=0.4109, **Fig. 1k**) and was within the range of the typical center-to-center S-layer spacings (2.5 to 35 nm) of other bacteria^31^.

Top views of tomograms from the IO tip suggest that the electron dense subunits are arranged in rows **(Supplementary** Fig. 1a-d). Additionally, a potential six-fold symmetry was detected **(Supplementary** Fig. 1e,f). This apparent hexagonal organization was particularly clear in a tomogram of a rat-SFB IO with a ‘broken tip’ phenotype whereby the S-layer showed abrupt discontinuity and was absent at the tip end **(Supplementary** Fig. 1f). The lack of a clear S-layer symmetry seen in the top view of SFB tomograms may be explained by the need to accommodate differences in SFB tip curvature. Intact SFB tips appear to be composed of a twisted lattice **(Supplementary** Fig. 1a). The identification of a 6-fold symmetry in the rat IOs with a ‘broken tip’ phenotype may therefore be due to a more relaxed S-layer **(Supplementary** Fig. 1f), although a definitive identification of the S-layer symmetry will require a more detailed analysis beyond what whole SFB cryo-ET can provide. In comparison, in the closely-related species *C. difficile,* a p2- symmetric S-layer was observed^28^, while the morphologically similar *C. crescentus* is covered by a p6-symmetrical S-layer^27^. Together, these data reveal the presence of a repetitive S-layer that surrounds IOs and is conserved in SFB from varying hosts.

### Identification of intracellular and extracellular SFB features

During the SFB surface characterization, several intracellular and extracellular structures were identified (**Fig. 2a-g , Supplementary movies 1-5)**. These structures were mainly characterized at the IO tip. The presence of flagella at the concave side of SFB IOs, as previously described^32^, was confirmed for both mouse- and rat-SFB (**Fig. 2a,f /g(i), Supplementary** Fig. 2a-d**(i))**. A structure typical of a chemosensory array, predicted to be present by genome analysis^19,20^, was found localized at the convex side of the cell body, opposite to where flagella are anchored, in IOs from both hosts (**Fig. 2b , Supplementary** Fig. 2a-d**(ii))**. Vesicles with a median diameter of 51 nm for mouse-SFB (average diameter of 54 ± 15 nm, n = 15) and 41 nm for rat-SFB (average diameter of 62 ± 53 nm, n = 6) **(Supplementary Table 2)** were identified inside the tip of IOs and filaments (**Fig. 2a,f /g(ii), Supplementary** Fig. 2a**(v),b(iii),c/d(vi), Supplementary Table 1)**. When vesicles were present at the tip or when the cytoplasmic membrane did not fully extended into the tip, electron-dense lines parallel to the cell wall, here designated as tracks, were visible at the tip (**Fig. 2a and Supplementary** Fig. 2a**/b(v),c(vi) and d(vii))**. Tracks were commonly identified at the mouse-SFB tip but rarely at the rat-SFB tip **(Supplementary Table 1, Supplementary** Fig. 2d**(vii))**, possibly due to the shorter tip length of rat-SFB compared to that of mouse-SFB **(Supplementary** Fig. 3). Given their location, we hypothesize that these structures may play a role in maintaining tip shape or in anchoring the cytoplasmic membrane within the elongated tip.

**Fig. 2.**
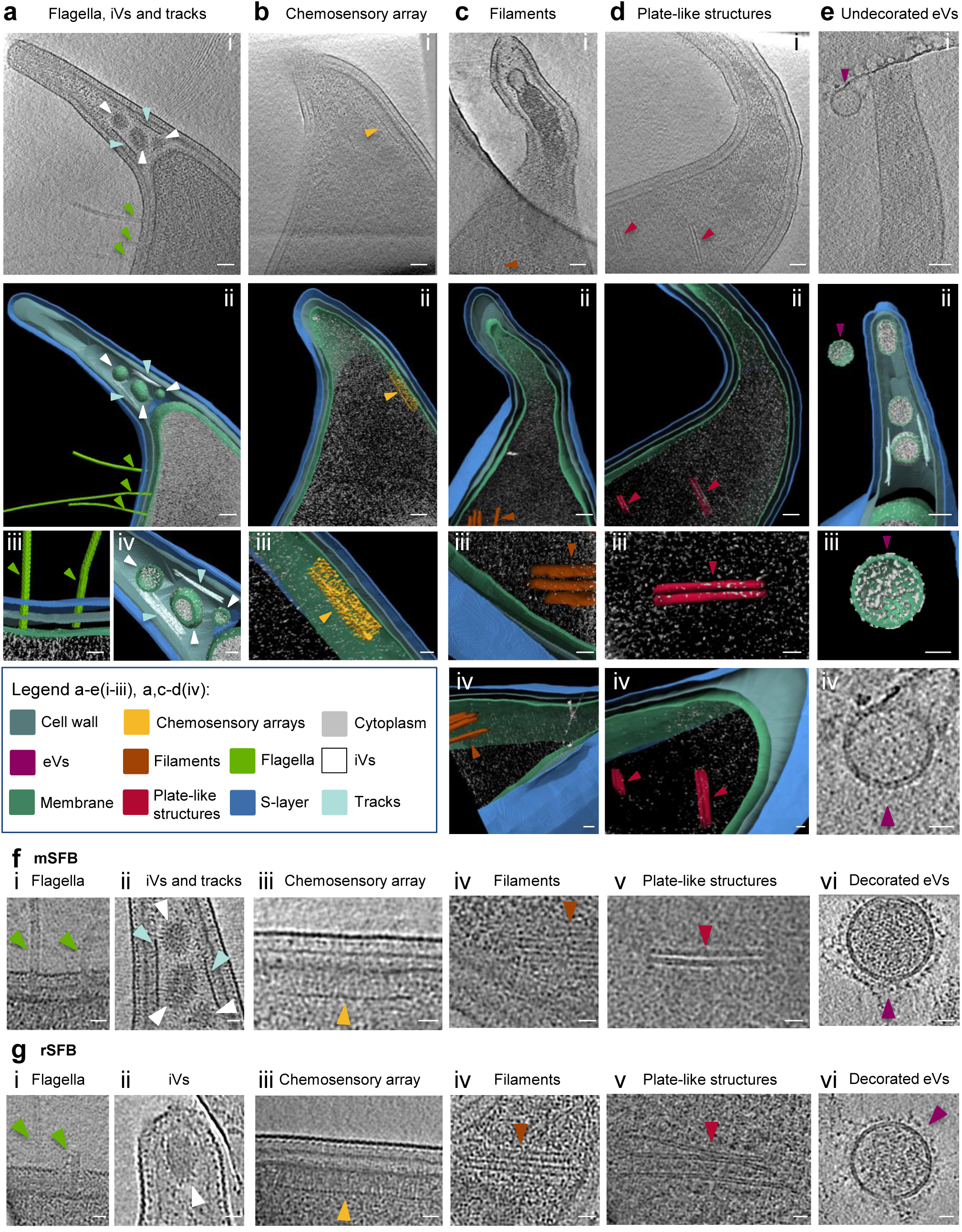
Description of intracellular and extracellular SFB features. **a-f,** Representative tomographic slices from mouse- SFB IOs containing: **(a, f(i-ii))** flagella, intracellular vesicles (iVs) and tracks, **(b, f(iii))** chemosensory array, **(c, f(iv))** filaments, **(d, f(v))** plate-like structures, **(e)** undecorated extracellular vesicles (eVs) and **f(vi)** decorated eVs. Representatives of the identified features are shown by arrows of the colours indicated in the legend. **a-e(i),** Tomographic slices from reconstructed tomograms EMD-52655 (Supplementary movie 1), EMD-52667 (Supplementary movie 2), EMD-52668 (Supplementary movie 3), EMD-52669 (Supplementary movie 4) and EMD-52670 (Supplementary movie 5) from SFB IOs with a length of: 2.7, 2.3, 2.0, 2.7 and 2.5 µm. **a-e(ii)** Segmentation rendering of the SFB tip and the identified intracellular and extracellular features present in the tomographic slices a-e(i). **a-e(iii) and a(iv),** Close ups of: **(a(iii))** flagellum, **(a(iv))** intracellular vesicles and tracks, **(b(iii))** chemosensory array, **(c(iii))** filaments, **(d(iii))** stacks and **(e(iii))** undecorated eVs from segmentation rendering shown in a-e(ii). **c-d(iv)**, close ups from the segmentation rendering shown in c-d(ii) to show the thickness in z of **c(iv)** filaments and **d(iv)** plate-like structures. **e(iv)**, Close up of the undecorated eV shown in e(i). **f,g(i-v),** Close ups of the **(f)** mouse-SFB and **(g)** rat-SFB features represented in a-d(iii) and a(iv) from the **(f)** mouse-SFB tomographic slices shown in a-d(i) and **(g)** from rat-SFB tomograms. **f,g(vi)**, Close up of decorated eVs from **f(vi)** mouse- SFB (EMD-52671) and **g(vi)** rat-SFB IOs (EMD-52689). The **g(i)** flagella and **g(iv)** filaments were isolated from the tomogram EMD-52689. The **g(ii)** intracellular vesicles, **g(iii)** chemosensory array and **g(v)** plate-like structures were isolated from tomograms EMD-52684, EMD-52690 and EMD-52691, respectively. eVs: extracellular vesicles; iVs: intracellular vesicles. **Scale bars:** a-e(i,ii): 50 nm; a-e(iii), a,c-e(iv), f/g: 20 nm.

Two types of intracellular filament-like structures were also identified in both mouse- and rat-SFB IOs **(Supplementary Table 1)**. One type consisted of multiple filaments clustered together with a thickness of approximately 28 ± 12 nm (n = 18) which were challenging to track (**Fig. 2c,f/g (iv))**, while the other filaments appeared to form plate-like structures with a thickness of approximately 63 ± 20 nm (n = 6) (**Fig. 2d,f /g(v))**. The width of individual clustered filaments and plate-like structures was significantly different (*p*<0.0001). Clustered filaments had a width of approximately 8/9 nm (mouse-SFB, 8 ± 1 nm, n = 18; rat-SFB, 9 ± 1 nm, n = 9), while the width of plate-like structures was 4/6 nm (mouse-SFB, 4 ± 1 nm, n = 6; rat-SFB, 6 ± 1 nm, n = 4) (**Fig. 2c/d , Supplementary** Fig. 2a**(ii-iii), b-d(iii-iv) and Supplementary Table 2)**. Clustered filaments and plate-like structures were at least 50 and 100 nm in length, respectively. Due to the technical limitations stemming from sample thickness and from the limited field of view, their full length could not always be determined.

Clustered filaments were found both in the centre of the cell body or parallel to the membrane in the concave side of IOs along the curvature of the cell **(Supplementary** Fig. 2a-b**(iii))**. These structures resembled the cytoskeletal filaments formed in *C. crescentus* by the enzyme CTP synthase^33^. In *C. crescentus*, these filaments are ∼500 nm in length, play a role in the regulation of bacterial curvature and, in initial developmental stages, they are found in the centre of the cell body but their position changes to the cell periphery parallel to the inner membrane in later developmental stages^33^. The presence of a CTP synthase-encoding gene in the genome of both mouse- and rat-SFB supports the hypothesis that the observed filaments may be of similar composition as the cytoskeleton filaments of *C. crescentus*. Conversely, plate-like structures were less common **(Supplementary Table 1)** and were found only in the centre of the IOs cell body either isolated or in groups of 2-3 plate-like structures, showing a maximum of 200 nm in length when their full length could be measured **(Supplementary** Fig. 2a**(ii),b(iv))**. To our knowledge, these plate-like structures do not resemble any previously characterized structures.

Lastly, two types of extracellular vesicles (eVs) were identified: undecorated (**Fig. 2e and Supplementary** Fig. 2e) and decorated (**Fig. 2f /g(vi) and Supplementary** Fig. 2a**(iv),b(iii),c/d(v))**. Similar undecorated eVs have been identified for the monoderm gut commensal *Bifidobacterium longum*^34^. In SFB, eVs were found in close proximity (**Fig. 2e and Supplementary** Fig. 2a**(iv),b(iii))** and in direct contact with the SFB tip **(Supplementary** Fig. 2e**(iii))**. Additionally, both undecorated and decorated vesicles were identified in SFB that possess both intact and ‘broken tip’ phenotypes (**Fig. 2e , Supplementary** Fig. 2a**(iv), b(iii) and Supplementary Table 1)**. Overall, the average diameter of eVs was 92 ± 44 nm (n = 15) for mouse-SFB and 103 ± 54 nm (n = 26) for rat-SFB **(Supplementary Table 2)**, placing them within the range of Gram-positive eVs^35^ and supporting their SFB-derived origin. The formation of eVs when SFB are in their native environment is supported by the identification of vesicles surrounding SFB IOs found attached to the epithelium of cows^36^. Overall, whole-cell cryo-ET thereby enabled the identification of expected, predicted, as well as unanticipated cellular features of mouse- and rat-SFB.

### The SFB tip undergoes a morphological developmental transition

We next refocused on the surface characterization of the SFB tip (**Fig. 3a-o , Supplementary movies 6-9)** as this structure mediates adhesion to the host epithelium and is therefore the primary location for potential attachment-related structures. The SFB imaged were grouped into stages based on the surface structures identified at the tip. In mouse-SFB IOs of less than 3.2 µm in length (n = 55), 24% were fully covered by an S-layer (stage 1 IOs, **Fig. 3a /f**). In the remaining IOs below 3.2 µm, the tip showed varying degrees of disordered hair-like structures (disHLS). DisHLS were found only at the tip edge in 20% of IOs (stage 2 IOs, **Fig. 3b/f**), while they were consistently present further down the tip in 18% of IOs (stage 3 IOs, **Fig. 3c/f , Supplementary** Fig. 4a). In 20% of the IOs below 3.2 µm, discontinuities were observed in the bacterial S-layer without the presence of disHLS, leading to the ’broken tip’ phenotype that prevented us from identifying the tip stage (Undefined: UD, **Fig. 3f , Supplementary** Fig. 5a). This ‘broken’ phenotype was only found at the tip, suggesting increased fragility of this region, and was predominantly found in IOs (**Supplementary** Fig. 5b**/c**), potentially due to the S-layer to disHLS transition.

**Fig. 3.**
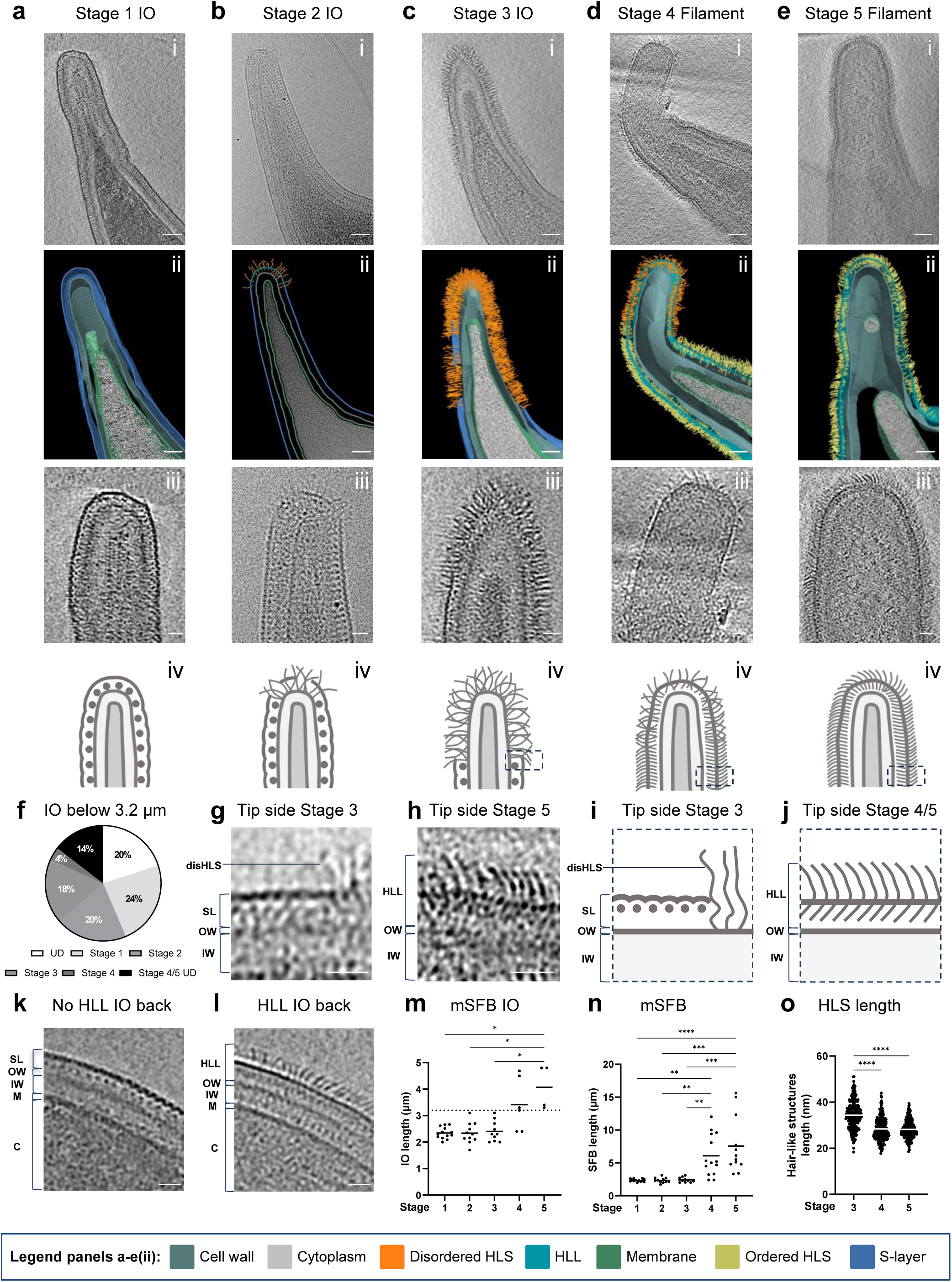
Identification of a morphological developmental transition at the tip of mouse-SFB. **a-e**, Morphological stages identified at the tip of mouse-SFB. **a,c,d,e(i),** Representative tomographic slices from reconstructed tomograms showing SFB tip of stages 1 (EMD-52685, Supplementary movie 6), 3 (EMD-52676, Supplementary movie 7), 4 (EMD-52677, Supplementary movie 8) and 5 (EMD-52678, Supplementary movie 9). **b(i),** Representative projection image of stage 2. SFB selected as representative of stages 1-5 are 2.2, 2.3, 3.1, 12.0 and 15.6 µm in length, respectively. **a-e(ii),** Segmentation rendering of mouse-SFB tip from stages 1-5. Colour legend is displayed at the bottom of the figure. Colouring of the hair-like structures (HLS) denotes their qualitative appearance. **a-e(iii),** Close ups of the mouse-SFB tip from stages 1-5. **a-e(iv),** Schematic representation of the mouse-SFB tip from stages 1-5. The region of the SFB tip represented in i,j is delimited by a dashed line**. f,** Pie chart showing the percentage of IOs below 3.2 µm in length identified in each tip stage shown in panels a-d. IOs for which the tip stage could not be identified (UD: unidentified) and for which the distinction between tip stages 4 and 5 could not be made are also shown. **g-h,** Close ups of side of the mouse-SFB tip from: **(g),** stage 3 and **(h),** stage 5. **i,j,** Schematic representation of the layers present at the side of mouse-SFB tip assigned to: **(i)** stage 3 and **(j)** stage 4/5. **k,l,** Representative tomographic slices of the back of a mouse-SFB IO in which an hair-like layer (HLL) was **(k)** absent (EMD-52696) and **(l)** present (EMD-52679). The length of the selected IOs was 2.4 and 2.3 µm, respectively. DisHLS: Disordered hair-like structures, HLL: Hair-like layer, SL: S-layer, disHLS: disordered hair-like structures, OW: outer wall zone, IW: inner wall zone, M: membrane, C: cytoplasm. **m,n,** Length of: **(m)** mouse-SFB IOs and **(n)** total mouse-SFB assigned to each stage. Sixty SFB, including 44 IOs, from 7 independent experiments were included in the analysis. A dashed line separates small (< 3.2 µm) and outgrowing IOs (> 3.2 µm) (m). **o**, Length of hair-like structures from the tip of SFB assigned to stages 3-5. Hair-like structures length was measured from tomograms of 5 bacteria per stage from 2 independent experiments. For each bacterium, 10 structures were measured from 5 tomographic slices. For m-o, individual measurements and the corresponding mean are shown. The statistical significance was assessed using a Kruskal–Wallis test followed by a Dunn’s test correction for multiple comparisons (**m:** Stage 1 vs 5, *p*= 0.0158; Stage 2 vs 5, *p*= 0.0160; Stage 3 vs 5, *p*= 0.0288; **n:** Stage 1 vs 4, *p*=0.0010; Stage 2 vs 4, *p*= 0.0016; Stage 3 vs 4, *p*= 0.0040; Stage 1 vs 5, *p*< 0.0001; Stage 2 vs 5, *p*= 0.0002; Stage 3 vs 5, *p*= 0.0005; **o**: *p<* 0.0001). **Scale bars:** a-e(i,ii): 50 nm; a-e(iii), g/h, k/l: 20 nm.

The distinctly repetitive S-layer subunits could no longer be seen in the regions occupied by disHLS (**Fig. 3c and Fig.S5a)**. However, the S-layer remained present in stages 2 and 3 IOs further down the tip towards the cell body (**Fig. 3b (iii),g/i)** as well as in the back, where disHLS were never observed (**Fig. 3k , Supplementary** Fig. 6a-b). Through tracking of the disHLS in tomograms, these structures were found to be in contact, or close proximity, to the electron dense line corresponding to the outermost region of the cell wall (**Fig. 3c/g/i , Supplementary** Fig. 4a) and to be 34 ± 6 nm in length (n = 250) (**Figure 3o**) and 1.3 ± 0.4 nm in width (n = 250). Together, these results reveal a structural transition from an S-layer to disHLS specifically at the tip of IOs.

As IOs transitioned to filaments, the bacterial surface became covered by a morphologically distinct outer layer, here designated as hair-like layer (HLL). This outer layer was found in 18% of IOs below 3.2 µm (**Fig. 3f**) and in 100% of bacteria above this length **(Supplementary** Fig. 5b**/c)**. It is composed of an array of ordered hair-like structures (ordHLS) at the SFB tip end (stages 4 and 5, **Fig. 3 d-e , Supplementary** Fig. 4b-c**(i-ii)**), tip side (**Fig. 3h/j , Supplementary** Fig. 4b**- c(iii))** and back (**Fig. 3l , Supplementary** Fig. 6c-d). In stage 4 SFB, the end of the tip possessed some disHLS, which, however, were significantly smaller (*p* < 0.0001) than those of stage 3 (28 ± 5 nm, n = 250, **Fig. 3o**). The cell body of stage 4 SFB was fully covered by the HLL. We only identified one IO (3.3 µm in length) out of the 14 stage 4 SFB imaged (7%) that showed all three features: HLL, disHLS and regions without either disHLS or HLL **(Supplementary** Fig. 7). This IO may therefore represent a rare transitional stage between stages 3 and 4.

In stage 5 SFB, the disHLS were no longer present (**Fig. 3e/h/j , Supplementary Fig.6d)** at the tip. Conservation of the HLL at the SFB back was confirmed in stages 4 and 5 SFB of variable lengths **(Supplementary Table 1)**, including filaments of over 30 µm **(Supplementary** Fig. 6d). As stages 4 and 5 SFB were significantly longer than the SFB stages 1 to 3 (*p* values from *p* = 0.0040 to *p* < 0.0001, **Fig. 3m-n**), stages 4 and 5 correspond to more advanced growth stages.

The filament-associated HLL was located above the cell wall and could be divided into an inner and outer part, separated by an electron dense layer. This electron dense layer was in a similar location to where the crescent-like elements of the IO-associated S-layer were in stage 1-3 SFB (**Fig. 3g-j**). Through manual tracing, individual electron dense hair-like subunits, comprising inner and outer parts of the HLL, were found to be 1.4 ± 0.4 nm (n = 250) in width and, at 28 ± 4 nm (n = 250) in length, to be significantly shorter than the disHLS of stage 3 (*p* < 0.0001, **Figure 3o**). Together, these results reveal a morphological transition of the SFB surface from an S-layer, surrounding small IOs, to a filament-associated hair-like layer characterized by ordered elongated subunits, with the transient appearance of longer disordered hair-like structures occurring only at the SFB tip. The disHLS are therefore in a location and present at a developmental stage consistent with playing a role in SFB attachment to the host.

### The developmental transition is conserved in SFB from different hosts

We next aimed to determine whether the morphological transition of the SFB outer layer was conserved across SFB from a different host by characterizing the outer layer of rat-SFB during the outgrowth of IOs into filaments. As observed for the mouse-SFB tip, features of the early development stages (stages 1 and 3), including the S-layer and the replacement of this layer by disHLS at the tip, were only found in IOs (**Fig. 4a-b**). In addition, the IO-associated S-layer and the absence of disHLS were conserved in rat-SFB at the back of stages 1 and 3 IOs (**Fig 4e , Supplementary** Fig. 6e**/f)**. Similarly, the later developmental stages (stages 4 and 5), characteristic also of longer rat-SFB (*p* values from *p* = 0.0023 to *p* < 0.0001, **Fig. 4g and Supplementary** Fig. 5d), possessed the filament-associated HLL at the tip (**Fig. 4c-d**) and at the back (**Fig. 4f and Supplementary** Fig. 6g). However, some differences were noted. There was a significant increase (*p* < 0.0001) in the proportion of IOs (74% of rat-SFB IOs vs 14% of mouse- SFB IOs) that could not be assigned to a tip stage (UD, **Supplementary** Fig. 5e-g). Of these, 31% also showed considerable S-layer discontinuities at the back (**Supplementary** Fig. 6h **and Supplementary Table 1**). As the purification conditions were the same for both mouse and rat- SFB, this may be due to an increased fragility of the rat-SFB S-layer. Despite these differences, our results confirm the conservation of the main developmental transitions at the tip and the back of rat-SFB including the S-layer, the disHLS at the tip, and the HLL surrounding SFB as they grow out into filaments.

**Fig. 4.**
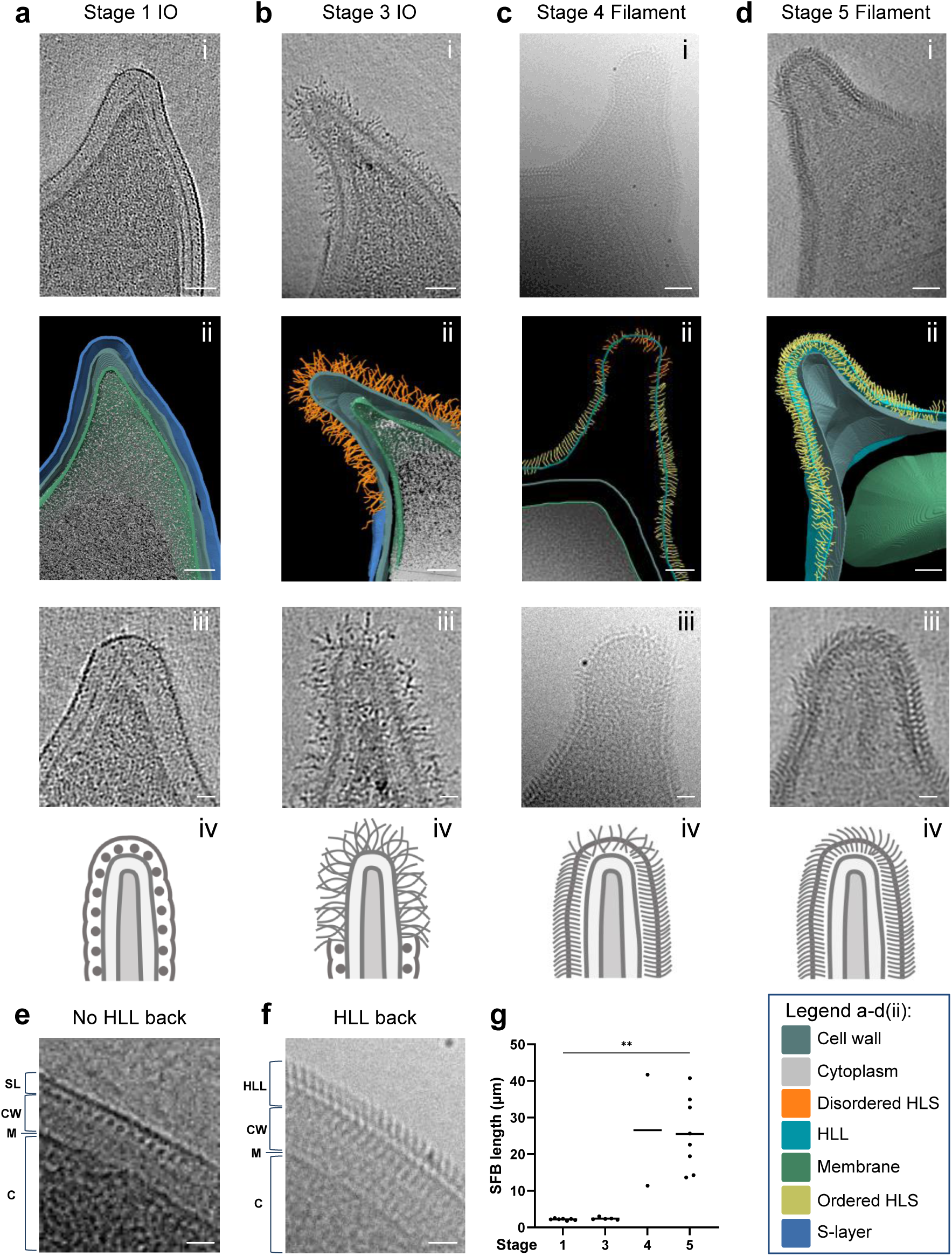
Conservation of the morphological developmental transition at the tip in rat-SFB. **a-d**, Rat-SFB morphological stages equivalent to those identified at the tip of mouse-SFB. **a,b,d(i),** Representative tomographic slices from reconstructed tomograms showing SFB tip of stages 1 (EMD-52695), 3 (EMD-52688) and 5 (EMD-52699). **c(i),** Representative projection images of the SFB tip of stage 4. The selected SFB from stages 1,3,4 and 5 have a length of 2.2, 3.0, 41.7 and 34.9 µm, respectively. **a-d(ii),** Segmentation rendering of rat-SFB tip from stages 1,3,4 and 5. Colour legend is displayed at the bottom of the figure. Colouring of the hair-like structures (HLS) denotes their qualitative appearance. **a-d(iii)**, Close ups of the SFB tip. **a-d(iv),** Schematics of the SFB tip. **e/f,** Representative projection images of the back of rat- SFB in which an hair-like layer (HLL) was **(e)** absent and **(f)** present. SL: S-layer, HLL: hair-like layer, HLS: hair-like structures, OW: outer wall zone, IW: inner wall zone, M: membrane, C: cytoplasm. **g**, Length of SFB assigned to each tip stage. 32 SFB from 5 independent experiments were included in the analysis. Individual measurements were plotted, the mean is shown and the statistical significance was assessed using the Kruskal–Wallis test followed by a Dunn’s test correction for multiple comparisons (Stage 1 vs 5, *p*=0.0023). **Scale bars:** a-d(i,ii): 50 nm; a-d(iii), e/f: 20 nm.

### The filament-associated hair-like layer is composed of repetitive subunits

We next aimed to obtain a more precise characterization of the hair-like layer by focussing on the inner (I-HLL) and outer (O-HLL) parts of this filament-associated layer. The I-HLL from stage 4-5 SFB was in a similar location to the IO-associated S-layer globular subunits from stage 1-3 IOs (**Fig. 3g-j**, **Fig. 5a/b**), whereas the O-HLL extended beyond the IO-associated crescent-like elements of the S-layer (**Fig. 5c-e**). While the outer and inner subunits appeared to be aligned, and may therefore be connected, an electron dense layer separated these subunits.

**Fig. 5.**
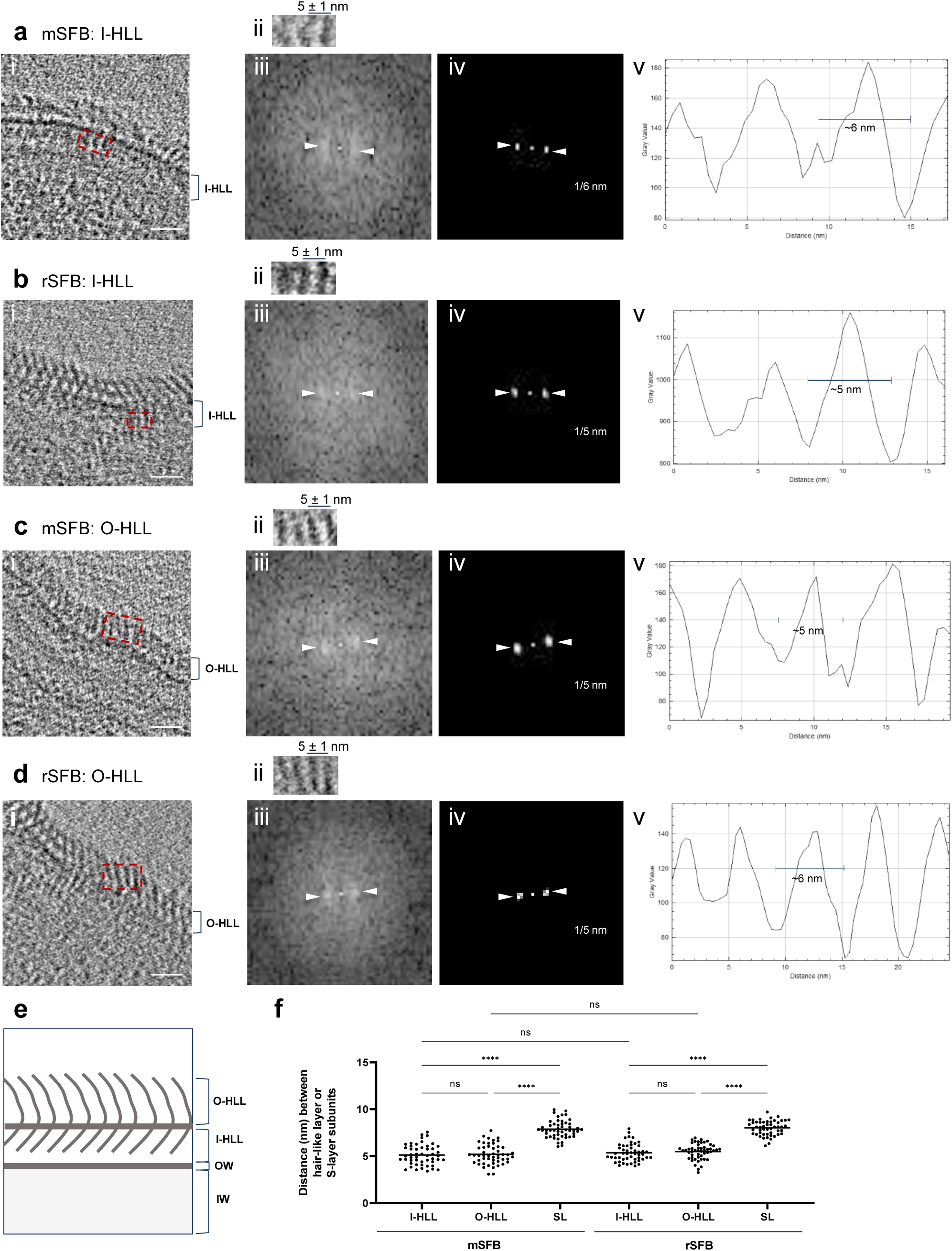
Characterization of the filament-associated hair-like layer. **a-d(i),** Close ups of the tip side of: **(a,c(i))** mouse-SFB and **(b,d(i))** rat-SFB from reconstructed tomograms EMD-52677 and EMD-52699, respectively. The regions used for hair- like layer (HLL) analysis are delimitated by red rectangles. **a-d(ii),** Regions containing: **a,b(ii)** the inner part of the hair-like layer (I-HLL) and **c,d(ii)** the outer part of the hair-like layer (O-HLL) shown in a-d(iii). The average spacing and standard deviation between HLL subunits is shown. **a-d(iii),** Fast Fourier transform, **a-d(iv),** Maximum intensity peaks (indicated by white arrows) and their corresponding frequency, and **a-d(v),** Plot profiles are shown for: **(a(i))** mouse-SFB I-HLL, **(b(i))** rat- SFB I-HLL, **(c(i))** mouse-SFB O-HLL, **(d(i))** rat-SFB O-HLL. **e,** Schematic representation of inner and outer parts of the HLL. OW: outer wall zone, IW: inner wall zone. **f,** Distance between HLL and S-layer (SL) subunits in mouse-SFB and rat-SFB. Ten measurements were performed for each of the 5 SFB from 2 independent experiments. The mean is shown and the statistical significance was assessed using the One-way ANOVA (****: *p*<0.0001, ns: not significant). The most important comparisons are shown. Mouse-SFB: mSFB, Rat-SFB: rSFB. CW: cell wall. **Scale bars:** a-d(i): 20nm.

The distance between the individual subunits of the inner and outer HLL was estimated by Fourier spectra analysis and plot profile tracing on the side view of tomograms of the SFB filament tips and combined with manual measurements (**Fig. 5a-d**). The subunits in both the inner and outer layer were found to be repetitive and approximately 5 nm apart (n = 50), both in mouse and rat- SFB (**Fig. 5a-d,f**). This distance between subunits was significantly smaller (*p* < 0.0001) than the approximately 8 nm separating the globular subunits of the S-layer (**Fig. 5f**). These results support the replacement of the IO-associated S-layer with a morphologically distinct hair-like layer composed of more tightly arranged repetitive units as SFB transition from IOs to filaments.

### Th17Ag is accessible at the tip of SFB filaments

The hair-like layer described is, to our knowledge, morphologically distinct from any surface structure previously identified by cryo-EM/ET and its molecular composition remains unknown. The major Th17 antigen (Th17Ag)^7^ was previously found to surround attached bacteria of mouse- SFB-NYU^18^, making it a potential candidate for a component of the hair-like structures constituting the HLL. To assess the localization of this SFB factor in more detail, we recombinantly expressed and purified the mouse-SFB-NL homolog (AID45212) **(Supplementary** Fig. 9a-b). Th17Ag was localized at the mouse-SFB-NL surface both by immunogold labelling followed by cryo-EM, and by immunofluorescence followed by confocal microscopy (**Fig. 6a-g , Supplementary** Fig. 10a**/b, Supplementary** Fig. 11a-g).

**Fig. 6.**
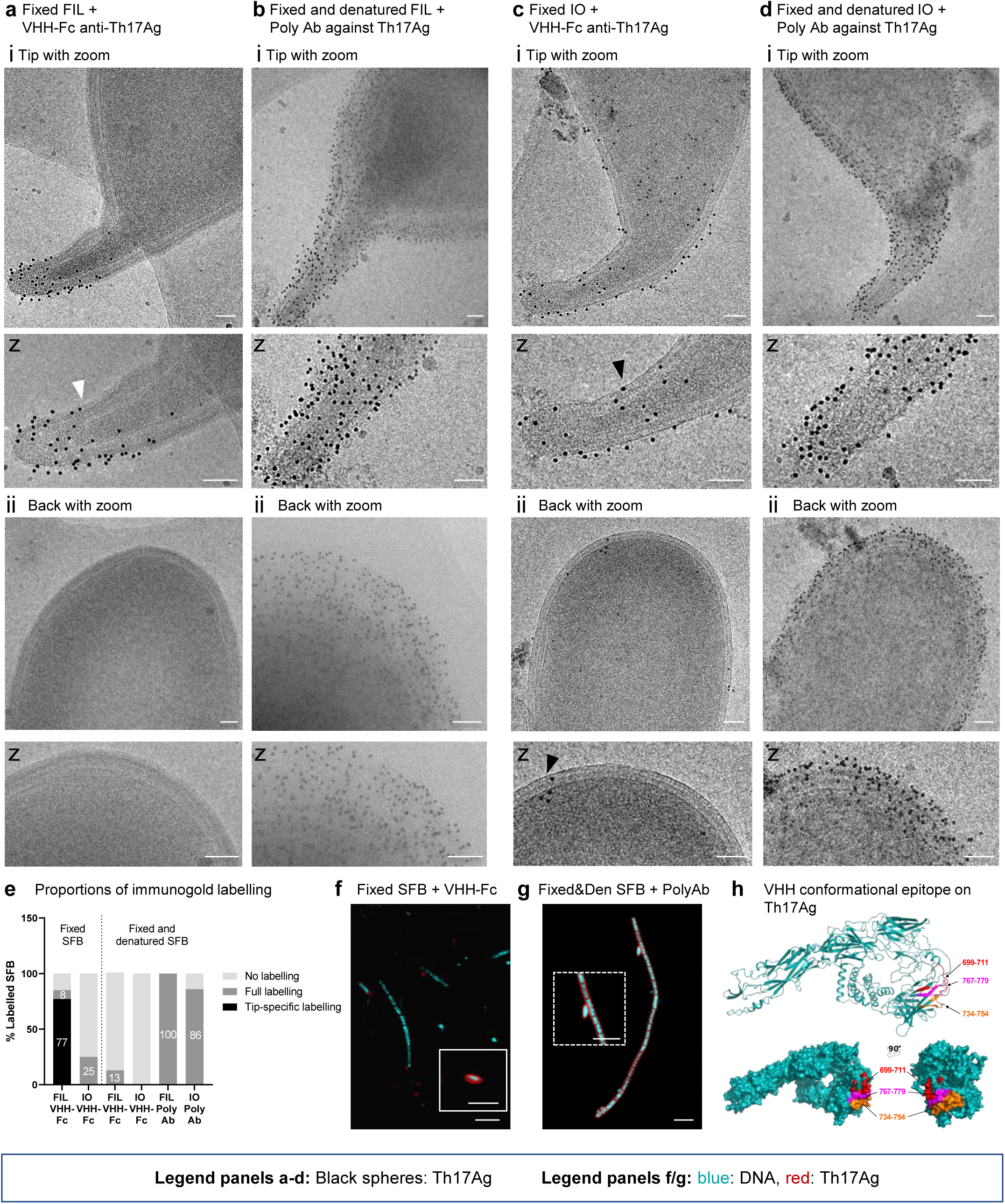
Localization of Th17Ag at the SFB surface. **a-d,** Projection images from purified SFB **(a,b)** filaments (FIL) and **(c,d)** IOs stained using immunogold labelling with gold-conjugated Protein A. Fixed SFB **(a)** filaments and **(c)** IOs were labelled with a VHH-Fc anti-Th17Ag. Fixed and denatured SFB **(b)** filaments and **(d)** IOs were labelled with a rabbit polyclonal antibody against Th17Ag. Imaging was performed at the SFB **(a-d(i))** tip and **(a-d(ii))** back. Close ups (z) of the SFB tip and back are shown under each panel. Examples of labelled regions both with and without visible hair-like structures are shown by white and black arrows, respectively. Projection images were acquired with a Tecnai F20 electron microscope equipped with a Falcon 2 camera. **e,** Assessment of immunogold labelling for SFB imaged in the conditions described for panels a/d and after labelling of fixed and denatured SFB with a VHH-Fc anti-Th17Ag. The percentage of SFB labelled and labelled specifically at the tip are indicated on the corresponding bar. SFB were considered labelled if co- localization with at least 20 gold particles (black spheres) was observed, or at least 10 gold particles if labelling was restricted to the SFB tip. Between 7 and 20 SFB were imaged for each condition**. f,g,** Immunofluorescence images of **(f)** fixed SFB incubated with a biotinylated VHH-Fc anti-Th17Ag and Streptavidin-Alexa568 and **(g)** fixed and denatured SFB incubated with a rabbit polyclonal antibody (Ab) against Th17Ag and a secondary antibody anti-rabbit conjugated with Alexa 568. All SFB were additionally labelled with DAPI. An extra image of a labelled IO was included as an insert delimited by a white line (f). A close up of the IO shown in panel g was included as an insert delimited by a dashed white line. For all immunogold and immunofluorescence experiments, labelling was performed in two distinct days using two biological replicates for each experiment. **h,** Localization of the nanobody binding epitope on the structure of Th17Ag predicted by AlphaFold^57^. Structure prediction with surface representations of Th17Ag highlighting the protein regions (in red, orange and magenta) that form the conformational epitope recognized by the nanobody. Epitope location was determined by Hydrogen/Deuterium eXchange-Mass Spectrometry (HDX-MS). **Scale bars**: a-d: 100 nm; f/g: 5 µm.

Labelling with a nanobody (VHH) targeting Th17Ag **(Supplementary** Fig. 9c), fused to the Fc region of human IgG1 (VHH-Fc), revealed distinct labelling patterns in SFB filaments and IOs. For filaments, Th17Ag labeling was seen in 85% of the filaments imaged (n = 13) and was strikingly restricted to the filament tip in 91% (n = 10) of those (**Fig. 6a/e**). For IOs, Th17Ag labelling was observed only in 25% of the IOs imaged (n = 20) (**Fig. 6e**). This labelling occurred at the tip, but also on the remaining cell body, including the back (**Fig. 6c**). In agreement, IOs showed labelling with the VHH-Fc targeting Th17Ag using immunofluorescence, while filaments did not (**Fig. 6f**).

The tip-restricted labelling of SFB filaments **(Supplementary** Fig. 11a**/e)** contrasts with earlier reports showing labelling of Th17Ag surrounding SFB in histological sections^18^. As histological processing involves sample permeabilization, we treated PFA-fixed SFB with 0.5% triton X-100 at 95°C for 5 minutes, which disrupts the outermost layer without compromising cell shape integrity. This treatment largely prevented all labelling with the VHH-Fc (**Fig. 6e , Supplementary** Fig. 11b), but using the rabbit polyclonal antibody raised against Th17Ag^7^, recapitulated the published findings of intense labelling surrounding 100% of the filaments (n = 7), including the tip structure (**Fig. 6b/e/g**). In addition, the rabbit polyclonal antibodies showed strong staining of the bacterial surface in 86% of IOs imaged (n = 7) (**Fig. 6d/e**). Under non-denaturing conditions, the rabbit polyclonal antibody displayed similar staining patterns as the VHH-Fc **(Supplementary** Fig. 11a**/c/e/f)**. Epitope mapping of the VHH, following assessment of binding affinity **(Supplementary** Fig. 12a**, Supplementary Table 3)**, identified three target regions of Th17Ag towards the protein C-terminus (**Fig. 6h**). The nearly nonexistent immunolabelling with the VHH-Fc under denaturing conditions (**Fig. 6e , Supplementary** Fig. 11b**/g)**, is therefore likely due to the disruption of the conformational epitope recognized by VHH (**Fig. 6h , Supplementary** Fig. 12b**/c)**. These results identify Th17Ag as an abundant component of the SFB surface for both IOs and filaments whose accessibility is largely restricted to the tip in filaments and moderately accessible over the full surface in a subset of IOs. Th17Ag, which contains a domain of unknown function also found in SFB proteins annotated as peptidoglycan hydrolases^19–22^, is therefore likely a cell wall-anchored protein whose accessibility in filaments is prevented by the HLL. For IOs, as only a small subset was labelled, we hypothesize that the increased accessibility may be related to the transition of the S-layer to the HLL, rather than a general greater permeability of the S-layer. As Th17Ag was detected on the entire surface of IOs, including at the back where disHLS were not present (**Fig. 6c and Supplementary** Fig. 11c), it is unlikely that Th17Ag is a component of the disHLS. Overall, our results point towards an increased exposure of the cell wall specifically at the SFB tip.

### SFB growth in a heterologous host affects adhesive tip morphology

We next wanted to evaluate the identified morphological stages of the tip of mouse-SFB in a context where adhesion was not supported. Attachment, but not growth, of SFB has been shown to be host-specific for rat-SFB under monocolonization conditions in mice^14^. Similarly, mouse-SFB did not attach to the ileal epithelium in rats while mouse-SFB could still grow robustly in the rat intestinal lumen **(Supplementary** Fig. 8). When mouse-SFB were grown in rats, all five developmental stages were identified, as well as the association of stages 4 and 5 with the outgrowth into filaments (**Fig. 7a-f**). However, we observed a decrease in tip length for mouse- SFB, both in IOs (575.6 ± 146.8 nm in mice, n = 24, vs 467.4 ± 95.0 nm in rats, n = 36, *p* = 0.0010) and filaments (625.0 ± 138.4 nm in mice, n = 39, vs 439.2 ± 193.4 nm in rats, n = 10, *p* = 0.0087) (**Fig. 7g/h,k/l**) when SFB of equivalent lengths were compared (**Fig. 7i/j**). Additionally, stage 2 IOs were present at a significantly higher proportion when mouse-SFB were grown in rats compared to when grown in mice (61% vs 32%, *p* = 0.0147) (**Fig. 7m-n**). These findings support a role of attachment in the elongation of the tip and in the potential transition of SFB from stage 2 to stage 3, the stage where the disHLS are most prominent.

**Fig. 7.**
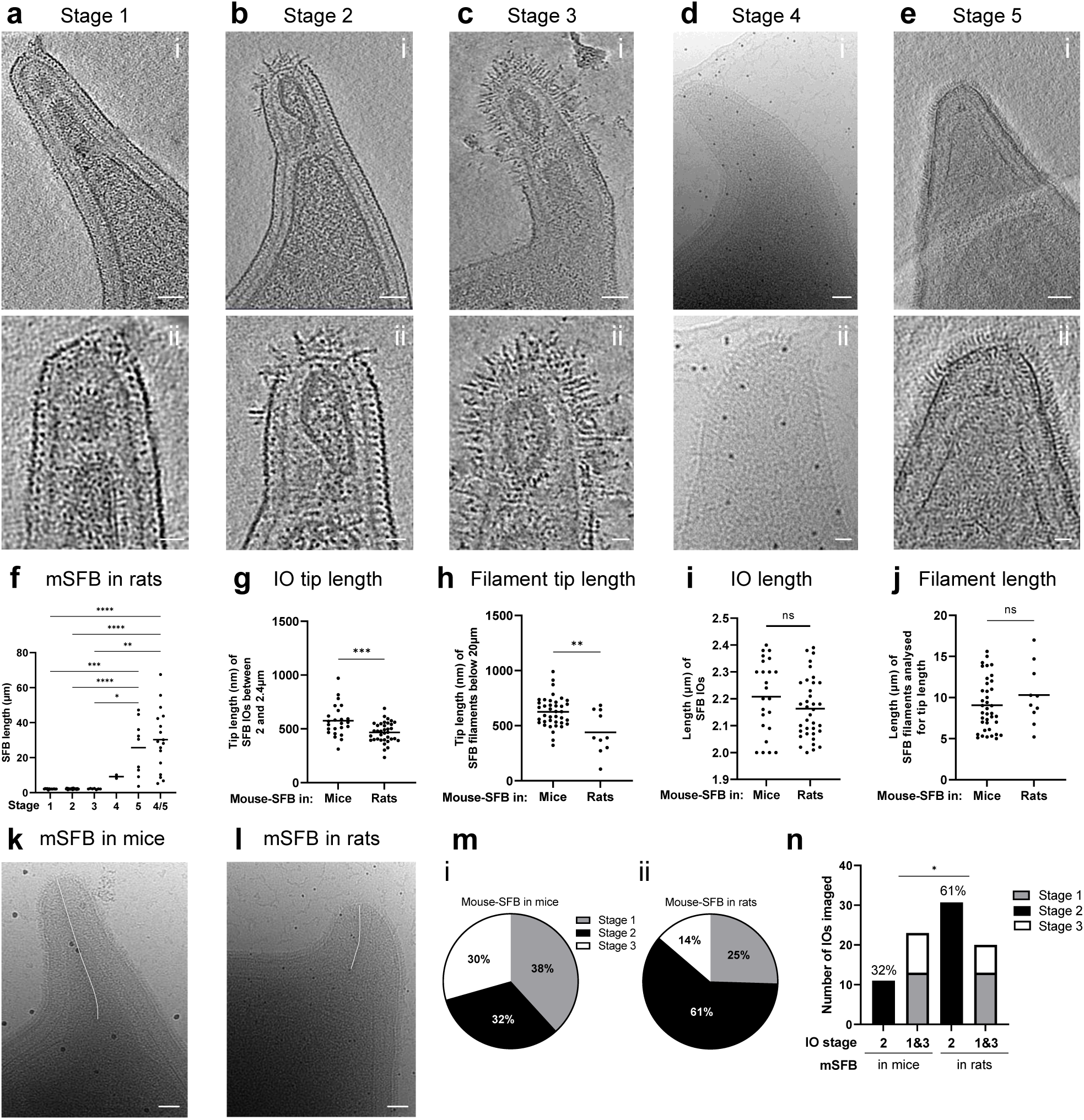
Effect of SFB growth in a heterologous host on tip morphology. **a-e,** Mouse-SFB morphological stages, when SFB was grown in rats, equivalent to those identified at the tip of mouse-SFB colonizing mice. **a-c,e(i),** Representative tomographic slices from reconstructed tomograms showing SFB tip of stages 1 (EMD-52700), 2 (EMD-52701), 3 (EMD- 52702) and 5 (EMD-52703). **d(i),** representative projection image showing SFB tip of stage 4. The selected representative SFB from stages 1-5 have a length of 2.1, 2.3, 1.6, 8.2 and 3.7 µm. **a-e(ii),** Close ups of the SFB tip shown in a-e(i). **f,** Length of mouse-SFB grown in rats assigned to each tip stage, including filaments where no distinction could be made between stages 4 and 5. A total of 79 SFB from 4 independent experiments was included in the analysis. Individual measurements and the corresponding mean are shown. The statistical significance was assessed using the Kruskal-Wallis test followed by a Dunn’s test correction for multiple comparisons (Stage 1 vs 5, *p*=0.0006; Stage 2 vs 5, *p*<0.0001; Stage 3 vs 5. *p*=0.0181, Stage 1 or 2 vs Stage 4/5, *p*<0.0001; Stage 3 vs 4/5, *p*=0.0027). **g,h,** Length of the tip of mouse-SFB **(g)** IOs and **(h)** filaments analysed. **i,j**, Length of corresponding SFB **(i)** IOs and **(j)** filaments used for tip length analysis of SFB grown in mice and rats. IOs between 2 and 2.4 µm of length were included in the analysis (27 grown in mice and 38 grown in rats). Individual measurements and the corresponding mean are shown. **h,** Length of mouse-SFB filaments tip and **j,** length of corresponding SFB filaments used for tip length analysis of SFB grown in mice and rats. Filaments below 20 µm length were included in the analysis (39 from mice and 10 from rats). Individual measurements and the corresponding mean are shown. The statistical significance was assessed using a t-test for panels g and i (g: *p*=0.0004, i: ns: not significant) and using a Mann–Whitney U test for panels h and j (h: *p*=0.0087, j: ns: not significant). **k,l**, Representative projection images of mouse-SFB filaments grown in **(k)** mice and **(l)** rats. The tip length measurement is shown by a white line. **m,** Proportions of IOs assigned to stages 1-3 when grown **(m(i))** in mice and **(m(ii))** in rats. **n,** Comparison of the proportions of stage 2 IOs of mouse-SFB grown in mice and in rats. A total of 33 mouse-SFB IOs grown in mice and 51 mouse-SFB IOs grown in rats was used for the analysis. The statistical significance was assessed using the Fisher’s exact test (Stage 2 in mice vs in rats, *p*=0.0147. **i,** Length of mouse-SFB filaments tip and **j,** length of corresponding SFB filaments used for tip length analysis of SFB grown in mice and rats. All filaments included in the analysis were below 20 µm in length (39 grown in mice and 10 grown in rats). Individual measurements and the corresponding mean are shown. The statistical significance was assessed using a Mann–Whitney U test (i: *p*=0.0087, j: ns: not significant). **Scale bars:** a-e(i), k/l: 50 nm; a-e(ii): 20 nm.

## Discussion

In this study, we characterized the SFB surface by cryo-EM and cryo-ET. We describe two morphologically distinct outer layers that are conserved in SFB from different rodent hosts. The first possesses the characteristics of a bacterial S-layer with morphological similarity to the S-layer of *C. crescentus* and related Clostridia^27–29^. The second, here designated as hair-like layer (HLL), is composed of regularly spaced subunits that resemble hair-like structures but, to our knowledge, possesses no clear resemblance to any other outermost layer previously imaged.

The S-layer is only seen in small unicellular SFB, characteristic of early developmental stages, while the HLL is present in longer IOs and in all SFB filaments, characteristic of later developmental stages. The globular subunits of the S-layer can no longer be seen when the HLL is present and the subunits spacing is significantly different between the S-layer and the HLL.

Our results suggest that the transition to the HLL occurs rapidly, as it was challenging to identify transitional stages in which both layers were concomitantly present at the SFB surface. The HLL may itself constitute an S-layer since it is not only repetitive but also appears to prevent the access of antibodies to proteins like the abundant Th17Ag that is part of the bacterial cell wall. The potential replacement of one S-layer with another is reminiscent of the transition from an exponential phase-associated (Sap) S-layer to a stationary phase-associated (EA1) S-layer previously described for *Bacillus anthracis*^37^.

S-layers can play a role in host adhesion in bacteria with similar cell ends^30^. However, since in SFB it is the polar tip that mediates epithelial cell attachment, we hypothesize that both the S-layer and the HLL are not involved in adhesion but mainly in bacterial protection and/or maintenance of cell wall integrity. Extracellular vesicles were observed near IOs showing S-layer discontinuities. This phenotype resembled the membrane blebbing seen in hypotonic conditions upon EA1 S- layer depolymerization^38^, supporting the hypothesis of a role for the IO-associated S-layer in cell wall integrity.

Additional surface structures identified here are the tip-specific disHLS. Since these structures were morphologically different from the subunits of the HLL (ordHLS) and only located at the bacterial tip, we hypothesize that these structures may also be functionally different. The disHLS found at the tip of IOs otherwise surrounded by an S-layer were longer and morphologically distinct compared to the ordHLS found at the tip and surrounding long IOs and filaments, suggesting that the disHLS and ordHLS may have a different molecular composition. For type IV pili of *Thermus thermophilus*, the morphological differences observed using cryo-EM were due to distinct protein compositions leading to functional differences^39^. Nevertheless, since no differences were found between the length of disHLS found in Stage 4 SFB and ordHLS, experimental evidence at the molecular level is needed to elucidate the observed variations in HLS length.

Since attachment is mediated by the SFB tip^17^, the tip-restricted location of disHLS suggests these structures may be involved in SFB attachment to host cells. Furthermore, their emergence prior to bacterial elongation and the increase in the proportion of IOs without fully developed disHLS in a heterologous host, where attachment is not supported, further support this hypothesis. Growth of mouse-SFB in rats also revealed that attachment is not required for the developmental transitions identified at the bacterial surface. This is consistent with the developmental transitions that occur during the life cycle of the environmental bacterium *C. crescentus,* which attaches to abiotic surfaces using a stalk. In this species, adhesion is not necessary but the transition from the motile to sessile stage is promoted by surface contact^40^. Notably, the presence of a higher proportion of IOs without fully developed disHLS suggests a delay in SFB development when attachment is not supported. These results point towards the influence of internal and external signals in the regulation of the disHLS formation.

The tip length of mouse-SFB was also reduced when colonizing rats. A longer tip may provide better anchoring to the epithelium and higher access to host-derived nutrients which may be critical for SFB development, given that these bacteria lack most genes involved in the biosynthesis of nucleotides, amino acids, vitamins and cofactors^19–22^. Attachment-related cues may therefore play a role in the regulation of tip elongation, similar to what has been observed for the stalk length in *C. crescentus*^41^. Nevertheless, the tip of rat-SFB was shorter than the tip of mouse-SFB when the characterization was performed in their respective hosts. As the genetic background of the host can influence the colonization of microbiota members^42^, we cannot exclude the possibility that it also influences the development of morphological characteristics associated with the establishment of a niche, such as tip elongation.

Regarding the molecular details of the SFB surface, immunogold labelling of Th17Ag revealed novel findings related to protein exposure at the SFB surface. Th17Ag was previously found to surround SFB filaments in intestinal sections and to be taken up from the filament tip through the formation of host endocytic vesicles^18^. We obtained similar labelling with permeabilization but without permeabilization this protein was surface-exposed only at the filament tip for the majority of the filaments imaged. Similar to filaments, strong labelling was observed in IOs under permeabilization conditions. However, in non-permeabilization conditions only approximately a quarter of the IOs imaged showed labelling and labelling was not restricted to a specific region of the bacterial surface. These results suggest that the HLL in SFB filaments prevents access to this potential cell wall-anchored protein, while the S-layer appears to generally have a similar function for IOs. We hypothesize that the increased surface exposure in a subset of IOs may be due to changes in cell wall accessibility during the S-layer to HLL transition. These findings provide further evidence of a developmental transition occurring at the SFB tip.

One of the main limitations of this study is the thickness of our samples, which limited the resolution and prevented a more detailed characterization of the intracellular features identified, such as filaments, plate-like structures, tracks and chemosensory arrays. Future studies using cryo-FIB^43^ could overcome this limitation. The lack of a fully conclusive characterization of the SFB S-layer symmetry can also be considered a limitation. However, due to the twisted lattice at the bacterial tip, in-depth *in situ* symmetry identification, as it was performed for the long stalk of *C. crescentus*^27^, may necessitate the extraction of S-layer proteins from IOs. Yet, given the challenges related to the study of SFB, including the use of monocolonized mice for bacterial propagation, a lack of availability of large quantities of IOs, and the lack of knowledge on the proteins that compose the IO-associated S-layer, obtaining S-layer sheets in solution will be challenging. Here, we provide the first identification of the SFB S-layer and novel observed structures, opening the way for further characterization.

Overall, this study identifies novel surface structures and other intracellular and extracellular features, providing critical information on the SFB life cycle. We describe developmental changes on the SFB surface associated with the transition from unicellular to filamentous stage and identify tip-restricted disordered hair-like structures potentially involved in host attachment. Furthermore, we show a restricted surface accessibility of Th17Ag specifically at the SFB filament tip, identifying the SFB tip as a particularly immunologically vulnerable site where cell wall components are accessible. Given that major immunostimulatory properties of SFB are dependent on adhesion to host cells and that surface-exposed supramolecular structures generally play critical roles in host-bacterial interactions, the future characterization, on a molecular level, of the novel surface structures identified here should provide further insights on how attachment is established.

## Data availability

Reconstructed tomograms used as representative have been deposited in the EMDB (EMD- 52655; EMD-52667 to EMD-52670; EMD-52671 to EMD-52680; EMD-52682 to EMD-52685; EMD-52687 to EMD-52703; EMD-52856. The EMDB accession numbers for reconstructed tomograms were indicated in the figures which include their corresponding tomographic slices. The mass spectrometry data have been deposited to the ProteomeXchange Consortium via the PRIDE^56^ partner repository with the dataset identifier PXD060041.

## Author contributions

ARC and PS were responsible for experimental design. SFB purification, immunofluorescence, immunogold labelling and confocal microscopy imaging by ARC. Cryo-EM/ET sample preparation by ARC, THT, GPA and ADG, and data acquisition by ARC, BW, GPA, GPA, ADG, and JMW Cryo-EM/ET data analysis by ARC, BW, AG, ASR, ADG, OM and PS. SFB protein expression and purification by ARC and AL. Germ free animal maintenance by MB and ARC. VHH production and selection by GA, PL and ARC. VHH epitope mapping by HDX-MS by SB. Funding acquisition, project conception and supervision by PS. Manuscript writing by ARC and PS.

## Acknowledgements

We are thankful for the technical support provided by the members of the Center for Animal Resources and Research of the Institut Pasteur (Martine Jacob, Eddie Maranghi, Thierry Angelique, Jérôme Toutain and Marvin Pery) and SFR Necker (Emilie Panafieu, Amaury Gensou and Cristian Dicu). We thank Dr. Sylvie Rabot (INRAE) for providing germ-free rats. We acknowledge Dr. Dan Littman (NYU) for generously sharing the anti-P3340 (Th17Ag) rabbit polyclonal antibody. We acknowledge the SFR Necker Histology Platform, especially Damien Conrozier, for the technical support provided in the preparation of rodent intestinal cryosections. We thank Dr. François Bontems for the helpful discussions. We thank Gaëlle Chauveau-Le Friec for her technical work. The lab of PS is supported by INSERM, CNRS, Université Paris Cité. A.R.C was supported by a PhD fellowship from BioSPC, Université Paris Cité. This work was funded through the ERC grant NICHEADAPT (866222), the Bill and Melinda Gates Grand Challenge grant (OPP1141322), and the Bettencourt Foundation Coups d’élan prize awarded to PS. The work conducted at University of Zurich was funded through the Swiss National Foundation grant (10000885) awarded to OM. Open Access enabled by the ERC CoG NICHEADAPT (866222). The authors thank the center of microscopy and image analysis (ZMB) at the University of Zurich. The HDX-MS platform was funded by the CACSICE Equipex ANR-11-EQPX-0008. We are also grateful for support for Ultrastructural BioImaging Core Facility equipment from the GIS-IBISA, the DIM One Health, the French Government Programme Investissements d’Avenir France BioImaging (FBI, N° ANR-10-INSB-04-01) and the French gouvernement (Agence Nationale de la Recherche) Investissement d’Avenir programme, Laboratoire d’Excellence “Integrative Biology of Emerging Infectious Diseases” (ANR-10-LABX-62-IBEID).

## Methods

### SFB monocolonization

Germ-free C57BL/6J mice were bred and housed either at the animal facility of Institut Pasteur (IP) (agreement number 75-15-01) or at the animal facility of SFR Necker (agreement number A75-15-34) and selected regardless of their sex, based on availability. Germ-free male F344 rats between 5 and 9 weeks-old were bought from the Plateforme Anaxem de l’Institut Micalis (INRAE, Jouy-en-Josas) and housed at the IP animal facility. Monocolonization with mouse-SFB-NL was established for 7-13-week-old germ-free mice. Monocolonization with mouse-SFB-NL or rat-SFB- Yit was established for 7-17-week-old germ-free rats. To achieve SFB monocolonization, all rodents were kept under axenic conditions and gavaged with previously frozen fecal pellets of rodents monocolonized with the SFB strain of interest resuspended in water. Monocolonization was verified by Gram staining (Sigma Aldrich) and culture-based techniques. All rodents were sacrificed between 7- and 18-days post gavage for further SFB purification.

### Ethics statement

Animal experiments were performed in accordance with French and European regulations on the protection of the animals used for scientific purposes (Directive 2010/63 of the European Parliament and French decree of February 1, 2013). Rodent (dap210054 and dap220096) and alpaca (2020-27412) experiments were approved by the IP ethical committee for animal experimentation (Comité d’Ethique en Expérimentation Animale, CETEA, registry number #89) and authorized by the Ministère de l’Enseignement Supérieur, de la Recherche et de l’Innovation.

### SFB purification from rodent intestinal content

SFB were purified from the intestinal content of monocolonized mice and rats as previously described^32^ with some modifications. Briefly, all liquid reagents were pre-equilibrated overnight in a hypoxic chamber at 2% oxygen. Intestines were dissected in hypoxic conditions and contents homogenized in 20 mL of PBS 1X by gentle pipetting. The intestinal suspension was layered in one volume of 50% nycodenz (Proteogenix) and spun down at 4,000 x *g* for 15 minutes in a swinging bucket at room temperature. The interphase was collected, homogenized, layered in ½ volume of 30% nycodenz and spun down in the conditions above. Pellets were discarded and the remaining sample was washed in PBS 1X and spun down at 100 x *g* for 5 minutes. Pellets were again discarded, and supernatants spun down at 4,000 x *g* for 20 minutes. When necessary, samples were passed through a 5 μm filter to obtain an IO-only fraction and concentrated at 4,000 x *g* for 30 minutes. Reverse filtration was used to recover the filament-enriched fraction. The pellet was resuspended in PBS 1X and the suspension layered on 1 volume of 15% nycodenz and spun at 4,000 x *g* for 15 minutes. The pellet was recovered, washed in PBS 1X and the suspension was spun down as described above. Pellets corresponding to SFB fractions were resuspended in approximately 200 µL of PBS 1X and diluted from 1:2 to 1:10 depending on the purified SFB density verified by Gram staining. SFB suspensions were kept at 4 °C for a maximum of 24 h prior to preparation of cryo-EM samples.

### SFB preparation for Cryo-electron microscopy and tomography

Gold particles of 5 or 10 nm (ProteinA-gold, Aurion) were added to SFB samples in a 1:20 or 1:40 dilution, respectively, and gently homogenized. 4 µL of this suspension were deposited onto Lacey S166-3 mesh 300 copper grids (Agar Scientific) or Quantifoil R 2/1 or R2/2 200 mesh copper grids previously glow-discharged at 2 mA for 1 minute using an ELMO glow discharge system (Cordouan Technologies). 2 µL of PBS 1X were deposited on the grid and back blotting was performed for 2 to 6 seconds using an automatic plunge freezer (EMGP, Leica). During blotting, the temperature and humidity of the chamber were set to 10 °C and 98%, respectively. The samples were then cryofixed at −180 °C in liquid ethane and stored in liquid nitrogen until imaging. Grids containing frozen-hydrated SFB were subsequently clipped into autogrids (Thermo Fisher Scientific: TFS) whenever subsequent imaging was performed using a 300 kV Titan Krios electron microscope.

### Cryo-electron microscopy and tomography

Initial screening of the samples was performed using a 200 kV Tecnai F20 (FEI company) equipped with a direct detector Falcon II (TFS) at the IP Ultrastructural BioImaging Platform. Projection images of the tip and back of SFB were acquired using the software EPU (TFS) at a pixel size of 2.01 or 4.14 Å and are listed in **Supplementary Table 1**. Representative images of the identified phenotypes (Supplementary Table 1) were acquired with a 300 kV TITAN Krios electron microscope equipped with a Falcon 4i direct electron detector and a Selectris X energy filter (10 eV slit width), here designated as TITAN Krios IP, unless otherwise stated. Imaging was performed at the IP NanoImaging Core facility. Projection images of the tip and back of SFB were acquired with -10 µm defocus at a calibrated pixel size of 1.59 or 2.01 Å. Data acquisition was controlled by TOMO5 (version 5.14 or 5.17, TFS). The magnification used to obtain a full SFB in the field of view to determine SFB length varied according to the bacterial size (from less than 2 to more than 65 µm in length). Projection images were used to classify tip stages and to measure SFB length, tip length and vesicle diameter.

To obtain a 3D view and enable the measurement of the SFB structures observed, tilt series were acquired mainly at the SFB tip at high magnifications. We used either the TITAN Krios IP or a Titan Krios microscope (FEI company) operated at 300kV equipped with a post-column energy filter (20 eV slit width, Gatan) and a K2 direct electron detector (Gatan) at the University of Zurich, here designated as TITAN Krios UZH. For the latter microscope, data acquisition was controlled by SerialEM 3.8^44^. The tilt series acquisition parameters used for the TITAN Krios IP (a) and the TITAN Krios UZH (b) were set as follows. Tilt series were acquired at a calibrated pixel size of 1.59 Å (a), 1.75 Å (b), 1.9 Å (a) or 2.21 Å (b). Two types of tilt-series acquisition schemes were used: a dose-symmetric scheme^45^ starting at 0 ° from -51 ° to +51 ° with a 3 ° increment (a) and a bidirectional scheme starting at -30 ° from -60 ° to +60 ° with a 3 ° increment (b). The projection dose was calibrated for each sample to reach a total dose of approximately 120 to 140 e^−^/A^2^ and the defocus used was between −2 and -6 µm.

### Cryo-EM data processing and visualization

For each tilt series acquired, frame alignment was performed using either MotionCor2 1.6.4 or the alignframes program from the IMOD software package (RRID:SCR_003297)^46^. IMOD was also used for tilt alignment using fiducial markers and tomogram reconstruction at a binning factor of 2 or 4 with an antialiasing filter using dose weighting and a SIRT-like filter with 10-20 iterations. For tomograms selected as representative (binned by a factor of 4), fiducials were erased with the IMOD findbeads3d tool. These tomograms were further subjected to topaz denoising^47^ using a default 10 Å/px pre-trained model, for visualization purposes. A maximum of 10 tomographic slices was summed using the IMOD slicer tool to increase contrast and shown as representative tomographic slices of the tomograms containing features of interest. Tomographic slices may have been rotated along the x and y axis for presentation and comparison purposes.

For representative tomograms showing tip stages, intracellular and extracellular features, IMOD models were manually prepared for the membrane, cell wall and outer layer and for the features to be highlighted in each model. Linear interpolation was performed for all models except for vesicles (in which spherical interpolation was performed), flagella, filament-like structures and tracks (for which no interpolation was performed). IMOD meshes were prepared for all features except tracks. Upon volume meshing, the cap and tube tool were used for filament-like structures and vesicles and for flagella, respectively, to ensure the features were represented as closed models. IMOD models for the three layers (membrane, cell wall and outer layer), vesicle membranes, intracellular filaments and flagella were used as segmentation masks in UCSF ChimeraX version 1.7 (RRID:SCR_015872)^48^ to prepare volume masks using the slab tool with the width measured for each feature. The outer layer corresponded to the electron dense layer seen in the S-layer and in the hair-like layer. The pad tool was used to represent the tomogram densities from the cytosol, the interior of extracellular vesicles and the chemosensory array while the coating of extracellular vesicles was defined using the slab tool. Tracks and hair-like structures were represented as IMOD contours, and their thickness adjusted using the atom and stickRadius commands. Final segmentation models were clipped, rotated and colored to show the features of interest.

Projection images selected as representative were binned 4 times and opened in FIJI (RRID:SCR_002285)^49^. Contrast was adjusted and a Gaussian low-pass filter with 0.5 to 1 px radius was applied using Fiji’s unsharped mask tool, for visualization purposes. No image treatment was applied to images acquired using the 200 kV Tecnai F20 (FEI company). For two representative images of morphological stages, IMOD models were prepared for the membrane, cell wall, outer layer and hair-like structures. IMOD models were opened as contours in Chimera X and their thickness adjusted using the atom and stickRadius commands. The cytoplasm of each original projection image was isolated using the imodmop command and included in the segmentation model prepared. A cytoplasm mask was defined and the segmentation models colored to show the features of interest.

### Cryo-EM data analysis

To measure SFB length, projection images containing a full SFB in the field of view were opened in IMOD or in FIJI and a contour/line was traced from the SFB back to the edge of the tip. Tip length was measured by tracing a contour/line from the base to the tip edge, as shown in Fig. 6k-l. To measure vesicle diameters, projection images or tomograms (binned 4 times) acquired in a 300kV electron microscope were open in IMOD, a circle shape was used to establish the contour of each vesicle at its maximum diameter and its area was measured using the imodinfo tool. Vesicle diameter was then deduced from the circle area formula (πr²).

Measurements of intracellular filaments, hair-like structures length and width and distance between hair-like structures and S-layer subunits from SFB tip tomograms (binned 4 times) was performed manually in IMOD by defining individual contours and using the imodinfo tool. SFB tip diameter was measured as described above in a tomographic slice from the center of the volume.

At least five tomograms were selected for the measurements of each feature/stage with the exception of plate-like structures since these filament-like structures were only present in three tomograms for mouse-SFB-NL and two tomograms for rat-SFB-Yit. Tip diameter and length and vesicle diameter was measured once per tomogram/feature. Ten measurements were performed for each filament-like structure and for the distance between S-layer and hair-like layer subunits. Ten measurements in five different tomographic slices were performed to estimate hair-like structures length and width. Individual measurement values and their corresponding mean were plotted while mean and standard deviation were disclosed in the text. The distance between S-layer and hair-like layer subunits was also confirmed by selecting regions from tomograms binned by a factor of 2 that contained clear S-layer or HLL repeats and by using a Fast Fourier transform in Fiji to assess repetitiveness. Plot profiles of the selected regions were also obtained and the distances between two peaks were measured.

### Histological analysis

Tissue from the terminal ileum of rodents monocolonized with SFB was fixed in 4% paraformaldehyde (Electron Microscopy Sciences) and preserved in OCT (TFS). Sections of 10 µm thickness were cut using a cryostat (Leica), fixed in 4% paraformaldehyde and stained for 10 minutes using Giemsa staining (Sigma Aldrich). Imaging was performed using an Olympus BX53 microscope.

### Protein expression and purification of Th17Ag

The Th17Ag gene from mouse-SFB-NL was synthesized with *Escherichia coli* codon optimization and cloned in the plasmid pET151/D-TOPO (GeneArt, TFS). Protein expression of Th17Ag in the BL21 (DE3) Star *E. coli* strain was performed at 16 °C in Luria-Bertani broth with an overnight induction using 1 mM Isopropyl β-d-1-thiogalactopyranoside (IPTG) (Sigma Aldrich). The recombinantly expressed Th17Ag was produced with a poly-histidine and V5 epitope-containing N-terminal tag. The tagged protein was purified by affinity chromatography followed by size-exclusion using a Cobalt 5 mL HiTrap TALON crude column (Cytiva) and a Superdex 200 Increase 10/300 GL column (Cytiva), respectively. Protein purification was performed using an AKTA-FPLC purification system following the instructions of the manufacturer. On-tube digestion using Tobacco Etch Virus (TEV) protease (PX P1108, Proteogenix) was performed to cleave the tag from purified Th17Ag using 1 TEV unit per 6 µg of purified protein. Removal of the remaining tags and TEV was achieved through re-purification by affinity chromatography using a Cobalt 5 mL HiTrap TALON crude column (Cytiva) and a 5 mL GSTrap HP column (Cytiva). Protein-containing fractions were analyzed by sodium dodecyl sulfate–polyacrylamide gel electrophoresis (SDS-PAGE) and western blot using an anti-histidine antibody (Sigma Aldrich, 1:5000). The tagged protein was used for subsequent experiments unless otherwise stated.

### ELISA with polyclonal antibody against Th17Ag

Purified Th17Ag was coated onto Nunc MaxiSorp flat-bottom plates (TFS) at a concentration of 10 µg/mL at 4 °C overnight. Coated plates were washed four times with 0.05% tween 20 (Sigma Aldrich) in PBS 1X (TFS) and incubated for 1 h at room temperature in blocking buffer (0.05% tween 20 in PBS 1X with 1% bovine serum albumin, BSA, Sigma Aldrich). Rabbit polyclonal antibodies produced against the Th17Ag from mouse-SFB-NYU (1:100 of a 1:1 mix containing two rabbit anti-3340 antibodies)^7^ and a goat polyclonal antibody anti-rabbit conjugated with alkaline phosphatase (Sigma Aldrich 1:10 000) were subsequently added to each well in blocking buffer for 2 hours and 1 hour, respectively. Four washing steps were performed after each incubation. A solution of 4.5 M NaCl (Sigma Aldrich), 0.1 M Tris-HCl (TFS) and 1 mg/mL 4- nitrophenyl phosphate disodium salt (Sigma Aldrich) was used to reveal binding. The absorbance was measured at 415 nm using an iMark Microplate Absorbance Reader (BioRad). The polyclonal antibodies used were kindly provided by Dr. Dan R Littman.

### Nanobody library and phage selection

An alpaca (*Lama pacos*) named Picchu was immunized three times by subcutaneous injection with SFB purified from the intestinal content of monocolonized mice (1x10^9^ to 4x10^10^ SFB genomes) and fixed with 4% paraformaldehyde (PFA). Two extra booster immunizations were performed 3 months later with SFB purified from the SFB-TC7 cell co-culture (3x10^7^ to 2x10^9^ SFB genomes) described by Schnupf *et al*.^11^ and fixed with 4% PFA. SFB were mixed with complete Freund’s adjuvant for the first immunization and with incomplete Freund’s adjuvant for subsequent injections. Approximately 250 mL of alpaca blood were collected 6 days after the last immunization. Peripheral blood lymphocytes were isolated by Ficoll (Sigma Aldrich) density gradient centrifugation and stored at -80 °C until further use. Total RNA was extracted from the lymphocytes and converted to cDNA as described by Lafaye *et al*.^50^. DNA sequences encoding the nanobodies, naturally produced by the immunized alpaca (VHH, variable domain of heavy chain), were amplified by PCR from cDNA and cloned into pHEN6 phagemid vector which were then electroporated into *E. coli* TG1 to obtain a VHH library of approximately 10^8^ different clones. Phage-nanobodies from the VHH library were produced using the helper phage M13K7 (New England Biolabs) and Th17Ag-binding VHH were selected by phage display^51^. Briefly, 10^11^ Phage-VHH were incubated with purified Th17Ag previously coated on Nunc-Immunotubes (TFS) for 1 hour at room temperature. Three rounds of panning were performed with increasingly lower concentrations of coated Th17Ag, starting at 10 µg/mL, and increasingly higher numbers of extensive washes. Th17Ag-binding phage-VHH were eluted in 100 mM triethylamine (TEA, Sigma Aldrich) and used to infect *E. coli* TG1. Individual colonies were used to produce phage-VHH in 96-wells plates, which were then tested by phage-ELISA for binding to Th17Ag previously coated onto 96-wells Nunc MaxiSorp flat-bottom plates (TFS) using an anti-M13 monoclonal antibody conjugated to horseradish peroxidase (Abcam, 1:2500). The sequence of positive clones was determined by Sanger sequencing using the M13rev-29 primer (Eurofins).

### Nanobody expression and selection

Positive VHH were expressed from pHEN6 vectors with an N-terminal *pelB* signal sequence and a C-terminal c-Myc and poly-histidine-containing tag. Periplasmic expression was performed in *E. coli* TG1 after overnight induction at 30 °C with 1 mM IPTG (Euromedex). Bacterial pellets were resuspended in PBS 1X containing 300 mM NaCl, 1 mg/mL polymyxin B sulfate (Sigma Aldrich) and Complete, EDTA-free Protease Inhibitor Cocktail (Roche) and incubated at 4 °C for 1 h with vigorous shaking (300 rpm). Periplasmic extracts obtained by centrifugation and used to purify the produced nanobodies by affinity chromatography using a Cobalt 5 mL HiTrap TALON crude column (Cytiva) followed by size exclusion chromatography on a Superdex 75 pg column (Cytiva). An AKTA-Start purification system was used according to the manufacturer instructions. Purified VHH were concentrated and stored in PBS 1X at −20 °C until further use.

Binding of purified VHH (10 µg/mL) to Th17Ag was assessed by ELISA using a mouse monoclonal antibody anti-c-Myc (Bio-Techne, 1:500) and a goat anti-mouse polyclonal antibody coupled to alkaline phosphatase (Sigma Aldrich, 1:10 000).

To enable large scale VHH production, a positive VHH, with c-Myc and 6xHis tag followed by a stop codon, was subcloned into the vector pFuse which was transfected into Expi293 cells using the Expi293 Expression System Kit (TFS), according to the manufacturer’s instructions. Expi293 cells were grown at 37 °C for five days and VHH-Fc were purified from their supernatants by affinity chromatography as described above. Purified VHH were concentrated and stored in PBS 1X at - 20 °C until further use.

### Expression of nanobodies fused to Fc region

The coding region of the selected VHH was subcloned into the vector pFuse^52^ in order to allow the expression of dimeric VHH-Fc fusion proteins. VHH-Fc were expressed in Expi293 cells as described above, and purified using a HiTrap protein A HP (Cytiva) and an AKTA-Start purification system, according to the manufacturer instructions. Purified VHH were concentrated and stored in PBS 1X at −20 °C until further use.

Binding of purified VHH-Fc (10 µg/mL) to Th17Ag was assessed by ELISA using a mouse monoclonal antibody anti-human IgG1 Fc coupled to horseradish peroxidase (TFS, 1:1000) and Nunc MaxiSorp flat-bottom plates (TFS) coated with 10 µg/mL of purified protein. 3,3′,5,5′- Tetramethylbenzidine (TMB, Sigma Aldrich) was used to reveal binding following the manufacturer’s instructions. The absorbance was measured at 450 nm.

VHH-Fc was biotinylated using EZ-Link Sulfo NHS-Biotin (TFS) according to the manufacturer guidelines for subsequent use in immunofluorescence.

### Immunofluorescence and immunogold labelling of Th17Ag

Purified SFB from monocolonized mice (IOs only and filament-enriched fractions) were fixed in 4% PFA at room temperature for 30 minutes and kept in 1% PFA at 4 °C overnight. SFB samples were washed with PBS 1X and either left untreated (fixed SFB) or incubated in PBS 1X (TFS) with 0.5% triton X-100 (Sigma Aldrich) at 95 °C for 5 minutes followed by a PBS 1X wash (fixed and denatured SFB). Immunolabelling was performed in 1.5 mL tubes using PBS 1X with 1% BSA (Sigma Aldrich) as blocking buffer and PBS 1X as washing buffer. After 1 hour in blocking buffer, SFB were incubated either with rabbit polyclonal (1:100) or with VHH-Fc (10 µg/mL) in blocking buffer for 2 hours at room temperature. For immunogold labelling, Protein A coupled to 10 nm gold particles (Aurion, 1:20) was added for 1 h in blocking buffer. For immunofluorescence, streptavidin coupled to Alexa 568 (1:200, TFS) and a goat polyclonal antibody anti-rabbit coupled to Alexa 568 (1:500, TFS) was used for detection of labelling with the biotinylated VHH-Fc and the polyclonal antibody, respectively. DAPI (4 µg/mL, TFS) was added together with the fluorescently coupled antibody/streptavidin to make the identification of SFB easier. Two washing steps were performed after each incubation.

For immunogold labelling, samples were diluted from 1:4 to 1:2 and cryo-EM grids were prepared immediately after staining, as described above, and imaged using a 200 kV Tecnai F20 (FEI company) equipped with a direct detector Falcon II (TFS).

For immunofluorescence, samples were diluted from 1:10 to 1:20, dried onto glass slides and preserved in ProLong Gold antifade medium (TFS) at 4 °C until imaging by confocal microscopy using an SP8 confocal microscope (Leica).

### Quantification of immunogold labelling

Representative projection images acquired using a 200 kV Tecnai F20 (FEI company) were selected for each immunogold labelling condition and the ccderaser function of imod was used to erase the black pixel located at the center of the original images. Counting of gold particles co- localized with SFB cells was performed manually in IMOD. Images with over 10 gold particles in the background were not considered for analysis. Additionally, to avoid considering poorly labelled SFB, SFB were only considered labelled if co-localization with at least 20 gold particles was observed, or at least 10 gold particles if labelling was restricted to the SFB tip.

### VHH binding kinetics

The binding kinetics of the VHH anti-Th17Ag VHH were assisted by Biolayer Interferometry (BLI) using an Octet HTX system (Sartorius) at 30 °C. The His-tagged VHH was immobilized onto Ni-NTA sensor tips at 5 μg/mL for 180 seconds following a baseline in PBS 1X supplemented with tween 20 at 0.1% (PBS/T) for 120 seconds. The VHH-coated tips were then dipped into different concentrations of Th17Ag (2264, 905, 226.4, 90.5 and 0 nM) for 600 seconds to allow association. Subsequently, biosensors were placed in PBS/T for 15 min to initiate dissociation. Raw data was processed using Octet Data Analysis Studio (version 13.0) the data were fitted to a 1:1 Langmuir binding model.

### Identification of VHH binding epitopes on Th17Ag

A summary of the Hydrogen/ Deuterium eXchange-Mass Spectrometry (HDX-MS) data is provided in **Supplementary Table 4**^53^.The quality of Th17Ag was assessed by intact mass measurement (measured mass: 113 762.00 Da; expected mass: 113 760.7381 Da; Δm = 1.26 Da 11.1 ppm). Th17Ag alone (7.1 µM in PBS1X) or in complex with a 1.5X molar excess VHH was equilibrated for 30 min at room temperature. Continuous labelling was initiated by adding 107 µL of deuterated buffer (PBS1X, pD 7.45) to 13 µL of equilibrated protein solution, resulting in a final D_2_O/H_2_O ratio of 90/10. The experimental K_D_ value was used to adjust the protein concentrations so that more than 90% of complex remained during labelling. The exchange reaction was quenched after 10 seconds, 1, 10, 30, 60 and 120 minutes labelling at room temperature by mixing 20 µL of labelling reaction with 40 µL of an ice-cold solution of 0.15% formic acid and 6 M guanidinium chloride to reduce the pH to 2.5. A similar procedure was used for the preparation of undeuterated control samples. Quenched samples were snap-frozen in liquid nitrogen and stored at -80 °C until further use. Triplicate labelling experiments were performed for each time point and condition (1 biological and 2 technical replicates).

Quenched samples were injected onto a nanoACQUITY UPLC system (Waters Corporation) equipped with an HDX manager maintained at 0 °C. ∼11 pmol of labelled Th17Ag either alone or with 16.5 pmol of VHH were digested using an in-house packed cartridge (2.0 x 20 mm, 63 µL bed volume) of immobilized pepsin beads (Thermo Fisher Scientific) for 2 min at 20 °C. Peptides were directly trapped and desalted onto a C18 trap column (VanGuard BEH 1.7 mm, 2.1 x 5 mm, Waters Corporation) at a flow rate of 100 µL/min (0.15% formic acid, pH 2.5) and separated by a 8 min linear gradient of acetonitrile from 5 to 30%, followed by a 2 min increase from 30 to 40%, at 40 µL/min using an ACQUITY UPLC BEH C18 analytical column (1.7 µm, 1.0 mm × 100 mm, Waters Corporation). After each run, the pepsin column was manually cleaned with two consecutive washes of 1.0% formic acid, 5% acetonitrile, 1.5 M guanidinium chloride. Blank injections were performed after each sample to confirm the absence of carry-over.

Mass spectra were acquired in resolution and positive ion mode (*m/z* 50-1950) on a Synapt G2- Si HDMS mass spectrometer (Waters Corporation, Milford, MA) equipped with a standard ESI source and lock-mass correction. Peptides were identified in undeuterated samples by a combination of data independent acquisition (MS^E^) and exact mass measurement (below 5.0 ppm mass error) using the same chromatographic conditions as for the deuterated samples. To maximize sequence coverage, the fragmentation of Th17Ag peptide ions was conducted using four distinct collision energy ramps in the trap region of the instrument: Low: 10 to 30V; Med: 15 to 35V; High: 20 to 45V; Mix: 10 to 45V.

The initial peptide map of Th17Ag was generated by database searching in ProteinLynX Global server 3.0 (Waters corporation) using the following processing and workflow parameters: low and elevated intensity thresholds set to 250.0 and 100.0 counts; intensity threshold =750.0 counts; automatic peptide and fragment tolerance; non-specific primary digest reagent; false discovery rate = 4%. Each fragmentation spectrum was manually inspected for assignment confirmation. The peptide map was refined in DynamX 3.0 (Waters corporation, Milford, MA) using the following import PLGS results filters: minimum intensity = 10 000; minimum products per amino acid value = 0.3; minimum PLGS score = 7.0; maximum MH+ error = 5.0 ppm. A total of 154 peptides covering 90.6% of the Th17Ag protein sequence were selected and analyzed by HDX-MS.

DynamX 3.0 was used to extract the centroid masses of all peptides selected for HDX-MS analyses. Results are reported as relative deuterium exchange levels (no back exchange correction) expressed in either mass unit or as fractional exchange. Fractional exchange values were calculated by dividing the experimental uptake value by the theoretically maximum number of exchangeable backbone amide hydrogens that could be replaced in each peptide, considering the final excess of deuterium present in the labelling mixture.

To facilitate HDX-MS data interpretation, models of Th17Ag were generated with Alphafold^54^ using the default parameters without energy minimization.

## Statistical analysis

Data was plotted and statistical analysis was performed using GraphPad Prism 9.10 (GraphPad Software Inc, *: *p* < 0.05, **: *p* < 0.01, ***: *p* < 0.001, ****: *p* < 0.0001). Statistical significance was assessed for individual measurements following a normal distribution using a t-test or a one-way ANOVA in cases where only two or multiple groups were compared, respectively. Statistical significance was assessed for individual measurements with a non-normal distribution using a Mann-Whitney U test or a Kruskal-Wallis test followed by a Dunn’s test in cases where only two or multiple groups were compared, respectively. To compare proportions between two groups, a Fisher’s exact test was used. The MEMHDX software was used to statistically validate HDX-MS datasets (Wald test, false discovery rate of 5%)^55^.

## Notes

### Competing Interest Statement

The authors have declared no competing interest.

## References

1. Daniel, N., Lécuyer, E. & Chassaing, B. Host/microbiota interactions in health and diseases—Time for mucosal microbiology! Mucosal Immunol 14, 1006–1016 (2021).

2. Gaboriau-Routhiau, V., Rakotobe, S., Lécuyer, E., Mulder, I., Lan, A., Bridonneau, C., Rochet, V., Pisi, A., De Paepe, M., Brandi, G., Eberl, G., Snel, J., Kelly, D. & Cerf-Bensussan, N. The Key Role of Segmented Filamentous Bacteria in the Coordinated Maturation of Gut Helper T Cell Responses. Immunity 31, 677–689 (2009).

3. Ivanov, I. I., Atarashi, K., Manel, N., Brodie, E. L., Shima, T., Karaoz, U., Wei, D., Goldfarb, K. C., Santee, C. A., Lynch, S. V., Tanoue, T., Imaoka, A., Itoh, K., Takeda, K., Umesaki, Y., Honda, K. & Littman, D. R. Induction of Intestinal Th17 Cells by Segmented Filamentous Bacteria. Cell 139, 485– 498 (2009).

4. Goto, Y., Obata, T., Kunisawa, J., Sato, S., Ivanov, I. I., Lamichhane, A., Takeyama, N., Kamioka, M., Sakamoto, M., Matsuki, T., Setoyama, H., Imaoka, A., Uematsu, S., Akira, S., Domino, S. E., Kulig, P., Becher, B., Renauld, J. C., Sasakawa, C., Umesaki, Y., Benno, Y. & Kiyono, H. Innate lymphoid cells regulate intestinal epithelial cell glycosylation. Science (1979) 345, 1254009 (2014).

5. Lécuyer, E., Rakotobe, S., Lengliné-Garnier, H., Lebreton, C., Picard, M., Juste, C., Fritzen, R., Eberl, G., McCoy, K. D., Macpherson, A. J., Reynaud, C. A., Cerf-Bensussan, N. & Gaboriau-Routhiau, V. Segmented filamentous bacterium uses secondary and tertiary lymphoid tissues to induce gut IgA and specific T helper 17 cell responses. Immunity 40, 608–620 (2014).

6. Brabec, T., Schwarzer, M., Kováčová, K., Dobešová, M., Schierová, D., Březina, J., Pacáková, I., Šrůtková, D., Ben-Nun, O., Goldfarb, Y., Šplíchalová, I., Kolář, M., Abramson, J., Filipp, D. & Dobeš, J. Segmented filamentous bacteria–induced epithelial MHCII regulates cognate CD4+ IELs and epithelial turnover. Journal of Experimental Medicine 221, e20230194 (2024).

7. Yang, Y., Torchinsky, M. B., Gobert, M., Xiong, H., Xu, M., Linehan, J. L., Alonzo, F., Ng, C., Chen, A., Lin, X., Sczesnak, A., Liao, J. J., Torres, V. J., Jenkins, M. K., Lafaille, J. J. & Littman, D. R. Focused specificity of intestinal TH17 cells towards commensal bacterial antigens. Nature 510, 152–156 (2014).

8. Gauguet, S., D’Ortona, S., Ahnger-Pier, K., Duan, B., Surana, N. K., Lu, R., Cywes-Bentley, C., Gadjeva, M., Shan, Q., Priebe, G. P. & Pier, G. B. Intestinal microbiota of mice influences resistance to Staphylococcus aureus pneumonia. Infect Immun 83, 4003–4014 (2015).

9. Ngo, V. L., Lieber, C. M., Kang, H. ji, Sakamoto, K., Kuczma, M., Plemper, R. K. & Gewirtz, A. T. Intestinal microbiota programming of alveolar macrophages influences severity of respiratory viral infection. Cell Host Microbe 32, 335–348.e8 (2024).

10. Schnupf, P., Gaboriau-Routhiau, V., Sansonetti, P. J. & Cerf-Bensussan, N. Segmented filamentous bacteria, Th17 inducers and helpers in a hostile world. Curr Opin Microbiol 35, 100–109 (2017).

11. Schnupf, P., Gaboriau-Routhiau, V., Gros, M., Friedman, R., Moya-Nilges, M., Nigro, G., Cerf- Bensussan, N. & Sansonetti, P. J. Growth and host interaction of mouse segmented filamentous bacteria in vitro. Nature 520, 99–103 (2015).

12. Klaasen, H. L. B. M., Koopman, J. P., Poelma, F. G. J. & Beynen, A. C. Intestinal, segmented, filamentous bacteria. FEMS Microbiol Lett 88, 165–180 (1992).

13. Schnupf, P., Gaboriau-Routhiau, V. & Cerf-Bensussan, N. Host interactions with Segmented Filamentous Bacteria: An unusual trade-off that drives the post-natal maturation of the gut immune system. Semin Immunol 25, 342–351 (2013).

14. Atarashi, K., Tanoue, T., Ando, M., Kamada, N., Nagano, Y., Narushima, S., Suda, W., Imaoka, A., Setoyama, H., Nagamori, T., Ishikawa, E., Shima, T., Hara, T., Kado, S., Jinnohara, T., Ohno, H., Kondo, T., Toyooka, K., Watanabe, E., Yokoyama, S. I., Tokoro, S., Mori, H., Noguchi, Y., Morita, H., Ivanov, I. I., Sugiyama, T., Nuñez, G., Camp, J. G., Hattori, M., Umesaki, Y. & Honda, K. Th17 Cell Induction by Adhesion of Microbes to Intestinal Epithelial Cells. Cell 163, 367–380 (2015).

15. Tannock, G. W., Miller, J. R. & Savage, D. C. Host Specificity of Filamentous, Segmented Microorganisms Adherent to the Small Bowel Epithelium in Mice and Rats. Appl Environ Microbiol 47, 441–442 (1984).

16. Jepson, M. A., Clark, M. A., Simmons, N. L. & Hirst, B. H. Actin accumulation at sites of attachment of indigenous apathogenic segmented filamentous bacteria to mouse ileal epithelial cells. Infect Immun 61, 4001–4004 (1993).

17. Chase, D. G. & Erlandsen, S. L. Evidence for a Complex Life Cycle and Endospore Formation in the Attached, Filamentous, Segmented Bacterium from Murine Ileum. J Bacteriol 127, 572–583 (1976).

18. Ladinsky, M. S., Araujo, L. P., Zhang, X., Veltri, J., Galan-Diez, M., Soualhi, S., Lee, C., Irie, K., Pinker, E. Y., Narushima, S., Bandyopadhyay, S., Nagayama, M., Elhenawy, W., Coombes, B. K., Ferraris, R. P., Honda, K., Iliev, I. D., Gao, N., Bjorkman, P. J. & Ivanov, I. I. Endocytosis of commensal antigens by intestinal epithelial cells regulates mucosal T cell homeostasis. Science 363, eaat4042 (2019).

19. Prakash, T., Oshima, K., Morita, H., Fukuda, S., Imaoka, A., Kumar, N., Sharma, V. K., Kim, S. W., Takahashi, M., Saitou, N., Taylor, T. D., Ohno, H., Umesaki, Y. & Hattori, M. Complete genome sequences of rat and mouse segmented filamentous bacteria, a potent inducer of Th17 cell differentiation. Cell Host Microbe 10, 273–284 (2011).

20. Bolotin, A., de Wouters, T., Schnupf, P., Bouchier, C., Loux, V., Rhimi, M., Jamet, A., Dervyn, R., Boudebbouze, S., Blottière, H. M., Sorokin, A., Snel, J., Cerf-Bensussan, N., Gaboriau-Routhiau, V., van de Guchte, M. & Maguin, E. Genome sequence of ‘Candidatus Arthromitus’ sp. strain SFB- mouse-NL, a commensal bacterium with a key role in postnatal maturation of gut immune functions. Genome Announc 2, e00705–e00714 (2014).

21. Sczesnak, A., Segata, N., Qin, X., Gevers, D., Petrosino, J. F., Huttenhower, C., Littman, D. R. & Ivanov, I. I. The genome of Th17 cell-inducing segmented filamentous bacteria reveals extensive auxotrophy and adaptations to the intestinal environment. Cell Host Microbe 10, 260–272 (2011).

22. Pamp, S. J., Harrington, E. D., Quake, S. R., Relman, D. A. & Blainey, P. C. Single-cell sequencing provides clues about the host interactions of segmented filamentous bacteria (SFB). Genome Res 22, 1107–1119 (2012).

23. Klaasen, H. L. B. M., Van Der Heijden, P. J., Stok, W., Poelma, F. G. J., Koopman, J. P., Van Den Brink, M. E., Bakker, M. H., Eling, W. M. C. & Beynen, A. C. Apathogenic, Intestinal, Segmented, Filamentous Bacteria Stimulate the Mucosal Immune System of Mice. Infect Immun 61, 303–306 (1993).

24. Klaasen, H. L. B. M., Koopman, J. P., Van Den Brink, M. E., Van Wezel, H. P. N. & Beynen, A. C. Mono- association of mice with non-cultivable, intestinal, segmented, filamentous bacteria. Arch Microbiol 156, 148–151 (1991).

25. Snel, J., Heinen, P. P., Blok, H. J., Carman, R. J., Duncan, A. J., Allen, P. C. & COLLINSh, M. D. Comparison of 16s rRNA Sequences of Segmented Filamentous Bacteria Isolated from Mice, Rats, and Chickens and Proposal of ‘Candidatus Arthromitus’. International Union of Microbiological Societies 45, (1995).

26. Matias, V. R. F. & Beveridge, T. J. Cryo-electron microscopy reveals native polymeric cell wall structure in Bacillus subtilis 168 and the existence of a periplasmic space. Mol Microbiol 56, 240– 251 (2005).

27. Bharat, T. A. M., Kureisaite-Ciziene, D., Hardy, G. G., Yu, E. W., Devant, J. M., Hagen, W. J. H., Brun, Y. V., Briggs, J. A. G. & Löwe, J. Structure of the hexagonal surface layer on Caulobacter crescentus cells. Nat Microbiol 2, 1–6 (2017).

28. Lanzoni-mangutchi, P., Banerji, O., Wilson, J., Barwinska-sendra, A., Kirk, J. A., Vaz, F., Beirne, S. O., Baslé, A., Omari, K. El, Wagner, A., Fairweather, N. F., Douce, G. R., Bullough, P. A., Fagan, R. P. & Salgado, P. S. Structure and assembly of the S-layer in C. difficile. Nat Commun 13, 970 (2022).

29. Tatli, M., Moraïs, S., Tovar-Herrera, O. E., Bomble, Y. J., Bayer, E. A., Medalia, O. & Mizrahi, I. Nanoscale resolution of microbial fiber degradation in action. Elife 11, e76523 (2022).

30. Fagan, R. P. & Fairweather, N. F. Biogenesis and functions of bacterial S-layers. Nat Rev Microbiol 12, 211–222 (2014).

31. Sleytr, U. B. & Beveridge, T. J. Bacterial S-layers. Trends Microbiol 7, 253–260 (1999).

32. Nkamba, I., Mulet, C., Dubey, G. P., Gorgette, O., Couesnon, A., Salles, A., Moya-Nilges, M., Jung, V., Gaboriau-Routhiau, V., Guerrera, I. C., Shima, T., Umesaki, Y., Nigro, G., Krijnse-Locker, J., Bérard, M., Cerf-Bensussan, N., Sansonetti, P. J. & Schnupf, P. Intracellular offspring released from SFB filaments are flagellated. Nat Microbiol 5, 34–39 (2020).

33. Ingerson-Mahar, M., Briegel, A., Werner, J. N., Jensen, G. J. & Gitai, Z. The metabolic enzyme CTP synthase forms cytoskeletal filaments. Nat Cell Biol 12, 739–746 (2010).

34. Mandelbaum, N., Zhang, L., Carasso, S., Ziv, T., Lifshiz-Simon, S., Davidovich, I., Luz, I., Berinstein, E., Gefen, T., Cooks, T., Talmon, Y., Balskus, E. P. & Geva-Zatorsky, N. Extracellular vesicles of the Gram-positive gut symbiont Bifidobacterium longum induce immune-modulatory, anti- inflammatory effects. NPJ Biofilms Microbiomes 9, (2023).

35. Brown, L., Wolf, J. M., Prados-rosales, R. & Casadevall, A. Through the wall : extracellular vesicles in Gram-positive bacteria , mycobacteria and fungi. Nat Rev Microbiol 13, (2015).

36. Smith, T. M. Segmented Filamentous Bacteria in the Bovine Small Intestine. J. Comp. Path 117, 185– 190 (1997).

37. Mignot, T., Mesnage, S., Couture-Tosi, E., Mock, M. & Fouet, A. Developmental switch of S-layer protein synthesis in Bacillus anthracis. Mol Microbiol 43, 1615–1627 (2002).

38. Sogues, A., Fioravanti, A., Jonckheere, W., Pardon, E., Steyaert, J. & Remaut, H. Structure and function of the EA1 surface layer of Bacillus anthracis. Nat Commun 14, 7051 (2023).

39. Neuhaus, A., Selvaraj, M., Salzer, R., Langer, J. D., Kruse, K., Kirchner, L., Sanders, K., Daum, B., Averhoff, B. & Gold, V. A. M. Cryo-electron microscopy reveals two distinct type IV pili assembled by the same bacterium. Nat Commun 11, 2231 (2020).

40. Li, G., Brown, P. J. B., Tang, J. X., Xu, J., Quardokus, E. M., Fuqua, C. & Brun, Y. V. Surface contact stimulates the just-in-time deployment of bacterial adhesins. Mol Microbiol 83, 41–51 (2012).

41. Klein, E. A., Schlimpert, S., Hughes, V., Brun, Y. V., Thanbichler, M. & Gitai, Z. Physiological role of stalk lengthening in caulobacter crescentus. Commun Integr Biol 6, (2013).

42. Wos-Oxley, M. L., Bleich, A., Oxley, A. P. A., Kahl, S., Janus, L. M., Smoczek, A., Nahrstedt, H., Pils, M. C., Taudien, S., Platzer, M., Hedrich, H. J., Medina, E. & Pieper, D. H. Comparative evaluation of establishing a human gut microbial community within rodent models. Gut Microbes 3, 1–16 (2012).

43. Wagner, F. R., Watanabe, R., Schampers, R., Singh, D., Persoon, H., Schaffer, M., Fruhstorfer, P., Plitzko, J. & Villa, E. Preparing samples from whole cells using focused-ion-beam milling for cryo- electron tomography. Nat Protoc 15, 2041–2070 (2020).

44. Mastronarde, D. N. SerialEM: A Program for Automated Tilt Series Acquisition on Tecnai Microscopes Using Prediction of Specimen Position. Microscopy and Microanalysis 9, 1182–1183 (2003).

45. Hagen WJH, Wan W, B. J. Implementation of a cryo-electron tomography tilt-scheme optimized for high resolution subtomogram averaging. J Struct Biol 197, 191–198 (2017).

46. Mastronarde, D. N. & Held, S. R. HHS Public Access. J Struct Biol 197, 102–113 (2017).

47. Bepler, T., Kelley, K., Noble, A. J. & Berger, B. Topaz-Denoise: general deep denoising models for cryoEM and cryoET. Nat Commun 11, 5208 (2020).

48. Goddard, T. D., Huang, C. C., Meng, E. C., Pettersen, E. F., Couch, G. S., Morris, J. H. & Ferrin, T. E. TOOLS FOR PROTEIN SCIENCE UCSF ChimeraX : Meeting modern challenges in visualization and analysis. Protein Science 27, 14–25 (2018).

49. Schindelin, J., Arganda-carreras, I., Frise, E., Kaynig, V., Pietzsch, T., Preibisch, S., Rueden, C., Saalfeld, S., Schmid, B., Tinevez, J., White, D. J., Hartenstein, V., Tomancak, P. & Cardona, A. Fiji - an Open Source platform for biological image analysis. Nat Methods 9, (2012).

50. Lafaye, P., Achour, I., England, P., Duyckaerts, C. & Rougeon, F. Single-domain antibodies recognize selectively small oligomeric forms of amyloid β, prevent Aβ-induced neurotoxicity and inhibit fibril formation. Mol Immunol 46, 695–704 (2009).

51. Gransagne, M., Aymé, G., Brier, S., Chauveau-Le Friec, G., Meriaux, V., Nowakowski, M., Dejardin, F., Levallois, S., de Melo, G. D., Donati, F., Prot, M., Brûlé, S., Raynal, B., Bellalou, J., Goncalves, P., Montagutelli, X., Di Santo, J. P., Lazarini, F., England, P., Petres, S., Escriou, N. & Lafaye, P. Development of a highly specific and sensitive VHH-based sandwich immunoassay for the detection of the SARS-CoV-2 nucleoprotein. Journal of Biological Chemistry 298, (2022).

52. Moutel, S., El Marjou, A., Vielemeyer, O., Nizak, C., Benaroch, P., Dübel, S. & Perez, F. A multi-Fc- species system for recombinant antibody production. BMC Biotechnol 9, (2009).

53. Masson, G. R., Burke, J. E., Ahn, N. G., Anand, G. S., Borchers, C., Brier, S., Bou-Assaf, G. M., Engen, J. R., Englander, S. W., Faber, J., Garlish, R., Griffin, P. R., Gross, M. L., Guttman, M., Hamuro, Y., Heck, A. J. R., Houde, D., Iacob, R. E., Jørgensen, T. J. D., Kaltashov, I. A., Klinman, J. P., Konermann, L., Man, P., Mayne, L., Pascal, B. D., Reichmann, D., Skehel, M., Snijder, J., Strutzenberg, T. S., Underbakke, E. S., Wagner, C., Wales, T. E., Walters, B. T., Weis, D. D., Wilson, D. J., Wintrode, P. L., Zhang, Z., Zheng, J., Schriemer, D. C. & Rand, K. D. Recommendations for performing, interpreting and reporting hydrogen deuterium exchange mass spectrometry (HDX-MS) experiments. Nat Methods 16, 595–602 (2019).

54. Jumper, J., Evans, R., Pritzel, A., Green, T., Figurnov, M., Ronneberger, O., Tunyasuvunakool, K., Bates, R., Žídek, A., Potapenko, A., Bridgland, A., Meyer, C., Kohl, S. A. A., Ballard, A. J., Cowie, A., Romera-Paredes, B., Nikolov, S., Jain, R., Adler, J., Back, T., Petersen, S., Reiman, D., Clancy, E., Zielinski, M., Steinegger, M., Pacholska, M., Berghammer, T., Bodenstein, S., Silver, D., Vinyals, O., Senior, A. W., Kavukcuoglu, K., Kohli, P. & Hassabis, D. Highly accurate protein structure prediction with AlphaFold. Nature 596, 583–589 (2021).

55. Hourdel, V., Volant, S., O’Brien, D. P., Chenal, A., Chamot-Rooke, J., Dillies, M. A. & Brier, S. MEMHDX: An interactive tool to expedite the statistical validation and visualization of large HDX- MS datasets. Bioinformatics 32, 3413–3419 (2016).

56. Perez-Riverol, Y., Bai, J., Bandla, C., García-Seisdedos, D., Hewapathirana, S., Kamatchinathan, S., Kundu, D. J., Prakash, A., Frericks-Zipper, A., Eisenacher, M., Walzer, M., Wang, S., Brazma, A. & Vizcaíno, J. A. The PRIDE database resources in 2022: A hub for mass spectrometry-based proteomics evidences. Nucleic Acids Res 50, D543–D552 (2022).

