## Supplementary File for "Segmented filamentous bacteria undergo a structural transition at their adhesive tip during unicellular to filament development"

#### Content:

- **Supplementary Table 1:** Characteristics of all SFB analyzed.
- **Supplementary Table 2:** Characteristics of vesicles, intracellular filaments and plate-like structures identified at the SFB tip.
- **Supplementary Table 3:** Binding affinity between the VHH anti-Th17Ag and Th17Ag determined Biolayer Interferometry (BLI).
- **Supplementary Table 4:** HDX-MS summary table.
- **Supplementary Fig. 1:** Top view of the S-layer arrangement at the SFB tip.
- **Supplementary Fig. 2:** Additional examples of the intracellular and extracellular features identified in SFB and related to Figure 2.
- **Supplementary Fig. 3:** Differences in tip length between mouse-SFB and rat-SFB.
- **Supplementary Fig. 4:** Additional examples of the developmental stages of mouse-SFB IOs, related to figure 3.
- **Supplementary Fig. 5:** Identification of undefined SFB stages.
- **Supplementary Fig. 6:** Assessment of the presence of the hair-like layer at the SFB back.
- **Supplementary Fig. 7:** Potential transitional stage from stage 3 to stage 4.
- **Supplementary Fig. 8:** Assessment of host-specificity of SFB attachment.
- **Supplementary Fig. 9:** Purification and detection of a Th17 antigen (Th17Ag).
- **Supplementary Fig. 10:** Negative controls of immunofluorescence and immunogold labelling.
- **Supplementary Fig. 11:** Additional data for the localization of Th17Ag at the SFB surface.
- **Supplementary Fig. 12:** Kinetic and HDX-MS analysis of Th17Ag binding to monovalent VHH anti-Th17Ag.
- **Supplementary Movie Legends**

### Supplementary Tables

### Supplementary Table 1

Cruz, *et al.*, 2025

**Supplementary Table 1. Characteristics of all SFB analyzed.** HLL: hair-like layer, NA: not assessed, UD: unidentified.

| Experiment no. | Electron microscope | Projection Image/Tomogram | SFB strain | Host species | SFB size (µm) | Tip length (nm) | Tip diameter (nm) | SFB tip Stage | SFB back stage | Presence of flagella | Presence of chemosensory array | Presence of filaments | Presence of plate-like structures | Presence of intracellular vesicles | Tracks | Presence of extracellular vesicles | EMDB accession number |
|---|---|---|---|---|---|---|---|---|---|---|---|---|---|---|---|---|---|
| Exp1 | TITAN Krios UZH | Tomogram | Candidatus Arthromitus sp. strain SFB-mouse-NL | Mus musculus C57BL/6J | 2.4 | UD | NA | UD | NA | No | No | No | No | No | No | Undecorated | EMD-52675 (tip) |
| Exp1 | TITAN Krios UZH | Tomogram | Candidatus Arthromitus sp. strain SFB-mouse-NL | Mus musculus C57BL/6J | 3.9 | UD | NA | Stage 4/5_UD | NA | No | No | No | No | No | No | No | |
| Exp1 | TITAN Krios UZH | Tomogram | Candidatus Arthromitus sp. strain SFB-mouse-NL | Mus musculus C57BL/6J | 6.6 | UD | NA | Stage 4/5_UD | NA | No | No | No | No | No | No | No | |
| Exp1 | TITAN Krios UZH | Tomogram | Candidatus Arthromitus sp. strain SFB-mouse-NL | Mus musculus C57BL/6J | 5.2 | 368.4 | NA | Stage 4 | NA | No | No | No | No | No | No | No | |
| Exp1 | TITAN Krios UZH | Tomogram | Candidatus Arthromitus sp. strain SFB-mouse-NL | Mus musculus C57BL/6J | 4.8 | 552.2 | NA | Stage 4/5_UD | NA | No | No | No | No | No | No | No | |
| Exp1 | TITAN Krios UZH | Tomogram | Candidatus Arthromitus sp. strain SFB-mouse-NL | Mus musculus C57BL/6J | 2.2 | 442.1 | NA | Stage 3 | NA | No | No | No | No | No | No | No | |
| Exp1 | TITAN Krios UZH | Tomogram | Candidatus Arthromitus sp. strain SFB-mouse-NL | Mus musculus C57BL/6J | 9.0 | 504.4 | NA | Stage 4/5_UD | NA | No | No | No | No | Yes | No | No | |
| Exp1 | TITAN Krios UZH | Tomogram | Candidatus Arthromitus sp. strain SFB-mouse-NL | Mus musculus C57BL/6J | 7.7 | 440.6 | NA | Stage 5 | NA | No | No | No | No | No | No | No | |
| Exp1 | TITAN Krios UZH | Tomogram | Candidatus Arthromitus sp. strain SFB-mouse-NL | Mus musculus C57BL/6J | 10.0 | 627.2 | NA | Stage 4 | NA | No | No | No | No | Yes | No | No | |
| Exp1 | TITAN Krios UZH | Tomogram | Candidatus Arthromitus sp. strain SFB-mouse-NL | Mus musculus C57BL/6J | 2.3 | 400.5 | NA | Stage 3 | NA | No | No | No | No | No | No | No | |
| Exp1 | TITAN Krios UZH | Tomogram | Candidatus Arthromitus sp. strain SFB-mouse-NL | Mus musculus C57BL/6J | 2.0 | 456.2 | NA | Stage 3 | NA | No | No | No | No | No | No | No | |
| Exp1 | TITAN Krios UZH | Tomogram | Candidatus Arthromitus sp. strain SFB-mouse-NL | Mus musculus C57BL/6J | 2.2 | UD | NA | UD | NA | No | No | No | No | No | No | No | |
| Exp1 | Tecnai F20 | Projection Image | Candidatus Arthromitus sp. strain SFB-mouse-NL | Rattus norvegicus F344 | 2.2 | 491.9 | NA | Stage 3 | No HLL | NA | NA | NA | NA | NA | NA | NA | |
| Exp1 | Tecnai F20 | Projection Image | Candidatus Arthromitus sp. strain SFB-mouse-NL | Rattus norvegicus F344 | 2.1 | 503.4 | NA | Stage 3 | No HLL | NA | NA | NA | NA | NA | NA | NA | |
| Exp1 | Tecnai F20 | Projection Image | Candidatus Arthromitus sp. strain SFB-rat-Yit | Rattus norvegicus F344 | 2.2 | 382.5 | NA | Stage 1 | No HLL | Yes | NA | NA | NA | No | No | Decorated | |
| Exp1 | Tecnai F20 | Projection Image | Candidatus Arthromitus sp. strain SFB-rat-Yit | Rattus norvegicus F344 | 2.3 | 511.6 | NA | Stage 1 | No HLL | No | NA | NA | NA | No | No | Undecorated | |
| Exp1 | Tecnai F20 | Projection Image | Candidatus Arthromitus sp. strain SFB-rat-Yit | Rattus norvegicus F344 | 2.1 | NA | NA | UD | No HLL | Yes | NA | NA | NA | Yes | No | No | |
| Exp1 | Tecnai F20 | Projection Image | Candidatus Arthromitus sp. strain SFB-rat-Yit | Rattus norvegicus F344 | 2.0 | NA | NA | UD | No HLL | Yes | NA | NA | NA | No | No | Decorated and Undecorated | |
| Exp1 | Tecnai F20 | Projection Image | Candidatus Arthromitus sp. strain SFB-rat-Yit | Rattus norvegicus F344 | 2.1 | NA | NA | UD | No HLL | Yes | NA | NA | NA | No | No | Undecorated | |
| Exp1 | Tecnai F20 | Projection Image | Candidatus Arthromitus sp. strain SFB-rat-Yit | Rattus norvegicus F344 | 2.1 | NA | NA | UD | No HLL | Yes | NA | NA | NA | No | No | Undecorated | |
| Exp1 | Tecnai F20 | Projection Image | Candidatus Arthromitus sp. strain SFB-rat-Yit | Rattus norvegicus F344 | 2.4 | NA | NA | UD | UD | No | NA | NA | NA | No | No | Decorated and Undecorated | |
| Exp2 | TITAN Krios UZH | Tomogram | Candidatus Arthromitus sp. strain SFB-mouse-NL | Mus musculus C57BL/6J | 15.6 | 591.8 | 211.7 | Stage 5 | NA | No | No | No | No | Yes | Yes | No | EMD-52678 (tip) |
| Exp2 | TITAN Krios UZH | Tomogram | Candidatus Arthromitus sp. strain SFB-mouse-NL | Mus musculus C57BL/6J | 6.1 | 514.6 | NA | Stage 5 | NA | No | No | No | No | No | No | No | |
| Exp2 | TITAN Krios UZH | Tomogram | Candidatus Arthromitus sp. strain SFB-mouse-NL | Mus musculus C57BL/6J | 5.3 | 547.2 | NA | Stage 4/5_UD | NA | No | No | No | No | No | No | Undecorated | |
| Exp2 | TITAN Krios UZH | Tomogram | Candidatus Arthromitus sp. strain SFB-mouse-NL | Mus musculus C57BL/6J | 2.3 | 424.3 | NA | Stage 1 | NA | No | No | No | No | No | No | No | |
| Exp2 | TITAN Krios UZH | Tomogram | Candidatus Arthromitus sp. strain SFB-mouse-NL | Mus musculus C57BL/6J | 3.7 | 672.0 | NA | Stage 4/5_UD | NA | No | No | No | No | No | No | No | |
| Exp2 | TITAN Krios UZH | Tomogram | Candidatus Arthromitus sp. strain SFB-mouse-NL | Mus musculus C57BL/6J | 9.4 | 550.8 | NA | Stage 4 | NA | No | No | No | No | No | No | No | |
| Exp2 | TITAN Krios UZH | Tomogram | Candidatus Arthromitus sp. strain SFB-mouse-NL | Mus musculus C57BL/6J | 5.3 | 694.1 | NA | Stage 4 | NA | No | No | No | No | No | No | No | |
| Exp2 | TITAN Krios UZH | Tomogram | Candidatus Arthromitus sp. strain SFB-mouse-NL | Mus musculus C57BL/6J | 9.6 | 579.5 | 134.0 | Stage 4 | NA | No | No | No | No | Yes | Yes | No | |
| Exp2 | TITAN Krios UZH | Tomogram | Candidatus Arthromitus sp. strain SFB-mouse-NL | Mus musculus C57BL/6J | 3.3 | 468.3 | NA | Stage 5 | NA | No | No | No | No | No | Yes | No | |
| Exp2 | TITAN Krios UZH | Tomogram | Candidatus Arthromitus sp. strain SFB-mouse-NL | Mus musculus C57BL/6J | 6.0 | 491.0 | NA | Stage 4/5_UD | NA | No | No | No | No | No | No | No | |
| Exp2 | Tecnai F20 | Projection Image | Candidatus Arthromitus sp. strain SFB-mouse-NL | Rattus norvegicus F344 | 2.1 | 394.3 | NA | Stage 2 | No HLL | NA | NA | NA | NA | NA | NA | NA | |
| Exp2 | Tecnai F20 | Projection Image | Candidatus Arthromitus sp. strain SFB-mouse-NL | Rattus norvegicus F344 | 2.2 | 395.3 | NA | Stage 2 | NA | NA | NA | NA | NA | NA | NA | NA | |
| Exp2 | Tecnai F20 | Projection Image | Candidatus Arthromitus sp. strain SFB-mouse-NL | Rattus norvegicus F344 | 23.9 | 167.3 | NA | Stage 4/5_UD | HLL | NA | NA | NA | NA | NA | NA | NA | |
| Exp2 | Tecnai F20 | Projection Image | Candidatus Arthromitus sp. strain SFB-mouse-NL | Rattus norvegicus F344 | 29.2 | 329.7 | NA | Stage 4/5_UD | HLL | NA | NA | NA | NA | NA | NA | NA | |
| Exp2 | Tecnai F20 | Projection Image | Candidatus Arthromitus sp. strain SFB-mouse-NL | Rattus norvegicus F344 | 55.6 | 370.4 | NA | Stage 4/5_UD | HLL | NA | NA | NA | NA | NA | NA | NA | |
| Exp2 | Tecnai F20 | Projection Image | Candidatus Arthromitus sp. strain SFB-mouse-NL | Rattus norvegicus F344 | 43.8 | 288.9 | NA | Stage 4/5_UD | HLL | NA | NA | NA | NA | NA | NA | NA | |
| Exp2 | Tecnai F20 | Projection Image | Candidatus Arthromitus sp. strain SFB-mouse-NL | Rattus norvegicus F344 | 30.2 | 511.0 | NA | Stage 4/5_UD | HLL | NA | NA | NA | NA | NA | NA | NA | |
| Exp2 | Tecnai F20 | Projection Image | Candidatus Arthromitus sp. strain SFB-mouse-NL | Rattus norvegicus F344 | 28.2 | 511.0 | NA | Stage 4/5_UD | HLL | NA | NA | NA | NA | NA | NA | NA | |
| Exp2 | Tecnai F20 | Projection Image | Candidatus Arthromitus sp. strain SFB-rat-Yit | Rattus norvegicus F344 | 14.4 | 343.1 | NA | Stage 4/5_UD | HLL | No | NA | NA | NA | No | No | No | |
| Exp2 | Tecnai F20 | Projection Image | Candidatus Arthromitus sp. strain SFB-rat-Yit | Rattus norvegicus F344 | 2.4 | NA | NA | UD | No HLL | Yes | NA | NA | NA | No | No | Decorated | |
| Exp3 | TITAN Krios IP | Tomogram | Candidatus Arthromitus sp. strain SFB-rat-Yit | Rattus norvegicus F344 | 2.0 | NA | NA | UD | No HLL | No | No | No | Yes | No | No | No | EMD-52691 (tip) |
| Exp3 | TITAN Krios IP | Tomogram | Candidatus Arthromitus sp. strain SFB-rat-Yit | Rattus norvegicus F344 | 2.0 | NA | NA | UD | UD | No | No | Yes | No | Yes | No | Decorated and Undecorated | EMD-52693 (tip) |
| Exp3 | TITAN Krios IP | Tomogram | Candidatus Arthromitus sp. strain SFB-rat-Yit | Rattus norvegicus F344 | 2.0 | NA | NA | UD | NA | Yes | No | No | Yes | No | No | Undecorated | EMD-52694 (tip) |
| Exp3 | TITAN Krios IP | Tomogram | Candidatus Arthromitus sp. strain SFB-mouse-NL | Rattus norvegicus F344 | 1.6 | 570.6 | NA | Stage 3 | No HLL | NA | NA | NA | NA | NA | NA | NA | EMD-52702 (tip) |
| Exp3 | Tecnai F20 | Projection Image | Candidatus Arthromitus sp. strain SFB-rat-Yit | Rattus norvegicus F344 | 11.3 | 162.6 | NA | Stage 4/5_UD | HLL | No | NA | NA | NA | No | No | No | |
| Exp3 | TITAN Krios IP | Projection Image | Candidatus Arthromitus sp. strain SFB-rat-Yit | Rattus norvegicus F344 | 22.6 | 264.5 | NA | Stage 5 | HLL | No | NA | NA | NA | Yes | No | No | |
| Exp3 | TITAN Krios IP | Projection Image | Candidatus Arthromitus sp. strain SFB-rat-Yit | Rattus norvegicus F344 | 2.4 | 357.7 | NA | Stage 3 | No HLL | Yes | NA | NA | NA | No | No | Decorated | |
| Exp3 | Tecnai F20 | Projection Image | Candidatus Arthromitus sp. strain SFB-mouse-NL | Mus musculus C57BL/6J | 2.7 | 788.7 | NA | Stage 1 | No HLL | Yes | NA | NA | NA | No | No | No | |
| Exp3 | Tecnai F20 | Projection Image | Candidatus Arthromitus sp. strain SFB-mouse-NL | Mus musculus C57BL/6J | 7.3 | 685.8 | NA | Stage 4/5_UD | HLL | No | NA | NA | NA | No | No | No | |
| Exp3 | Tecnai F20 | Projection Image | Candidatus Arthromitus sp. strain SFB-mouse-NL | Mus musculus C57BL/6J | 2.7 | 1059.8 | NA | Stage 1 | No HLL | No | NA | NA | NA | No | No | Undecorated | |
| Exp3 | Tecnai F20 | Projection Image | Candidatus Arthromitus sp. strain SFB-mouse-NL | Mus musculus C57BL/6J | 2.4 | 718.5 | NA | Stage 4/5_UD | HLL | No | NA | NA | NA | No | No | No | |
| Exp3 | Tecnai F20 | Projection Image | Candidatus Arthromitus sp. strain SFB-mouse-NL | Mus musculus C57BL/6J | 2.3 | 716.1 | NA | Stage 4/5_UD | HLL | No | NA | NA | NA | No | Yes | Undecorated | |
| Exp3 | Tecnai F20 | Projection Image | Candidatus Arthromitus sp. strain SFB-mouse-NL | Mus musculus C57BL/6J | 2.6 | 1096.2 | NA | Stage 1 | No HLL | No | NA | NA | NA | No | No | Undecorated | |
| Exp3 | Tecnai F20 | Projection Image | Candidatus Arthromitus sp. strain SFB-mouse-NL | Mus musculus C57BL/6J | 2.7 | 573.7 | NA | Stage 4/5_UD | HLL | No | NA | NA | NA | No | Yes | No | |
| Exp3 | Tecnai F20 | Projection Image | Candidatus Arthromitus sp. strain SFB-mouse-NL | Mus musculus C57BL/6J | 3.0 | 955.5 | NA | Stage 4/5_UD | HLL | No | NA | NA | NA | No | Yes | No | |
| Exp3 | Tecnai F20 | Projection Image | Candidatus Arthromitus sp. strain SFB-mouse-NL | Mus musculus C57BL/6J | 10.8 | 872.2 | NA | Stage 4/5_UD | HLL | No | NA | NA | NA | No | No | No | |
| Exp3 | Tecnai F20 | Projection Image | Candidatus Arthromitus sp. strain SFB-mouse-NL | Mus musculus C57BL/6J | 2.7 | 971.4 | NA | Stage 2 | No HLL | No | NA | NA | NA | Yes | No | No | |
| Exp3 | Tecnai F20 | Projection Image | Candidatus Arthromitus sp. strain SFB-mouse-NL | Mus musculus C57BL/6J | 2.2 | 570.0 | NA | Stage 2 | No HLL | No | NA | NA | NA | No | No | No | |
| Exp3 | Tecnai F20 | Projection Image | Candidatus Arthromitus sp. strain SFB-mouse-NL | Mus musculus C57BL/6J | 12.7 | 570.5 | NA | Stage 4/5_UD | HLL | No | NA | NA | NA | Yes | Yes | No | |
| Exp3 | Tecnai F20 | Projection Image | Candidatus Arthromitus sp. strain SFB-mouse-NL | Mus musculus C57BL/6J | 10.7 | 712.6 | NA | Stage 4/5_UD | HLL | No | NA | NA | NA | Yes | Yes | No | |
| Exp3 | Tecnai F20 | Projection Image | Candidatus Arthromitus sp. strain SFB-mouse-NL | Mus musculus C57BL/6J | 11.7 | 735.9 | NA | Stage 4/5_UD | HLL | No | NA | NA | NA | No | No | No | |
| Exp3 | Tecnai F20 | Projection Image | Candidatus Arthromitus sp. strain SFB-mouse-NL | Mus musculus C57BL/6J | 14.0 | 322.2 | NA | Stage 4/5_UD | HLL | No | NA | NA | NA | No | Yes | No | |
| Exp3 | Tecnai F20 | Projection Image | Candidatus Arthromitus sp. strain SFB-mouse-NL | Mus musculus C57BL/6J | 6.9 | 736.3 | NA | Stage 4/5_UD | HLL | No | NA | NA | NA | Yes | No | No | |
| Exp3 | Tecnai F20 | Projection Image | Candidatus Arthromitus sp. strain SFB-mouse-NL | Mus musculus C57BL/6J | 7.7 | 912.7 | NA | Stage 4/5_UD | HLL | No | NA | NA | NA | Yes | Yes | No | |
| Exp3 | Tecnai F20 | Projection Image | Candidatus Arthromitus sp. strain SFB-mouse-NL | Mus musculus C57BL/6J | 4.3 | 798.1 | NA | Stage 4/5_UD | HLL | No | NA | NA | NA | No | No | Undecorated | |
| Exp3 | Tecnai F20 | Projection Image | Candidatus Arthromitus sp. strain SFB-mouse-NL | Mus musculus C57BL/6J | 2.5 | 748.2 | NA | Stage 3 | No HLL | No | NA | NA | NA | No | No | Decorated | |
| Exp3 | Tecnai F20 | Projection Image | Candidatus Arthromitus sp. strain SFB-mouse-NL | Mus musculus C57BL/6J | 11.0 | 991.0 | NA | Stage 4/5_UD | HLL | No | NA | NA | NA | Yes | No | No | |
| Exp3 | Tecnai F20 | Projection Image | Candidatus Arthromitus sp. strain SFB-mouse-NL | Mus musculus C57BL/6J | 13.0 | UD | NA | UD | NA | No | NA | NA | NA | No | No | No | |
| Exp3 | Tecnai F20 | Projection Image | Candidatus Arthromitus sp. strain SFB-mouse-NL | Mus musculus C57BL/6J | 2.2 | UD | NA | UD | No HLL | No | NA | NA | NA | Yes | Yes | No | |
| Exp3 | Tecnai F20 | Projection Image | Candidatus Arthromitus sp. strain SFB-mouse-NL | Mus musculus C57BL/6J | 7.3 | UD | NA | UD | NA | No | NA | NA | NA | No | No | No | |
| Exp3 | Tecnai F20 | Projection Image | Candidatus Arthromitus sp. strain SFB-mouse-NL | Mus musculus C57BL/6J | 11.3 | UD | NA | UD | NA | No | NA | NA | NA | No | No | Undecorated | |
| Exp3 | Tecnai F20 | Projection Image | Candidatus Arthromitus sp. strain SFB-mouse-NL | Rattus norvegicus F344 | 2.0 | 234.5 | NA | Stage 1 | No HLL | NA | NA | NA | NA | NA | NA | NA | |
| Exp3 | Tecnai F20 | Projection Image | Candidatus Arthromitus sp. strain SFB-mouse-NL | Rattus norvegicus F344 | 2.2 | 479.4 | NA | Stage 1 | No HLL | NA | NA | NA | NA | NA | NA | NA | |
| Exp3 | Tecnai F20 | Projection Image | Candidatus Arthromitus sp. strain SFB-mouse-NL | Rattus norvegicus F344 | 2.0 | 405.7 | NA | Stage 2 | No HLL | NA | NA | NA | NA | NA | NA | NA | |
| Exp3 | Tecnai F20 | Projection Image | Candidatus Arthromitus sp. strain SFB-mouse-NL | Rattus norvegicus F344 | 2.3 | 359.6 | NA | Stage 1 | No HLL | NA | NA | NA | NA | NA | NA | NA | |
| Exp3 | Tecnai F20 | Projection Image | Candidatus Arthromitus sp. strain SFB-mouse-NL | Rattus norvegicus F344 | 10.3 | 272.1 | NA | Stage 4/5_UD | HLL | NA | NA | NA | NA | NA | NA | NA | |
| Exp3 | Tecnai F20 | Projection Image | Candidatus Arthromitus sp. strain SFB-mouse-NL | Rattus norvegicus F344 | 37.9 | 522.6 | NA | Stage 4/5_UD | HLL | NA | NA | NA | NA | NA | NA | NA | |
| Exp3 | Tecnai F20 | Projection Image | Candidatus Arthromitus sp. strain SFB-mouse-NL | Rattus norvegicus F344 | 42.3 | 511.6 | NA | Stage 4/5_UD | HLL | NA | NA | NA | NA | NA | NA | NA | |
| Exp3 | TITAN Krios IP | Projection Image | Candidatus Arthromitus sp. strain SFB-mouse-NL | Rattus norvegicus F344 | 2.2 | 478.3 | NA | Stage 2 | No HLL | NA | NA | NA | NA | NA | NA | NA | |
| Exp3 | TITAN Krios IP | Projection Image | Candidatus Arthromitus sp. strain SFB-mouse-NL | Rattus norvegicus F344 | 2.1 | 401.0 | NA | Stage 2 | No HLL | NA | NA | NA | NA | NA | NA | NA | |
| Exp3 | TITAN Krios IP | Projection Image | Candidatus Arthromitus sp. strain SFB-mouse-NL | Rattus norvegicus F344 | 2.0 | 480.6 | NA | Stage 2 | No HLL | NA | NA | NA | NA | NA | NA | NA | |
| Exp3 | TITAN Krios IP | Projection Image | Candidatus Arthromitus sp. strain SFB-mouse-NL | Rattus norvegicus F344 | 2.3 | 391.6 | NA | Stage 2 | No HLL | NA | NA | NA | NA | NA | NA | NA | |
| Exp3 | TITAN Krios IP | Projection Image | Candidatus Arthromitus sp. strain SFB-mouse-NL | Rattus norvegicus F344 | 1.5 | 223.3 | NA | Stage 1 | No HLL | NA | NA | NA | NA | NA | NA | NA | |
| Exp3 | TITAN Krios IP | Projection Image | Candidatus Arthromitus sp. strain SFB-mouse-NL | Rattus norvegicus F344 | 1.7 | 330.1 | NA | Stage 2 | NA | NA | NA | NA | NA | NA | NA | NA | |
| Exp3 | TITAN Krios IP | Projection Image | Candidatus Arthromitus sp. strain SFB-mouse-NL | Rattus norvegicus F344 | 1.8 | 327.8 | NA | Stage 2 | No HLL | NA | NA | NA | NA | NA | NA | NA | |
| Exp3 | TITAN Krios IP | Projection Image | Candidatus Arthromitus sp. strain SFB-mouse-NL | Rattus norvegicus F344 | 12.9 | 368.3 | NA | Stage 5 | HLL | NA | NA | NA | NA | NA | NA | NA | |
| Exp3 | TITAN Krios IP | Tomogram | Candidatus Arthromitus sp. strain SFB-mouse-NL | Rattus norvegicus F344 | 2.4 | 532.3 | NA | Stage 2 | No HLL | NA | NA | NA | NA | NA | NA | NA | |
| Exp3 | TITAN Krios IP | Tomogram | Candidatus Arthromitus sp. strain SFB-mouse-NL | Rattus norvegicus F344 | 2.5 | 580.7 | NA | Stage 2 | No HLL | NA | NA | NA | NA | NA | NA | NA | |
| Exp3 | TITAN Krios IP | Tomogram | Candidatus Arthromitus sp. strain SFB-mouse-NL | Rattus norvegicus F344 | 1.8 | 359.0 | NA | Stage 2 | No HLL | NA | NA | NA | NA | NA | NA | NA | |
| Exp3 | TITAN Krios IP | Tomogram | Candidatus Arthromitus sp. strain SFB-mouse-NL | Rattus norvegicus F344 | 2.0 | 383.4 | NA | Stage 1 | No HLL | NA | NA | NA | NA | NA | NA | NA | |
| Exp3 | TITAN Krios IP | Tomogram | Candidatus Arthromitus sp. strain SFB-mouse-NL | Rattus norvegicus F344 | 2.1 | 529.5 | NA | Stage 2 | No HLL | NA | NA | NA | NA | NA | NA | NA | |
| Exp3 | TITAN Krios IP | Tomogram | Candidatus Arthromitus sp. strain SFB-mouse-NL | Rattus norvegicus F344 | 2.1 | 554.2 | NA | Stage 2 | No HLL | NA | NA | NA | NA | NA | NA | NA | |
| Exp3 | TITAN Krios IP | Tomogram | Candidatus Arthromitus sp. strain SFB-mouse-NL | Rattus norvegicus F344 | 2.2 | 562.0 | NA | Stage 3 | No HLL | NA | NA | NA | NA | NA | NA | NA | |
| Exp3 | TITAN Krios IP | Tomogram | Candidatus Arthromitus sp. strain SFB-mouse-NL | Rattus norvegicus F344 | 2.3 | 409.4 | NA | Stage 2 | No HLL | NA | NA | NA | NA | NA | NA | NA | |
| Exp3 | TITAN Krios IP | Tomogram | Candidatus Arthromitus sp. strain SFB-mouse-NL | Rattus norvegicus F344 | 2.0 | 445.5 | NA | Stage 2 | No HLL | NA | NA | NA | NA | NA | NA | NA | |
| Exp3 | TITAN Krios IP | Tomogram | Candidatus Arthromitus sp. strain SFB-mouse-NL | Rattus norvegicus F344 | 1.9 | 451.6 | NA | Stage 2 | No HLL | NA | NA | NA | NA | NA | NA | NA | |
| Exp3 | TITAN Krios IP | Tomogram | Candidatus Arthromitus sp. strain SFB-mouse-NL | Rattus norvegicus F344 | 2.0 | 504.8 | NA | Stage 2 | No HLL | NA | NA | NA | NA | NA | NA | NA | |
| Exp3 | TITAN Krios IP | Tomogram | Candidatus Arthromitus sp. strain SFB-mouse-NL | Rattus norvegicus F344 | 2.2 | 461.2 | NA | Stage 2 | No HLL | NA | NA | NA | NA | NA | NA | NA | |
| Exp3 | TITAN Krios IP | Tomogram | Candidatus Arthromitus sp. strain SFB-mouse-NL | Rattus norvegicus F344 | 2.1 | 423.6 | NA | Stage 1 | No HLL | NA | NA | NA | NA | NA | NA | NA | |
| Exp3 | TITAN Krios IP | Tomogram | Candidatus Arthromitus sp. strain SFB-mouse-NL | Rattus norvegicus F344 | 2.0 | 542.3 | NA | Stage 1 | NA | NA | NA | NA | NA | NA | NA | NA | |
| Exp3 | TITAN Krios IP | Tomogram | Candidatus Arthromitus sp. strain SFB-mouse-NL | Rattus norvegicus F344 | 2.6 | 471.4 | NA | Stage 2 | No HLL | NA | NA | NA | NA | NA | NA | NA | |
| Exp3 | TITAN Krios IP | Tomogram | Candidatus Arthromitus sp. strain SFB-mouse-NL | Rattus norvegicus F344 | 1.7 | 400.7 | NA | Stage 2 | No HLL | NA | NA | NA | NA | NA | NA | NA | |
| Exp3 | TITAN Krios IP | Tomogram | Candidatus Arthromitus sp. strain SFB-mouse-NL | Rattus norvegicus F344 | 2.1 | 498.7 | NA | Stage 3 | No HLL | NA | NA | NA | NA | NA | NA | NA | |
| Exp3 | TITAN Krios IP | Tomogram | Candidatus Arthromitus sp. strain SFB-mouse-NL | Rattus norvegicus F344 | 10.2 | 644.5 | NA | Stage 4 | NA | NA | NA | NA | NA | NA | NA | NA | |
| Exp3 | TITAN Krios IP | Tomogram | Candidatus Arthromitus sp. strain SFB-mouse-NL | Rattus norvegicus F344 | 30.5 | 180.5 | NA | Stage 5 | NA | NA | NA | NA | NA | NA | NA | NA | |
| Exp3 | TITAN Krios IP | Tomogram | Candidatus Arthromitus sp. strain SFB-mouse-NL | Rattus norvegicus F344 | 32.3 | 236.3 | NA | Stage 5 | NA | NA | NA | NA | NA | NA | NA | NA | |
| Exp3 | TITAN Krios IP | Tomogram | Candidatus Arthromitus sp. strain SFB-mouse-NL | Rattus norvegicus F344 | 14.7 | 301.0 | NA | Stage 5 | NA | NA | NA | NA | NA | NA | NA | NA | |
| Exp3 | TITAN Krios IP | Tomogram | Candidatus Arthromitus sp. strain SFB-mouse-NL | Rattus norvegicus F344 | 9.1 | 588.1 | NA | Stage 5 | NA | NA | NA | NA | NA | NA | NA | NA | |
| Exp3 | TITAN Krios IP | Tomogram | Candidatus Arthromitus sp. strain SFB-mouse-NL | Rattus norvegicus F344 | 1.9 | NA | NA | UD | NA | NA | NA | NA | NA | NA | NA | NA | |
| Exp3 | TITAN Krios IP | Projection Image | Candidatus Arthromitus sp. strain SFB-rat-Yit | Rattus norvegicus F344 | 40.7 | 359.6 | NA | Stage 5 | HLL | No | NA | NA | NA | Yes | Yes | No | |
| Exp3 | TITAN Krios IP | Tomogram | Candidatus Arthromitus sp. strain SFB-rat-Yit | Rattus norvegicus F344 | 14.3 | 379.8 | NA | Stage 5 | NA | No | No | No | No | No | No | No | |
| Exp3 | TITAN Krios IP | Tomogram | Candidatus Arthromitus sp. strain SFB-rat-Yit | Rattus norvegicus F344 | 25.7 | 494.3 | NA | Stage 5 | NA | No | No | No | No | Yes | No | No | |
| Exp3 | TITAN Krios IP | Projection Image | Candidatus Arthromitus sp. strain SFB-rat-Yit | Rattus norvegicus F344 | 1.9 | NA | NA | UD | NA | No | NA | NA | NA | No | No | Decorated | |
| Exp3 | TITAN Krios IP | Tomogram | Candidatus Arthromitus sp. strain SFB-rat-Yit | Rattus norvegicus F344 | 2.0 | NA | NA | UD | UD | No | No | No | No | No | No | Undecorated | |
| Exp3 | TITAN Krios IP | Tomogram | Candidatus Arthromitus sp. strain SFB-rat-Yit | Rattus norvegicus F344 | 2.4 | NA | NA | UD | UD | Yes | No | No | No | No | No | No | |
| Exp3 | TITAN Krios IP | Tomogram | Candidatus Arthromitus sp. strain SFB-rat-Yit | Rattus norvegicus F344 | 2.4 | NA | NA | UD | No HLL | No | No | Yes | No | No | No | No | |
| Exp3 | TITAN Krios IP | Projection Image | Candidatus Arthromitus sp. strain SFB-rat-Yit | Rattus norvegicus F344 | 2.0 | NA | NA | UD | NA | No | NA | NA | NA | No | No | Undecorated | |
| Exp3 | TITAN Krios IP | Tomogram | Candidatus Arthromitus sp. strain SFB-rat-Yit | Rattus norvegicus F344 | 2.2 | NA | NA | UD | No HLL | Yes | No | No | No | No | No | Undecorated | |
| Exp4 | TITAN Krios UZH | Tomogram | Candidatus Arthromitus sp. strain SFB-mouse-NL | Mus musculus C57BL/6J | 2.7 | 630.2 | 130.8 | Stage 2 | NA | Yes | No | Yes | No | Yes | Yes | No | EMD-52655 (tip) |
| Exp4 | TITAN Krios IP | Tomogram | Candidatus Arthromitus sp. strain SFB-rat-Yit | Rattus norvegicus F344 | 3.0 | 367.8 | NA | Stage 3 | NA | No | No | No | No | No | No | No | EMD-52688 (tip) |
| Exp4 | TITAN Krios IP | Tomogram | Candidatus Arthromitus sp. strain SFB-rat-Yit | Rattus norvegicus F344 | 2.4 | NA | NA | UD | No HLL | Yes | No | Yes | No | No | No | Decorated | EMD-52689 (tip) |
| Exp4 | TITAN Krios UZH | Tomogram | Candidatus Arthromitus sp. strain SFB-mouse-NL | Mus musculus C57BL/6J | 2.3 | NA | NA | NA | NA | No | Yes | Yes | No | No | No | No | EMD-52667 (tip) |
| Exp4 | TITAN Krios UZH | Tomogram | Candidatus Arthromitus sp. strain SFB-mouse-NL | Mus musculus C57BL/6J | 2.0 | 498.8 | 166.2 | Stage 2 | NA | No | No | Yes | No | No | No | No | EMD-52668 (tip) |
| Exp4 | TITAN Krios UZH | Tomogram | Candidatus Arthromitus sp. strain SFB-mouse-NL | Mus musculus C57BL/6J | 2.7 | UD | NA | Stage 3 | NA | No | No | No | Yes | No | No | No | EMD-52669 (tip) |
| Exp4 | TITAN Krios UZH | Tomogram | Candidatus Arthromitus sp. strain SFB-mouse-NL | Mus musculus C57BL/6J | 2.5 | NA | NA | NA | NA | No | No | No | No | Yes | Yes | Undecorated | EMD-52670 (tip) |
| Exp4 | TITAN Krios UZH | Tomogram | Candidatus Arthromitus sp. strain SFB-mouse-NL | Mus musculus C57BL/6J | 5.1 | 497.9 | NA | Stage 5 | NA | No | No | No | No | Yes | Yes | Decorated | EMD-52671 (tip) |
| Exp4 | TITAN Krios IP | Tomogram | Candidatus Arthromitus sp. strain SFB-rat-Yit | Rattus norvegicus F344 | 2.3 | 414.3 | NA | Stage 3 | No HLL | No | No | No | No | Yes | No | Decorated | |
| Exp4 | TITAN Krios IP | Tomogram | Candidatus Arthromitus sp. strain SFB-rat-Yit | Rattus norvegicus F344 | 2.4 | NA | NA | UD | UD | No | Yes | Yes | No | No | No | Decorated | EMD-52690 (tip) |
| Exp4 | TITAN Krios UZH | Tomogram | Candidatus Arthromitus sp. strain SFB-mouse-NL | Mus musculus C57BL/6J | 2.3 | NA | NA | NA | NA | No | Yes | No | Yes | Yes | Yes | No | EMD-52673 (tip) |
| Exp4 | TITAN Krios UZH | Tomogram | Candidatus Arthromitus sp. strain SFB-mouse-NL | Mus musculus C57BL/6J | 2.5 | NA | NA | NA | NA | No | No | Yes | No | No | No | No | EMD-52674 (tip) |
| Exp4 | TITAN Krios IP | Tomogram | Candidatus Arthromitus sp. strain SFB-rat-Yit | Rattus norvegicus F344 | 2.2 | NA | NA | UD | NA | Yes | Yes | Yes | No | No | No | Decorated | EMD-52692 (tip) |
| Exp4 | TITAN Krios UZH | Tomogram | Candidatus Arthromitus sp. strain SFB-mouse-NL | Mus musculus C57BL/6J | 3.1 | 614.6 | 154.3 | Stage 3 | NA | Yes | No | No | No | No | Yes | No | EMD-52676 (tip) |
| Exp4 | TITAN Krios UZH | Tomogram | Candidatus Arthromitus sp. strain SFB-mouse-NL | Mus musculus C57BL/6J | 12.0 | 665.7 | 210.5 | Stage 4 | NA | No | No | No | No | No | Yes | No | EMD-52677 (tip) |
| Exp4 | TITAN Krios UZH | Tomogram | Candidatus Arthromitus sp. strain SFB-mouse-NL | Mus musculus C57BL/6J | 2.3 | NA | NA | NA | HLL | NA | NA | NA | NA | NA | NA | NA | EMD-52679 (back) |
| Exp4 | TITAN Krios UZH | Tomogram | Candidatus Arthromitus sp. strain SFB-mouse-NL | Mus musculus C57BL/6J | 2.3 | 670.2 | 147.1 | Stage 3 | NA | No | No | Yes | No | No | No | No | EMD-52680 (tip) |
| Exp4 | TITAN Krios UZH | Tomogram | Candidatus Arthromitus sp. strain SFB-mouse-NL | Mus musculus C57BL/6J | 4.8 | 551.5 | NA | Stage 5 | NA | No | No | No | No | No | Yes | No | EMD-52682 (tip) |
| Exp4 | TITAN Krios UZH | Tomogram | Candidatus Arthromitus sp. strain SFB-mouse-NL | Mus musculus C57BL/6J | 3.3 | 549.5 | NA | Stage 4 | NA | No | No | No | No | No | No | No | EMD-52683 (tip) |
| Exp4 | TITAN Krios IP | Tomogram | Candidatus Arthromitus sp. strain SFB-rat-Yit | Rattus norvegicus F344 | 34.9 | 292.9 | NA | Stage 5 | NA | No | No | No | No | Yes | No | No | EMD-52699 (tip) |
| Exp4 | TITAN Krios IP | Tomogram | Candidatus Arthromitus sp. strain SFB-mouse-NL | Rattus norvegicus F344 | 2.1 | 690.3 | NA | Stage 1 | No HLL | NA | NA | NA | NA | NA | NA | NA | EMD-52700 (tip) |
| Exp4 | TITAN Krios IP | Projection Image | Candidatus Arthromitus sp. strain SFB-mouse-NL | Rattus norvegicus F344 | 2.3 | 547.9 | NA | Stage 2 | No HLL | NA | NA | NA | NA | NA | NA | NA | EMD-52701 (tip) |
| Exp4 | TITAN Krios IP | Tomogram | Candidatus Arthromitus sp. strain SFB-mouse-NL | Rattus norvegicus F344 | 3.7 | 365.9 | NA | Stage 5 | HLL | NA | NA | NA | NA | NA | NA | NA | EMD-52703 (tip) |
| Exp4 | Tecnai F20 | Projection Image | Candidatus Arthromitus sp. strain SFB-rat-Yit | Rattus norvegicus F344 | 20.7 | 230.1 | NA | Stage 4/5_UD | HLL | No | NA | NA | NA | No | No | No | |
| Exp4 | TITAN Krios IP | Tomogram | Candidatus Arthromitus sp. strain SFB-rat-Yit | Rattus norvegicus F344 | 13.7 | 358.7 | NA | Stage 5 | NA | No | No | No | No | No | No | No | |
| Exp4 | TITAN Krios IP | Tomogram | Candidatus Arthromitus sp. strain SFB-rat-Yit | Rattus norvegicus F344 | 19.4 | 389.1 | NA | Stage 5 | NA | No | No | No | No | Yes | No | No | |
| Exp4 | TITAN Krios IP | Projection Image | Candidatus Arthromitus sp. strain SFB-rat-Yit | Rattus norvegicus F344 | 2.1 | 392.3 | NA | Stage 3 | No HLL | No | NA | NA | NA | No | No | No | |
| Exp4 | TITAN Krios IP | Projection Image | Candidatus Arthromitus sp. strain SFB-rat-Yit | Rattus norvegicus F344 | 41.7 | 422.2 | NA | Stage 4 | NA | No | NA | NA | NA | No | Yes | No | |
| Exp4 | TITAN Krios IP | Tomogram | Candidatus Arthromitus sp. strain SFB-rat-Yit | Rattus norvegicus F344 | 2.3 | 434.0 | NA | UD | No HLL | No | No | No | No | No | No | No | |

### Supplementary Table 1

Cruz, *et al.*, 2025

**Supplementary Table 1. Characteristics of all SFB analyzed (Continuation).** HLL: hair-like layer, NA: not assessed, UD: unidentified.

| Experiment no. | Electron microscope | Projection Image/Tomogram | SFB strain | Host species | SFB size (µm) | Tip length (nm) | Tip diameter (nm) | SFB tip Stage | SFB back stage | Presence of flagella | Presence of chemosensory array | Presence of filaments | Presence of plate-like structures | Presence of intracellular vesicles | Tracks | Presence of extracellular vesicles | EMDB accession number |
|---|---|---|---|---|---|---|---|---|---|---|---|---|---|---|---|---|---|
| Exp4 | TITAN Krios IP | Tomogram | Candidatus Arthromitus sp. strain SFB-rat-Yit | Rattus norvegicus F344 | 32.8 | 439.6 | NA | Stage 5 | NA | No | No | No | No | No | No | No | |
| Exp4 | Tecnai F20 | Projection Image | Candidatus Arthromitus sp. strain SFB-rat-Yit | Rattus norvegicus F344 | 11.9 | 448.0 | NA | Stage 4/5_UD | HLL | No | NA | NA | NA | Yes | No | No | |
| Exp4 | Tecnai F20 | Projection Image | Candidatus Arthromitus sp. strain SFB-rat-Yit | Rattus norvegicus F344 | 25.3 | 464.6 | NA | Stage 4/5_UD | HLL | No | NA | NA | NA | No | No | No | |
| Exp4 | TITAN Krios IP | Projection Image | Candidatus Arthromitus sp. strain SFB-rat-Yit | Rattus norvegicus F344 | 11.4 | 469.7 | NA | Stage 4 | NA | No | NA | NA | NA | No | Yes | No | |
| Exp4 | Tecnai F20 | Projection Image | Candidatus Arthromitus sp. strain SFB-mouse-NL | Mus musculus C57BL/6J | 3.1 | 872.8 | NA | Stage 4/5_UD | HLL | No | NA | NA | NA | No | No | No | |
| Exp4 | Tecnai F20 | Projection Image | Candidatus Arthromitus sp. strain SFB-mouse-NL | Mus musculus C57BL/6J | 8.6 | 784.1 | NA | Stage 4/5_UD | HLL | No | NA | NA | NA | No | No | No | |
| Exp4 | Tecnai F20 | Projection Image | Candidatus Arthromitus sp. strain SFB-mouse-NL | Mus musculus C57BL/6J | 4.3 | 935.3 | NA | Stage 4/5_UD | HLL | No | NA | NA | NA | No | No | No | |
| Exp4 | Tecnai F20 | Projection Image | Candidatus Arthromitus sp. strain SFB-mouse-NL | Mus musculus C57BL/6J | 13.8 | 707.2 | NA | Stage 4/5_UD | HLL | No | NA | NA | NA | No | Yes | No | |
| Exp4 | Tecnai F20 | Projection Image | Candidatus Arthromitus sp. strain SFB-mouse-NL | Mus musculus C57BL/6J | 9.0 | 728.8 | NA | Stage 4/5_UD | HLL | No | NA | NA | NA | No | Yes | No | |
| Exp4 | Tecnai F20 | Projection Image | Candidatus Arthromitus sp. strain SFB-mouse-NL | Mus musculus C57BL/6J | 5.4 | 727.1 | NA | Stage 4/5_UD | HLL | No | NA | NA | NA | No | Yes | No | |
| Exp4 | Tecnai F20 | Projection Image | Candidatus Arthromitus sp. strain SFB-mouse-NL | Mus musculus C57BL/6J | 2.5 | 638.6 | NA | Stage 2 | No HLL | No | NA | NA | NA | No | Yes | No | |
| Exp4 | Tecnai F20 | Projection Image | Candidatus Arthromitus sp. strain SFB-mouse-NL | Mus musculus C57BL/6J | 3.2 | 685.6 | NA | Stage 4/5_UD | HLL | No | NA | NA | NA | Yes | Yes | No | |
| Exp4 | Tecnai F20 | Projection Image | Candidatus Arthromitus sp. strain SFB-mouse-NL | Mus musculus C57BL/6J | 3.6 | 760.8 | NA | Stage 4/5_UD | HLL | No | NA | NA | NA | No | No | No | |
| Exp4 | Tecnai F20 | Projection Image | Candidatus Arthromitus sp. strain SFB-mouse-NL | Mus musculus C57BL/6J | 3.4 | 845.2 | NA | Stage 4/5_UD | HLL | No | NA | NA | NA | No | No | No | |
| Exp4 | Tecnai F20 | Projection Image | Candidatus Arthromitus sp. strain SFB-mouse-NL | Mus musculus C57BL/6J | 4.9 | 954.2 | NA | Stage 4/5_UD | HLL | No | NA | NA | NA | No | Yes | No | |
| Exp4 | Tecnai F20 | Projection Image | Candidatus Arthromitus sp. strain SFB-mouse-NL | Mus musculus C57BL/6J | 2.2 | 552.5 | NA | Stage 1 | No HLL | No | NA | NA | NA | No | No | Undecorated | |
| Exp4 | Tecnai F20 | Projection Image | Candidatus Arthromitus sp. strain SFB-mouse-NL | Mus musculus C57BL/6J | 3.9 | 765.0 | NA | Stage 4/5_UD | HLL | No | NA | NA | NA | No | Yes | No | |
| Exp4 | Tecnai F20 | Projection Image | Candidatus Arthromitus sp. strain SFB-mouse-NL | Mus musculus C57BL/6J | 2.3 | 782.4 | NA | Stage 2 | No HLL | No | NA | NA | NA | No | No | No | |
| Exp4 | Tecnai F20 | Projection Image | Candidatus Arthromitus sp. strain SFB-mouse-NL | Mus musculus C57BL/6J | 3.9 | 636.1 | NA | Stage 4/5_UD | HLL | No | NA | NA | NA | No | No | No | |
| Exp4 | Tecnai F20 | Projection Image | Candidatus Arthromitus sp. strain SFB-mouse-NL | Mus musculus C57BL/6J | 8.1 | 490.7 | NA | Stage 4/5_UD | HLL | No | NA | NA | NA | No | No | No | |
| Exp4 | Tecnai F20 | Projection Image | Candidatus Arthromitus sp. strain SFB-mouse-NL | Mus musculus C57BL/6J | 13.9 | 748.8 | NA | Stage 4/5_UD | HLL | No | NA | NA | NA | No | No | No | |
| Exp4 | Tecnai F20 | Projection Image | Candidatus Arthromitus sp. strain SFB-mouse-NL | Mus musculus C57BL/6J | 14.2 | 650.8 | NA | Stage 4/5_UD | HLL | No | NA | NA | NA | No | No | No | |
| Exp4 | TITAN Krios UZH | Tomogram | Candidatus Arthromitus sp. strain SFB-mouse-NL | Mus musculus C57BL/6J | 1.7 | 379.8 | NA | Stage 2 | NA | No | No | No | No | No | No | No | |
| Exp4 | TITAN Krios UZH | Tomogram | Candidatus Arthromitus sp. strain SFB-mouse-NL | Mus musculus C57BL/6J | 4.0 | NA | NA | NA | NA | No | Yes | Yes | No | No | No | No | |
| Exp4 | TITAN Krios UZH | Tomogram | Candidatus Arthromitus sp. strain SFB-mouse-NL | Mus musculus C57BL/6J | 4.3 | 615.7 | NA | Stage 4/5_UD | NA | No | No | No | No | No | Yes | No | |
| Exp4 | TITAN Krios UZH | Tomogram | Candidatus Arthromitus sp. strain SFB-mouse-NL | Mus musculus C57BL/6J | 5.0 | 676.1 | NA | Stage 4 | NA | No | No | No | No | No | No | No | |
| Exp4 | TITAN Krios UZH | Tomogram | Candidatus Arthromitus sp. strain SFB-mouse-NL | Mus musculus C57BL/6J | 15.0 | 582.7 | NA | Stage 5 | NA | No | No | No | No | No | Yes | No | |
| Exp4 | TITAN Krios UZH | Tomogram | Candidatus Arthromitus sp. strain SFB-mouse-NL | Mus musculus C57BL/6J | 7.0 | 558.2 | 182.0 | Stage 5 | NA | No | No | No | No | No | No | No | |
| Exp4 | TITAN Krios UZH | Tomogram | Candidatus Arthromitus sp. strain SFB-mouse-NL | Mus musculus C57BL/6J | 7.0 | 527.8 | NA | Stage 4/5_UD | NA | No | No | No | No | No | Yes | No | |
| Exp4 | TITAN Krios UZH | Tomogram | Candidatus Arthromitus sp. strain SFB-mouse-NL | Mus musculus C57BL/6J | 2.3 | UD | NA | UD | NA | No | No | No | No | No | No | No | |
| Exp4 | TITAN Krios UZH | Tomogram | Candidatus Arthromitus sp. strain SFB-mouse-NL | Mus musculus C57BL/6J | 8.0 | 679.8 | NA | Stage 4/5_UD | NA | No | No | No | No | No | No | No | |
| Exp4 | TITAN Krios UZH | Tomogram | Candidatus Arthromitus sp. strain SFB-mouse-NL | Mus musculus C57BL/6J | 4.8 | 704.0 | NA | Stage 5 | NA | No | No | No | No | No | Yes | No | |
| Exp4 | TITAN Krios UZH | Tomogram | Candidatus Arthromitus sp. strain SFB-mouse-NL | Mus musculus C57BL/6J | 2.0 | 522.0 | NA | Stage 3 | NA | No | No | No | No | No | No | No | |
| Exp4 | TITAN Krios UZH | Tomogram | Candidatus Arthromitus sp. strain SFB-mouse-NL | Mus musculus C57BL/6J | 3.2 | 518.7 | NA | Stage 4 | NA | No | No | No | No | No | No | No | |
| Exp4 | TITAN Krios UZH | Tomogram | Candidatus Arthromitus sp. strain SFB-mouse-NL | Mus musculus C57BL/6J | 3.1 | 420.8 | NA | Stage 2 | NA | No | No | No | Yes | No | No | No | |
| Exp4 | TITAN Krios UZH | Tomogram | Candidatus Arthromitus sp. strain SFB-mouse-NL | Mus musculus C57BL/6J | 5.5 | 536.4 | NA | Stage 5 | NA | No | No | No | No | No | Yes | No | |
| Exp4 | TITAN Krios UZH | Tomogram | Candidatus Arthromitus sp. strain SFB-mouse-NL | Mus musculus C57BL/6J | 2.8 | 565.4 | NA | Stage 4/5_UD | NA | No | No | No | No | No | Yes | No | |
| Exp4 | TITAN Krios UZH | Tomogram | Candidatus Arthromitus sp. strain SFB-mouse-NL | Mus musculus C57BL/6J | 5.5 | 678.4 | NA | Stage 4/5_UD | NA | No | No | No | No | No | Yes | No | |
| Exp4 | TITAN Krios UZH | Tomogram | Candidatus Arthromitus sp. strain SFB-mouse-NL | Mus musculus C57BL/6J | 3.4 | 582.9 | NA | Stage 5 | NA | No | No | No | No | Yes | No | No | |
| Exp4 | TITAN Krios UZH | Tomogram | Candidatus Arthromitus sp. strain SFB-mouse-NL | Mus musculus C57BL/6J | 2.1 | 501.8 | NA | Stage 2 | NA | No | Yes | Yes | No | No | No | No | |
| Exp4 | TITAN Krios UZH | Tomogram | Candidatus Arthromitus sp. strain SFB-mouse-NL | Mus musculus C57BL/6J | 2.4 | UD | NA | Stage 4 | NA | No | No | No | No | No | No | No | |
| Exp4 | TITAN Krios UZH | Tomogram | Candidatus Arthromitus sp. strain SFB-mouse-NL | Mus musculus C57BL/6J | 2.5 | 566.1 | NA | Stage 4/5_UD | NA | No | No | Yes | No | No | No | No | |
| Exp4 | TITAN Krios UZH | Tomogram | Candidatus Arthromitus sp. strain SFB-mouse-NL | Mus musculus C57BL/6J | 2.1 | NA | NA | NA | NA | No | No | No | No | No | No | Decorated | |
| Exp4 | TITAN Krios UZH | Tomogram | Candidatus Arthromitus sp. strain SFB-mouse-NL | Mus musculus C57BL/6J | 7.5 | 627.6 | NA | Stage 4/5_UD | NA | No | No | No | No | Yes | No | No | |
| Exp4 | TITAN Krios UZH | Tomogram | Candidatus Arthromitus sp. strain SFB-mouse-NL | Mus musculus C57BL/6J | 4.5 | 777.6 | NA | Stage 4 | NA | No | No | No | No | No | Yes | No | |
| Exp4 | TITAN Krios UZH | Tomogram | Candidatus Arthromitus sp. strain SFB-mouse-NL | Mus musculus C57BL/6J | 3.4 | 541.2 | NA | Stage 4/5_UD | NA | No | No | No | No | No | No | Undecorated | |
| Exp4 | TITAN Krios UZH | Tomogram | Candidatus Arthromitus sp. strain SFB-mouse-NL | Mus musculus C57BL/6J | 2.4 | 560.5 | NA | Stage 4 | NA | No | No | No | No | No | No | No | |
| Exp4 | TITAN Krios UZH | Tomogram | Candidatus Arthromitus sp. strain SFB-mouse-NL | Mus musculus C57BL/6J | 2.9 | UD | NA | Stage 3 | NA | No | No | No | No | No | No | No | |
| Exp4 | TITAN Krios UZH | Tomogram | Candidatus Arthromitus sp. strain SFB-mouse-NL | Mus musculus C57BL/6J | 2.0 | 592.9 | NA | Stage 1 | NA | No | No | No | No | No | Yes | No | |
| Exp4 | TITAN Krios UZH | Tomogram | Candidatus Arthromitus sp. strain SFB-mouse-NL | Mus musculus C57BL/6J | 2.2 | 466.7 | NA | Stage 1 | NA | No | No | No | No | No | No | No | |
| Exp4 | Tecnai F20 | Projection Image | Candidatus Arthromitus sp. strain SFB-mouse-NL | Mus musculus C57BL/6J | 11.9 | UD | NA | UD | NA | No | NA | NA | NA | No | No | No | |
| Exp4 | TITAN Krios UZH | Tomogram | Candidatus Arthromitus sp. strain SFB-mouse-NL | Mus musculus C57BL/6J | 2.5 | UD | NA | UD | NA | No | No | Yes | No | No | No | No | |
| Exp4 | TITAN Krios UZH | Tomogram | Candidatus Arthromitus sp. strain SFB-mouse-NL | Mus musculus C57BL/6J | 2.7 | UD | NA | UD | NA | No | No | No | No | No | No | No | |
| Exp4 | TITAN Krios UZH | Tomogram | Candidatus Arthromitus sp. strain SFB-mouse-NL | Mus musculus C57BL/6J | 2.3 | UD | NA | UD | NA | No | No | No | No | No | Yes | No | |
| Exp4 | Tecnai F20 | Projection Image | Candidatus Arthromitus sp. strain SFB-mouse-NL | Rattus norvegicus F344 | 2.4 | 594.3 | NA | Stage 1 | No HLL | NA | NA | NA | NA | NA | NA | NA | |
| Exp4 | Tecnai F20 | Projection Image | Candidatus Arthromitus sp. strain SFB-mouse-NL | Rattus norvegicus F344 | 2.1 | 467.6 | NA | Stage 2 | No HLL | NA | NA | NA | NA | NA | NA | NA | |
| Exp4 | Tecnai F20 | Projection Image | Candidatus Arthromitus sp. strain SFB-mouse-NL | Rattus norvegicus F344 | 2.5 | 517.3 | NA | Stage 2 | No HLL | NA | NA | NA | NA | NA | NA | NA | |
| Exp4 | Tecnai F20 | Projection Image | Candidatus Arthromitus sp. strain SFB-mouse-NL | Rattus norvegicus F344 | 17.0 | 389.6 | NA | Stage 4/5_UD | HLL | NA | NA | NA | NA | NA | NA | NA | |
| Exp4 | Tecnai F20 | Projection Image | Candidatus Arthromitus sp. strain SFB-mouse-NL | Rattus norvegicus F344 | 34.5 | 528.2 | NA | Stage 4/5_UD | HLL | NA | NA | NA | NA | NA | NA | NA | |
| Exp4 | Tecnai F20 | Projection Image | Candidatus Arthromitus sp. strain SFB-mouse-NL | Rattus norvegicus F344 | 48.8 | 595.7 | NA | Stage 4/5_UD | HLL | NA | NA | NA | NA | NA | NA | NA | |
| Exp4 | Tecnai F20 | Projection Image | Candidatus Arthromitus sp. strain SFB-mouse-NL | Rattus norvegicus F344 | 8.7 | 105.5 | NA | Stage 4/5_UD | HLL | NA | NA | NA | NA | NA | NA | NA | |
| Exp4 | TITAN Krios IP | Projection Image | Candidatus Arthromitus sp. strain SFB-mouse-NL | Rattus norvegicus F344 | 2.2 | 562.0 | NA | Stage 1 | No HLL | NA | NA | NA | NA | NA | NA | NA | |
| Exp4 | TITAN Krios IP | Projection Image | Candidatus Arthromitus sp. strain SFB-mouse-NL | Rattus norvegicus F344 | 2.4 | 512.3 | NA | Stage 3 | No HLL | NA | NA | NA | NA | NA | NA | NA | |
| Exp4 | TITAN Krios IP | Projection Image | Candidatus Arthromitus sp. strain SFB-mouse-NL | Rattus norvegicus F344 | 44.7 | 630.7 | NA | Stage 5 | HLL | NA | NA | NA | NA | NA | NA | NA | |
| Exp4 | TITAN Krios IP | Projection Image | Candidatus Arthromitus sp. strain SFB-mouse-NL | Rattus norvegicus F344 | 47.2 | 213.4 | NA | Stage 5 | HLL | NA | NA | NA | NA | NA | NA | NA | |
| Exp4 | TITAN Krios IP | Projection Image | Candidatus Arthromitus sp. strain SFB-mouse-NL | Rattus norvegicus F344 | 8.2 | 389.1 | NA | Stage 4 | NA | NA | NA | NA | NA | NA | NA | NA | |
| Exp4 | TITAN Krios IP | Projection Image | Candidatus Arthromitus sp. strain SFB-mouse-NL | Rattus norvegicus F344 | 67.5 | 457.2 | NA | Stage 4/5_UD | HLL | NA | NA | NA | NA | NA | NA | NA | |
| Exp4 | TITAN Krios IP | Tomogram | Candidatus Arthromitus sp. strain SFB-mouse-NL | Rattus norvegicus F344 | 2.2 | 324.0 | NA | Stage 2 | No HLL | NA | NA | NA | NA | NA | NA | NA | |
| Exp4 | TITAN Krios IP | Tomogram | Candidatus Arthromitus sp. strain SFB-mouse-NL | Rattus norvegicus F344 | 2.3 | 434.8 | NA | Stage 2 | No HLL | NA | NA | NA | NA | NA | NA | NA | |
| Exp4 | TITAN Krios IP | Tomogram | Candidatus Arthromitus sp. strain SFB-mouse-NL | Rattus norvegicus F344 | 2.5 | 632.6 | NA | Stage 3 | No HLL | NA | NA | NA | NA | NA | NA | NA | |
| Exp4 | TITAN Krios IP | Tomogram | Candidatus Arthromitus sp. strain SFB-mouse-NL | Rattus norvegicus F344 | 2.3 | 575.1 | NA | Stage 2 | No HLL | NA | NA | NA | NA | NA | NA | NA | |
| Exp4 | TITAN Krios IP | Tomogram | Candidatus Arthromitus sp. strain SFB-mouse-NL | Rattus norvegicus F344 | 2.0 | 455.8 | NA | Stage 2 | No HLL | NA | NA | NA | NA | NA | NA | NA | |
| Exp4 | TITAN Krios IP | Tomogram | Candidatus Arthromitus sp. strain SFB-mouse-NL | Rattus norvegicus F344 | 2.1 | 615.0 | NA | Stage 2 | No HLL | NA | NA | NA | NA | NA | NA | NA | |
| Exp4 | TITAN Krios IP | Tomogram | Candidatus Arthromitus sp. strain SFB-mouse-NL | Rattus norvegicus F344 | 5.2 | 683.9 | NA | Stage 4/5_UD | HLL | NA | NA | NA | NA | NA | NA | NA | |
| Exp4 | TITAN Krios IP | Tomogram | Candidatus Arthromitus sp. strain SFB-mouse-NL | Rattus norvegicus F344 | 2.6 | 736.1 | NA | Stage 2 | No HLL | NA | NA | NA | NA | NA | NA | NA | |
| Exp4 | TITAN Krios IP | Tomogram | Candidatus Arthromitus sp. strain SFB-mouse-NL | Rattus norvegicus F344 | 2.1 | 302.9 | NA | Stage 1 | No HLL | NA | NA | NA | NA | NA | NA | NA | |
| Exp4 | TITAN Krios IP | Tomogram | Candidatus Arthromitus sp. strain SFB-mouse-NL | Rattus norvegicus F344 | 2.4 | 551.9 | NA | Stage 1 | No HLL | NA | NA | NA | NA | NA | NA | NA | |
| Exp4 | TITAN Krios IP | Tomogram | Candidatus Arthromitus sp. strain SFB-mouse-NL | Rattus norvegicus F344 | 2.3 | 405.8 | NA | Stage 2 | No HLL | NA | NA | NA | NA | NA | NA | NA | |
| Exp4 | TITAN Krios IP | Tomogram | Candidatus Arthromitus sp. strain SFB-mouse-NL | Rattus norvegicus F344 | 2.1 | 398.6 | NA | Stage 1 | No HLL | NA | NA | NA | NA | NA | NA | NA | |
| Exp4 | TITAN Krios IP | Tomogram | Candidatus Arthromitus sp. strain SFB-mouse-NL | Rattus norvegicus F344 | 36.0 | 578.8 | NA | Stage 5 | NA | NA | NA | NA | NA | NA | NA | NA | |
| Exp4 | TITAN Krios IP | Tomogram | Candidatus Arthromitus sp. strain SFB-mouse-NL | Rattus norvegicus F344 | 6.8 | 649.6 | NA | Stage 4/5_UD | NA | NA | NA | NA | NA | NA | NA | NA | |
| Exp4 | TITAN Krios IP | Tomogram | Candidatus Arthromitus sp. strain SFB-mouse-NL | Rattus norvegicus F344 | 25.7 | 622.3 | NA | Stage 4/5_UD | NA | NA | NA | NA | NA | NA | NA | NA | |
| Exp4 | Tecnai F20 | Projection Image | Candidatus Arthromitus sp. strain SFB-rat-Yit | Rattus norvegicus F344 | 1.9 | NA | NA | UD | UD | Yes | NA | NA | NA | No | No | Decorated | |
| Exp4 | Tecnai F20 | Projection Image | Candidatus Arthromitus sp. strain SFB-rat-Yit | Rattus norvegicus F344 | 2.0 | NA | NA | UD | No HLL | No | NA | NA | NA | No | No | Decorated | |
| Exp4 | TITAN Krios IP | Tomogram | Candidatus Arthromitus sp. strain SFB-rat-Yit | Rattus norvegicus F344 | 2.1 | NA | NA | UD | UD | Yes | No | Yes | No | No | No | Decorated | EMD-52856 (back) |
| Exp4 | TITAN Krios IP | Tomogram | Candidatus Arthromitus sp. strain SFB-rat-Yit | Rattus norvegicus F344 | 2.2 | NA | NA | UD | No HLL | No | No | No | No | No | No | Decorated and Undecorated | |
| Exp4 | TITAN Krios IP | Projection Image | Candidatus Arthromitus sp. strain SFB-rat-Yit | Rattus norvegicus F344 | 2.2 | NA | NA | UD | NA | No | NA | NA | NA | No | No | Decorated | |
| Exp4 | TITAN Krios IP | Tomogram | Candidatus Arthromitus sp. strain SFB-rat-Yit | Rattus norvegicus F344 | 2.2 | NA | NA | UD | No HLL | Yes | No | No | No | No | No | Decorated | |
| Exp4 | TITAN Krios IP | Tomogram | Candidatus Arthromitus sp. strain SFB-rat-Yit | Rattus norvegicus F344 | 2.3 | NA | NA | UD | No HLL | Yes | No | No | No | No | No | Decorated and Undecorated | |
| Exp4 | TITAN Krios IP | Projection Image | Candidatus Arthromitus sp. strain SFB-rat-Yit | Rattus norvegicus F344 | 2.3 | NA | NA | UD | NA | Yes | NA | NA | NA | Yes | No | Decorated | |
| Exp4 | Tecnai F20 | Projection Image | Candidatus Arthromitus sp. strain SFB-rat-Yit | Rattus norvegicus F344 | 2.4 | NA | NA | UD | No HLL | No | NA | NA | NA | No | No | Undecorated | |
| Exp4 | TITAN Krios IP | Tomogram | Candidatus Arthromitus sp. strain SFB-rat-Yit | Rattus norvegicus F344 | 2.5 | NA | NA | UD | No HLL | Yes | No | No | No | No | No | Decorated | |
| Exp5 | TITAN Krios IP | Tomogram | Candidatus Arthromitus sp. strain SFB-rat-Yit | Rattus norvegicus F344 | 2.6 | 611.8 | NA | Stage 1 | No HLL | No | No | Yes | No | Yes | No | No | EMD-52684 (tip); EMD-52698 (back) |
| Exp5 | TITAN Krios IP | Tomogram | Candidatus Arthromitus sp. strain SFB-rat-Yit | Rattus norvegicus F344 | 2.2 | 233.3 | NA | Stage 1 | NA | Yes | No | No | No | No | Yes | No | EMD-52695 (tip) |
| Exp5 | Tecnai F20 | Projection Image | Candidatus Arthromitus sp. strain SFB-rat-Yit | Rattus norvegicus F344 | 21.4 | 130.2 | NA | Stage 4/5_UD | HLL | No | NA | NA | NA | Yes | No | Decorated | |
| Exp5 | Tecnai F20 | Projection Image | Candidatus Arthromitus sp. strain SFB-rat-Yit | Rattus norvegicus F344 | 27.7 | 215.6 | NA | Stage 4/5_UD | HLL | No | NA | NA | NA | Yes | No | No | |
| Exp5 | Tecnai F20 | Projection Image | Candidatus Arthromitus sp. strain SFB-rat-Yit | Rattus norvegicus F344 | 2.0 | 252.8 | NA | Stage 1 | No HLL | Yes | NA | NA | NA | No | No | No | |
| Exp5 | TITAN Krios IP | Tomogram | Candidatus Arthromitus sp. strain SFB-rat-Yit | Rattus norvegicus F344 | 2.2 | 323.4 | NA | Stage 3 | NA | Yes | No | No | No | Yes | No | No | |
| Exp5 | Tecnai F20 | Projection Image | Candidatus Arthromitus sp. strain SFB-rat-Yit | Rattus norvegicus F344 | 2.3 | 356.3 | NA | Stage 1 | No HLL | Yes | NA | NA | NA | No | No | No | |
| Exp5 | Tecnai F20 | Projection Image | Candidatus Arthromitus sp. strain SFB-rat-Yit | Rattus norvegicus F344 | 1.8 | 388.1 | NA | Stage 1 | No HLL | No | NA | NA | NA | No | No | No | |
| Exp5 | Tecnai F20 | Projection Image | Candidatus Arthromitus sp. strain SFB-rat-Yit | Rattus norvegicus F344 | 30.1 | 510.3 | NA | Stage 4/5_UD | HLL | No | NA | NA | NA | Yes | No | No | |
| Exp5 | Tecnai F20 | Projection Image | Candidatus Arthromitus sp. strain SFB-rat-Yit | Rattus norvegicus F344 | 42.8 | 550.3 | NA | Stage 4/5_UD | HLL | No | NA | NA | NA | No | No | No | |
| Exp5 | Tecnai F20 | Projection Image | Candidatus Arthromitus sp. strain SFB-rat-Yit | Rattus norvegicus F344 | 23.1 | 566.4 | NA | Stage 4/5_UD | HLL | No | NA | NA | NA | Yes | No | No | |
| Exp5 | Tecnai F20 | Projection Image | Candidatus Arthromitus sp. strain SFB-rat-Yit | Rattus norvegicus F344 | 1.5 | NA | NA | UD | NA | No | NA | NA | NA | No | No | Decorated | |
| Exp5 | Tecnai F20 | Projection Image | Candidatus Arthromitus sp. strain SFB-rat-Yit | Rattus norvegicus F344 | 1.5 | NA | NA | UD | UD | No | NA | NA | NA | No | No | Undecorated | |
| Exp5 | Tecnai F20 | Projection Image | Candidatus Arthromitus sp. strain SFB-rat-Yit | Rattus norvegicus F344 | 1.9 | NA | NA | UD | UD | Yes | NA | NA | NA | No | No | Decorated | |
| Exp5 | Tecnai F20 | Projection Image | Candidatus Arthromitus sp. strain SFB-rat-Yit | Rattus norvegicus F344 | 1.9 | NA | NA | UD | UD | Yes | NA | NA | NA | No | No | Undecorated | |
| Exp5 | Tecnai F20 | Projection Image | Candidatus Arthromitus sp. strain SFB-rat-Yit | Rattus norvegicus F344 | 1.9 | NA | NA | UD | UD | No | NA | NA | NA | No | No | Decorated and Undecorated | |
| Exp5 | Tecnai F20 | Projection Image | Candidatus Arthromitus sp. strain SFB-rat-Yit | Rattus norvegicus F344 | 2.0 | NA | NA | UD | No HLL | No | NA | NA | NA | No | No | Decorated | |
| Exp5 | TITAN Krios IP | Projection Image | Candidatus Arthromitus sp. strain SFB-mouse-NL | Mus musculus C57BL/6J | 8.1 | 525.8 | NA | Stage 4 | HLL | No | NA | NA | NA | No | Yes | No | |
| Exp5 | TITAN Krios IP | Projection Image | Candidatus Arthromitus sp. strain SFB-mouse-NL | Mus musculus C57BL/6J | 30.2 | 447.9 | NA | Stage 4/5_UD | HLL | No | NA | NA | NA | No | Yes | No | |
| Exp5 | TITAN Krios IP | Projection Image | Candidatus Arthromitus sp. strain SFB-mouse-NL | Mus musculus C57BL/6J | 12.3 | 533.5 | NA | Stage 5 | HLL | No | NA | NA | NA | No | Yes | No | |
| Exp5 | TITAN Krios IP | Projection Image | Candidatus Arthromitus sp. strain SFB-rat-Yit | Rattus norvegicus F344 | 2.2 | NA | NA | NA | No HLL | NA | NA | NA | NA | NA | NA | NA | |
| Exp6 | TITAN Krios IP | Projection Image | Candidatus Arthromitus sp. strain SFB-mouse-NL | Mus musculus C57BL/6J | 2.4 | 970.5 | NA | Stage 1 | No HLL | No | NA | NA | NA | No | Yes | No | |
| Exp6 | TITAN Krios IP | Projection Image | Candidatus Arthromitus sp. strain SFB-mouse-NL | Mus musculus C57BL/6J | 2.4 | 312.2 | NA | Stage 1 | No HLL | Yes | NA | NA | NA | No | Yes | No | |
| Exp6 | TITAN Krios IP | Projection Image | Candidatus Arthromitus sp. strain SFB-mouse-NL | Mus musculus C57BL/6J | 4.7 | 803.3 | NA | Stage 4 | HLL | No | NA | NA | NA | No | Yes | No | |
| Exp6 | TITAN Krios IP | Projection Image | Candidatus Arthromitus sp. strain SFB-mouse-NL | Mus musculus C57BL/6J | 2.3 | 816.7 | NA | Stage 2 | No HLL | Yes | NA | NA | NA | No | Yes | No | |
| Exp6 | TITAN Krios IP | Projection Image | Candidatus Arthromitus sp. strain SFB-mouse-NL | Mus musculus C57BL/6J | 2.1 | 581.5 | NA | Stage 1 | No HLL | Yes | NA | NA | NA | No | No | No | |
| Exp7 | TITAN Krios IP | Tomogram | Candidatus Arthromitus sp. strain SFB-mouse-NL | Mus musculus C57BL/6J | 2.2 | 588.2 | 162.8 | Stage 1 | NA | No | No | Yes | No | No | No | No | EMD-52685 (tip) |
| Exp7 | TITAN Krios IP | Tomogram | Candidatus Arthromitus sp. strain SFB-mouse-NL | Mus musculus C57BL/6J | 2.5 | 615.5 | 188.4 | Stage 1 | NA | Yes | No | No | No | No | No | Decorated | EMD-52687 (tip) |
| Exp7 | TITAN Krios IP | Tomogram | Candidatus Arthromitus sp. strain SFB-mouse-NL | Mus musculus C57BL/6J | 2.4 | NA | NA | NA | No HLL | Yes | NA | NA | NA | NA | NA | NA | EMD-52696 (back) |
| Exp7 | TITAN Krios IP | Tomogram | Candidatus Arthromitus sp. strain SFB-mouse-NL | Mus musculus C57BL/6J | 2.2 | UD | NA | UD | NA | No | No | Yes | No | No | No | No | EMD-52697 (tip) |
| Exp7 | TITAN Krios IP | Projection Image | Candidatus Arthromitus sp. strain SFB-mouse-NL | Mus musculus C57BL/6J | 2.7 | 905.6 | NA | Stage 4/5_UD | HLL | No | NA | NA | NA | No | No | Decorated and Undecorated | |
| Exp7 | TITAN Krios IP | Projection Image | Candidatus Arthromitus sp. strain SFB-mouse-NL | Mus musculus C57BL/6J | 2.1 | 619.4 | NA | Stage 2 | No HLL | No | NA | NA | NA | No | No | Decorated | |
| Exp7 | TITAN Krios IP | Tomogram | Candidatus Arthromitus sp. strain SFB-mouse-NL | Mus musculus C57BL/6J | 2.0 | 512.1 | NA | Stage 3 | NA | Yes | No | No | No | No | No | No | |
| Exp7 | TITAN Krios IP | Tomogram | Candidatus Arthromitus sp. strain SFB-mouse-NL | Mus musculus C57BL/6J | 2.5 | UD | NA | UD | NA | Yes | No | No | No | No | No | No | |
| Exp7 | TITAN Krios IP | Projection Image | Candidatus Arthromitus sp. strain SFB-mouse-NL | Mus musculus C57BL/6J | 2.4 | UD | NA | UD | NA | No | NA | NA | NA | No | Yes | Decorated | |
| Exp7 | TITAN Krios IP | Projection Image | Candidatus Arthromitus sp. strain SFB-mouse-NL | Mus musculus C57BL/6J | 2.3 | UD | NA | UD | NA | No | NA | NA | NA | No | No | No | |
| Exp7 | TITAN Krios IP | Tomogram | Candidatus Arthromitus sp. strain SFB-mouse-NL | Mus musculus C57BL/6J | 2.3 | 538.9 | NA | Stage 1 | No HLL | No | No | No | No | No | Yes | No | |
| Exp7 | TITAN Krios IP | Tomogram | Candidatus Arthromitus sp. strain SFB-mouse-NL | Mus musculus C57BL/6J | 2.5 | NA | NA | NA | NA | Yes | No | Yes | No | No | NA | No | |

**Supplementary Table 2. Characteristics of vesicles, intracellular filaments and plate-like structures identified at the SFB tip.**

| | | | | Diameter[a]/Width[b] (nm) | | |
|---|---|---|---|---|---|---|
| **SFB origin** | **Feature** | **No. SFB analysed** | **No. features analysed** | **Median** | **Mean** | **Standard deviation** |
| **Mouse-SFB** | Filaments[d] | 9[c] | 18 | 8 | 8 | 1 |
| | Plate-like structures[d] | 3 | 6 | 4 | 4 | 1 |
| | Intracellular vesicles | 10[c] | 15 | 51 | 54 | 15 |
| | Extracellular vesicles | 7[c] | 15 | 90 | 92 | 44 |
| **Rat-SFB** | Filaments[d] | 7[c] | 9 | 9 | 9 | 1 |
| | Plate-like structures[d] | 2 | 4 | 6 | 5 | 1 |
| | Intracellular vesicles | 5[c] | 6 | 41 | 62 | 53 |
| | Extracellular vesicles | 12[c] | 26 | 103 | 103 | 54 |

[a] Diameter: measurement for intracellular and extracellular vesicles. SFB with a ‘broken tip’ phenotype were not included in the intracellular vesicle analysis.
[b] Width: measurement for filaments and plate-like structures.
[c] Data from at least two biological replicates.
[d] Mann Whitney U test revealed statistical significance between the width of filaments and plate-like structures for both mouse-SFB ($p<0.0001$) and rat-SFB ($p<0.0001$).

### Supplementary Table 3

**Supplementary Table 3. Binding affinity between the VHH anti-Th17Ag and Th17Ag determined Biolayer Interferometry (BLI).**

| Association rate constant ($k_{on}$) | Dissociation rate constant ($k_{off}$) | Equilibrium dissociation constant ($K_D$) |
|---|---|---|
| $7.29 \times 10^{3}$ $M^{-1}$ $s^{-1}$ | $2.51 \times 10^{-4}$ $s^{-1}$ | 39 nM |

**Supplementary Table 4. HDX-MS summary table.**

| **HDX EXPERIMENT** | **EPITOPE MAPPING** | |
|---|---|---|
| Data Set | Apo state (Control) | VHH-bound state |
| HDX reaction details | | |
| Labelling buffer : | PBS 1X, pD 7.45 | PBS 1X, pD 7.45 |
| Temperature: | 23°C | 23°C |
| Deuterium excess: | 90% | 90% |
| Molar Excess VHH | N/A | 1.5X |
| % Complex during labelling# | N/A | > 90% |
| HDX time course analyzed (min) | 0.16, 1, 10, 30, 60, and 120 | 0.16, 1, 10, 30, 60, and 120 |
| Number of peptides | 154 | 154 |
| Sequence coverage* | 90.6% | 90.6% |
| Average peptide length | 15.21 | 15.21 |
| Redundancy | 2.44 | 2.44 |
| Average peptide length / Redundancy ratio | 6.23 | 6.23 |
| Replicates | 2 technical replicates & 1 biological replicate | |
| Repeatability (pooled standard deviation) | 0.090 Da | 0.092 Da |
| Significant difference between state^ | Wald test, $p<0.05$ | |

#considering a Kd of 39 nM and a 1 to 1 binding stoichiometry
*sequence coverage after labelling with 1.5X molar excess VHH
^MEMHDX software[55]
N/A: not applicable

### Supplementary Figures

### Supplementary Fig. 1



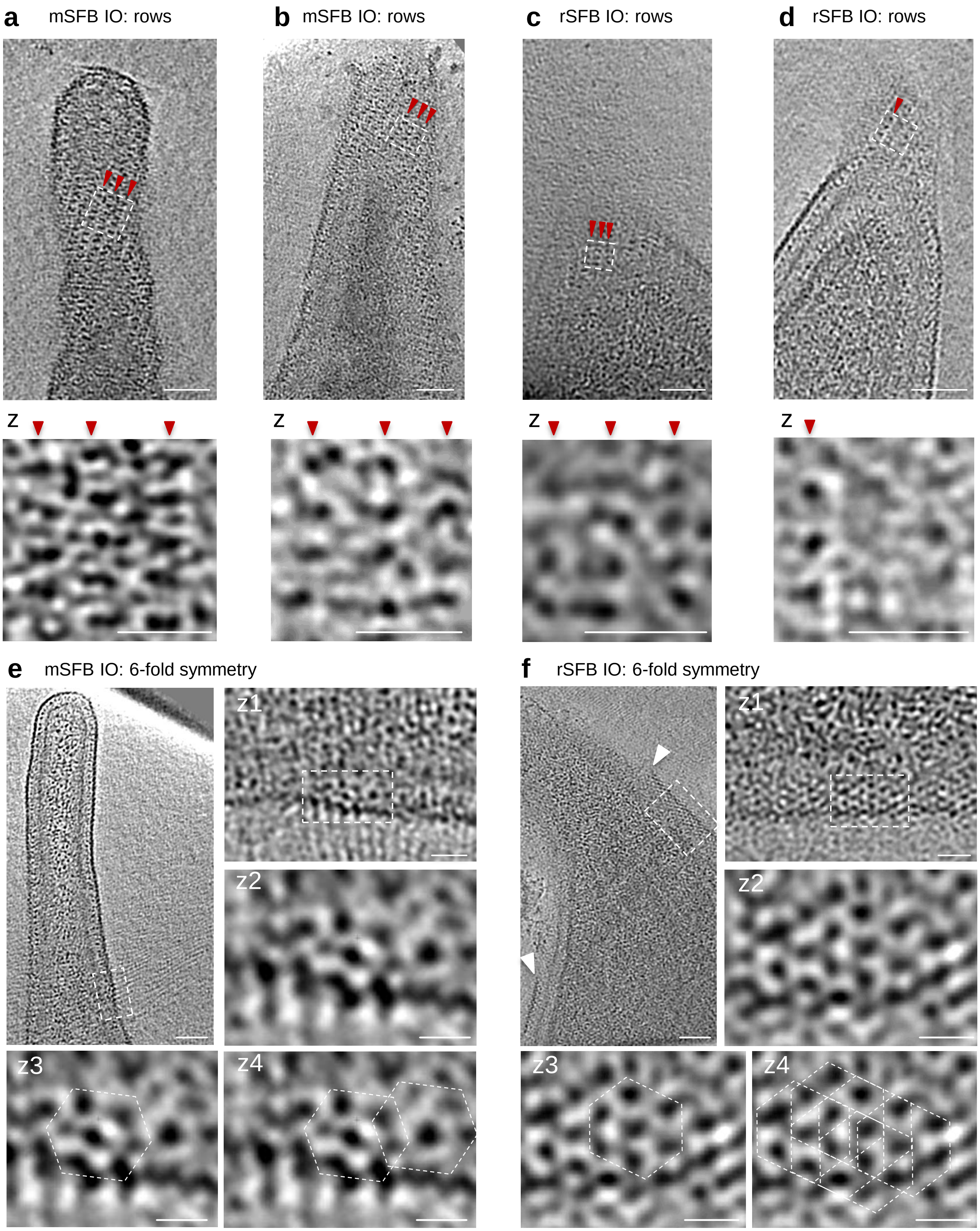

**Supplementary Fig. 1. Top view of the S-layer arrangement at the SFB tip. a-c,** Top view of the organization of S-layer subunits in rows seen in tomograms from the tip of **(a,b)** mouse-SFB (EMD-52685, EMD-52687) and **(c,d)** rat-SFB IOs (EMD-52688,EMD-52684). Close ups (z) of the regions delimited by a white dashed line were included below each panel. Red arrows show S-layer subunits arranged in rows. **d,e,** Top view of tomograms which include the tip of **(e)** mouse-SFB (EMD-52655) and **(f)** rat-SFB IOs (EMD-52689), showing a potential six-fold symmetry of the S-layer subunits. Close ups (z) of the regions delimited by a white dashed line were included below and next to each panel. Regions where a potential 6-fold symmetry was identified are shown by white dashed hexagons in **(z3)** one and in **(z4)** multiple regions of the tomogram. S-layer discontinuities at the SFB tip resulting in a 'broken tip' phenotype are shown by white arrows. mSFB: mouse-SFB, rSFB: rat-SFB. **Scale bars:** a-f**:** 50 nm; a-f(z): 20nm.

### Supplementary Fig. 2

Cruz, *et al.*, 2025

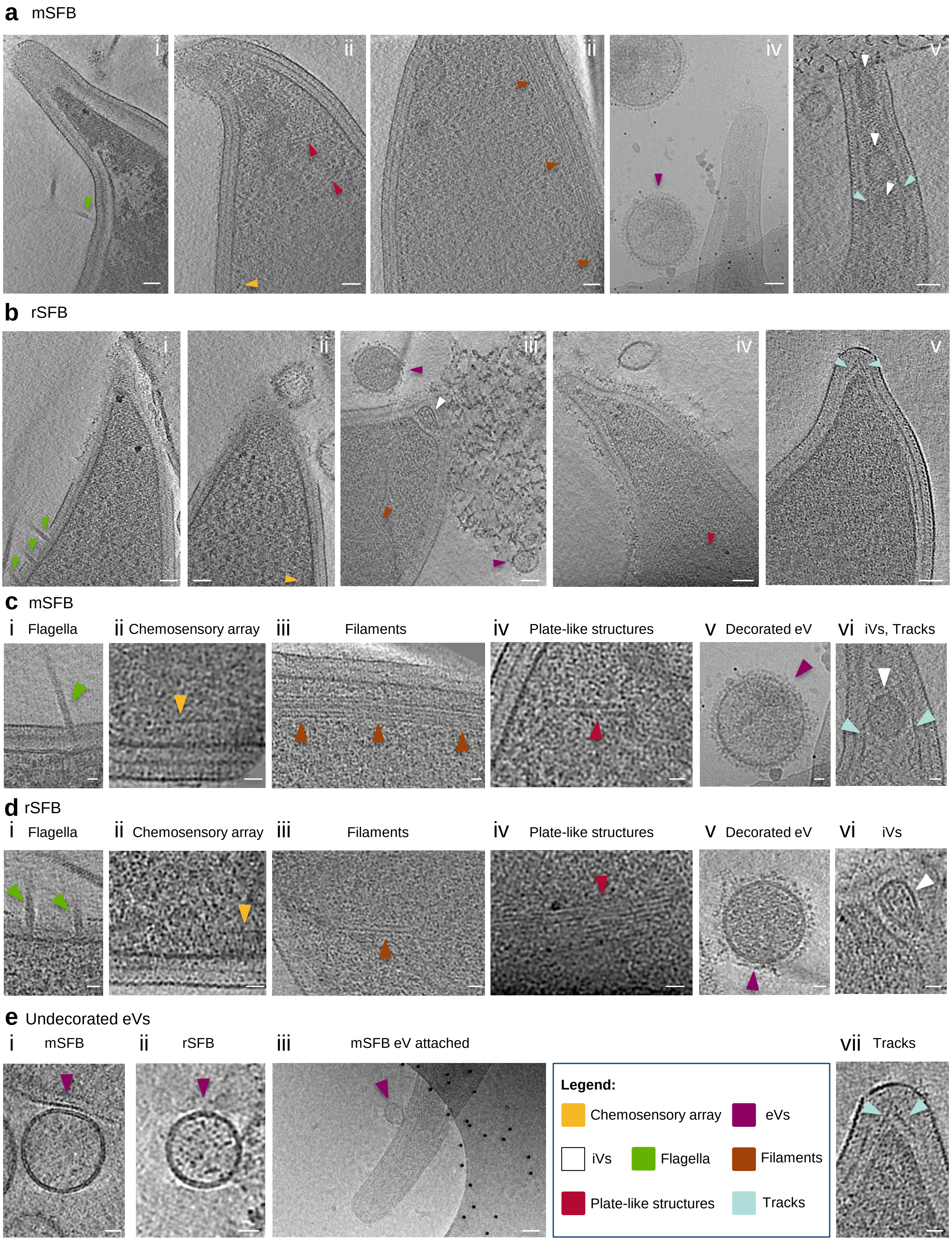

**Supplementary Fig. 2. Additional examples of the intracellular and extracellular features identified in SFB and related to Figure 2. a(i-iii,v),** Representative tomographic slices from mouse-SFB IOs tomograms EMD-52687, EMD-52673, EMD-52674 and EMD-52670 containing intracellular and extracellular features. The corresponding IOs had the following length: 2.5, 2.3, 2.5, 2.1, 2.5 µm. **a(iv)** Representative projection image from mouse-SFB containing decorated extracellular vesicles. **b(i-ii)**, representative tomographic slices showing **b(i)** flagella and **b(ii)** a chemosensory array present in an IO with 2.2 µm in length (EMD-52692). **b(iii-v),** Representative tomographic slices from rat-SFB IOs tomograms EMD-52693, EMD-52694 and EMD-52695 showing **b(iii)** filaments, iVs and eVs **b(iv)** plate-like structures and **b(v)** tracks. The corresponding IOs had the following length: 2.0, 2.0 and 2.2 µm. **c,d,** Close ups from the tomographic slices of **(c)** mouse-SFB and **(d)** rat-SFB shown in a and b, respectively, that contain the following features: **c-d(i)** flagella, **c-d(ii)** chemosensory array, **c-d(iii)** filaments, **c-d(iv)** plate-like structures, **c-d(v)** decorated extracellular vesicles (eVs), **c-d(vi)** intracellular vesicles and **c(vi),d(vii)** tracks. **e(i-ii)**, Tomographic slices of undecorated eVs found near **(e(i))** mouse-SFB (EMD-52675) and **(e(ii))** rat-SFB IOs (EMD-52693). **e(iii)**, Projection image containing an extracellular vesicle in direct contact with the tip of a mouse-SFB IO. The image was acquired with a Tecnai F20 electron microscope equipped with a Falcon 2 camera. The identified features are shown by arrows of the colours indicated in the legend. mSFB: mouse-SFB, rSFB: rat-SFB, iVs: intracellular vesicles, eVs: extracellular vesicles. **Scale bars:** a-b, e(iii): 50 nm; c/d, e(i-ii): 20 nm.

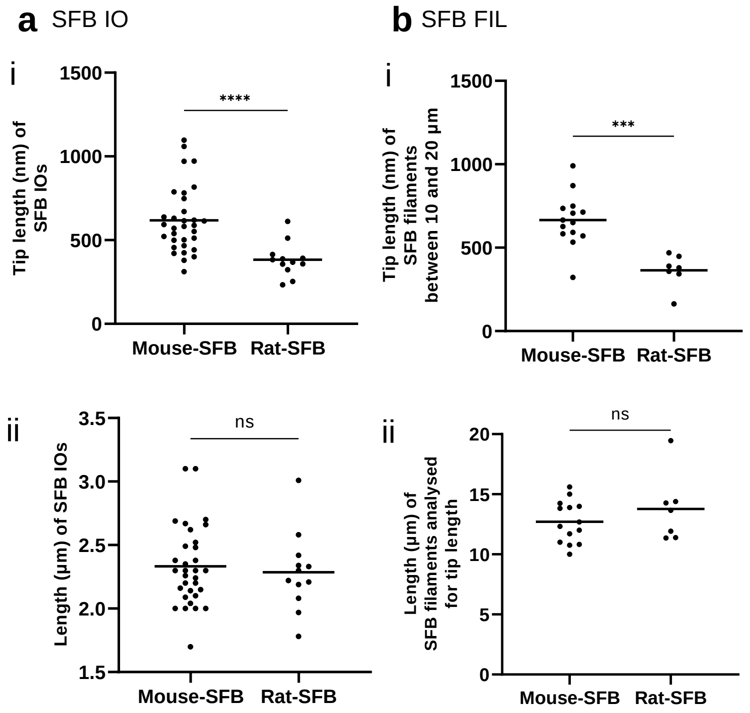


**Supplementary Fig. 3. Differences in tip length between mouse-SFB and rat-SFB. a,b(i),** Tip length of mouse-SFB and rat-SFB **a(i)** IOs and **b(i)** filaments. **a,b(ii),** Length of mouse-SFB and rat-SFB **a(ii)** IOs and **b(ii)** filaments included in the tip length analysis. Only IOs assigned to stages 1-3 were included in the analysis (32 mouse-SFB IOs and 12 rat-SFB IOs) of a(ii). Only filaments with a length between 10 and 20 µm were included in the analysis (14 mouse-SFB filaments and 7 rat-SFB filaments) of b(ii). Individual measurements and the corresponding mean are shown. The statistical significance was assessed using the Mann–Whitney U test (a(i): $p$<0.0001, b(i): $p$=0.0005, a,b(ii): ns: not significant).

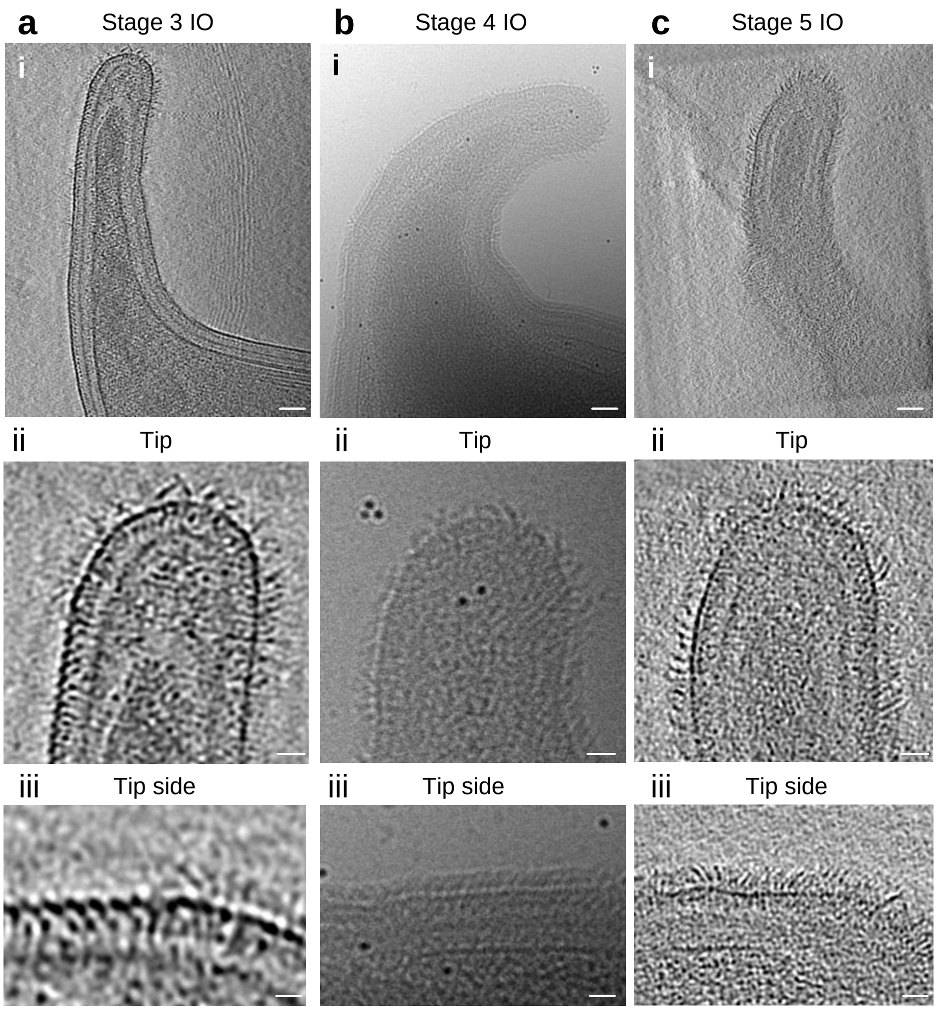


**Supplementary Fig. 4. Additional examples of the developmental stages of mouse-SFB IOs, related to figure 3. a-c,** Representative **(a,c)** tomographic slices from reconstructed tomograms EMD-52680 and EMD-52682 and **(b)** projection image showing the tip of mouse-SFB IOs assigned to stages **a(i)** 3, **b(i)** 4 and **c(i)** 5. The selected IOs had a length of 2.3, 4.7 and 4.8 µm, respectively. **a-c(ii-iii),** Close ups of the **a-c(ii)** IOs tip and **a-c(iii)** tip side from stages 3, 4 and 5, respectively. **Scale bars:** a-c(i): 50 nm; a-c(ii,iii): 10 nm.

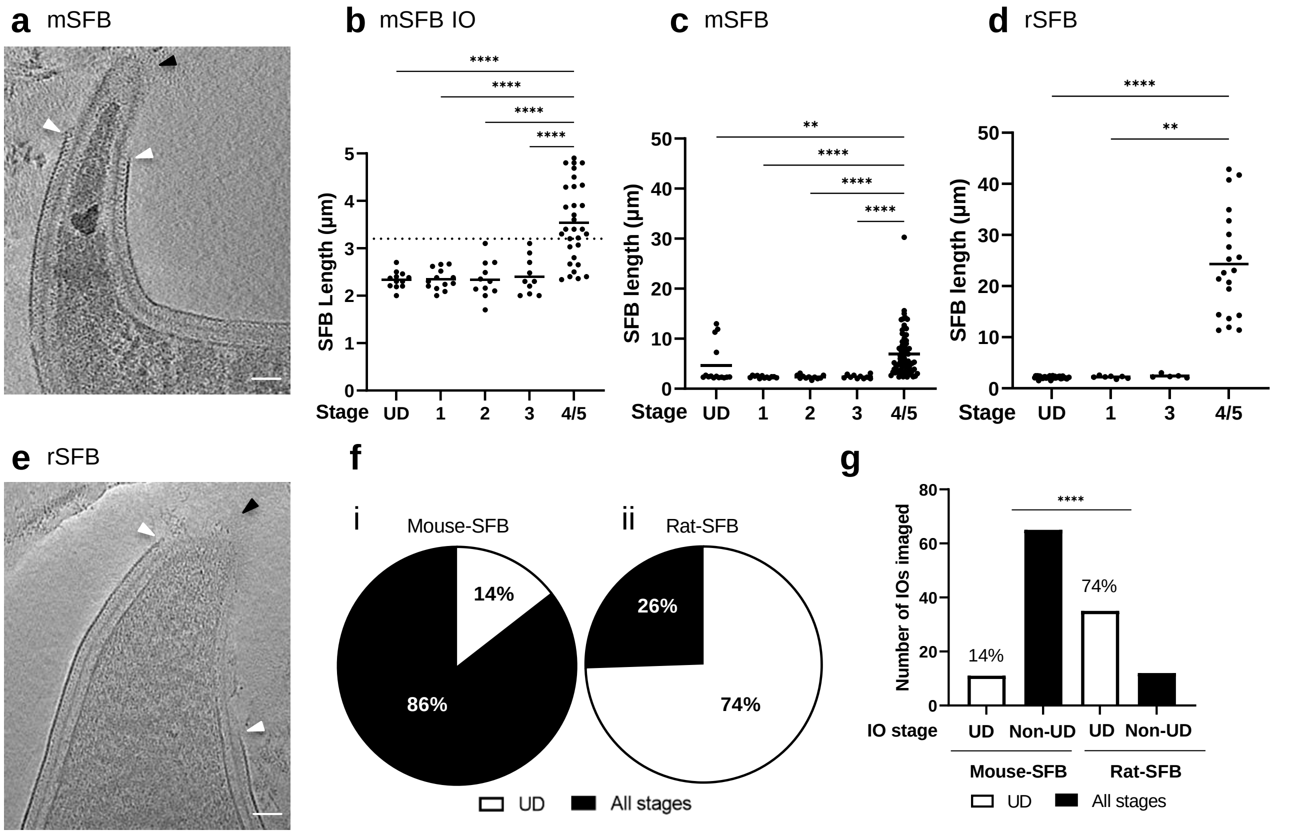


**Supplementary Fig. 5. Identification of undefined SFB stages. a/e,** Tomographic slices of representative **(a)** mouse-SFB (EMD-52697) and **(e)** rat-SFB IOs (EMD-52690) that could not be assigned to a tip stage (UD). Discontinuities in the IOs S-layer are shown by white arrows. The absence of disHLS at the tip end is shown by black arrows. **b-d,** Length of: **(b)** mouse-SFB IOs grown in mice, **(c)** total mouse-SFB grown in mice and **(d)** total rat-SFB grown in rats assigned to each tip stage. All SFB imaged were included in the analysis even if no distinction between stages 4 and stage 5 could be made (4/5) and if the tip stage could not be identified (UD, undetermined) (a: n=123, 7 independent experiments; b: n=77, 7 independent experiments; c: n=63, 5 independent experiments; d: n=79, 4 independent experiments). Individual measurements and the corresponding mean are shown. For panel a, the statistical significance was assessed using the one-way ANOVA ($p$<0.0001). A dashed line was included at an IO length of 3.2 µm. For panels b and c, the statistical significance was assessed using the Kruskal-Wallis test followed by a Dunn's test correction for multiple comparisons (b: Stage 4/5 vs Stage1, Stage 2 and Stage 3: $p$<0.0001, Stage 4/5 vs Stage UD: $p$= 0.0077; c: Stage 1 vs Stage 4/5, $p$=0.0014; UD vs Stage 4/5, $p$<0.0001). **f,** Proportion of **(i)** mouse-SFB and **(ii)** rat-SFB IOs assigned to a tip stage. mSFB: mouse-SFB, rSFB: rat-SFB. **g**, Number of mouse-SFB and rat-SFB IOs in which the tip stage could not be determined due to discontinuities in the S-layer. The statistical significance between the proportion of the tip stages of mouse-SFB and rat-SFB IOs was assessed using the Fisher's exact test ($p$<0.0001). **Scale bar**: a/e: 50 nm.

### Supplementary Fig. 6

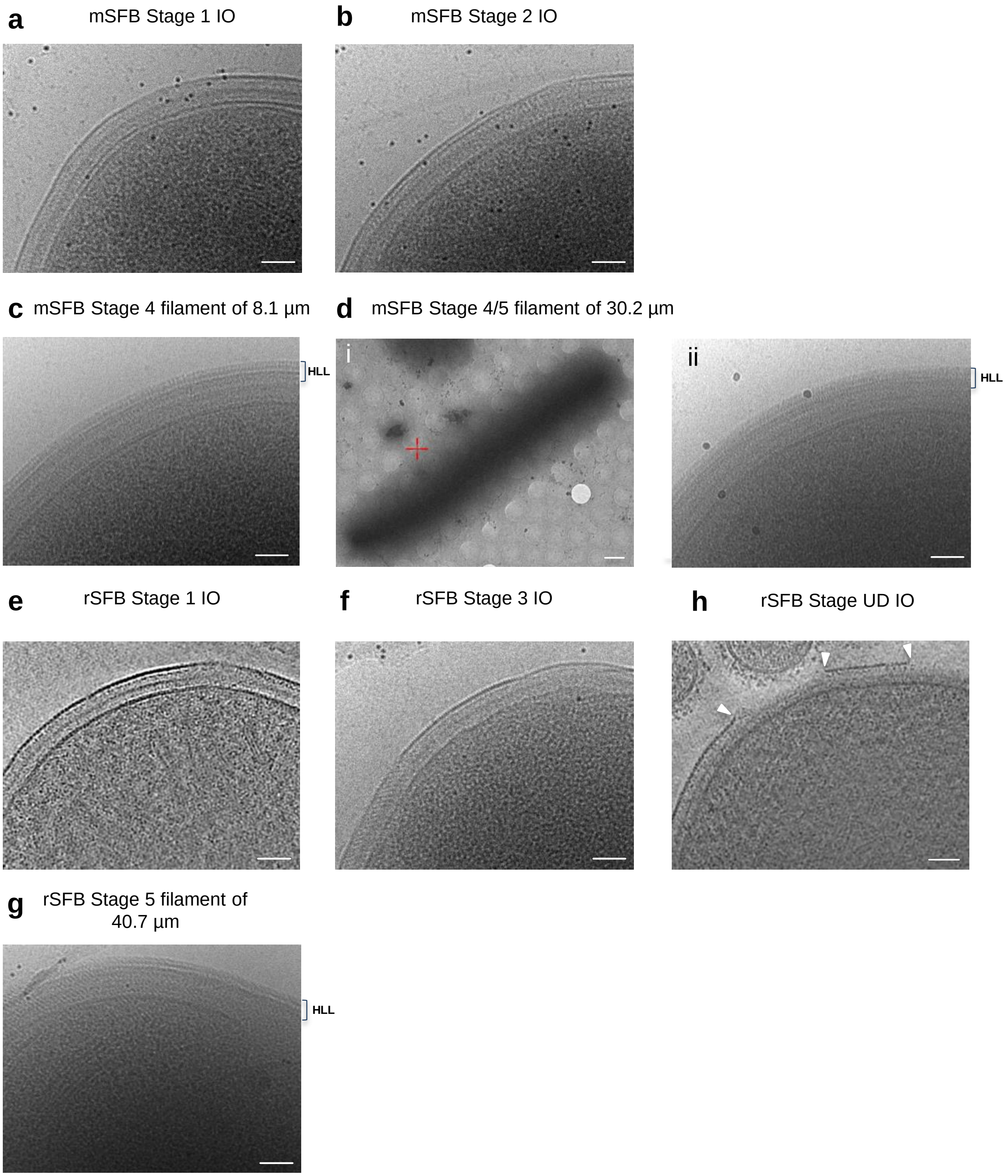


**Supplementary Fig. 6. Assessment of the presence of the hair-like layer at the SFB back. a-b,** Representative projection images of the back of mouse-SFB IOs assigned to stages **(a)** 1 and **(b)** 2. **c-d**, Representative projection images of mouse-SFB filaments assigned to stages **(c)** 4 (8.1 µm filament length) and **(d) 4/5** (30.2 µm filament length). **d(i)**, A long SFB filament (30.2 µm filament length) and the back of mouse-SFB filaments assigned to stages **(c)** 4 and **(d(ii)) 4/5** are shown. **e,** Representative tomographic slice of the back of rat-SFB IOs assigned to stage 1 (EMD-52698). **f,** Representative projection image of the back of rat-SFB IOs assigned to stage 3. **g**, Representative projection image of the back of a rat-SFB filament assigned to stage 5 (40.7 µm filament length). The hair-like layer (HLL) is delimited when present. **h**, Representative tomographic slice from the back of a rat-SFB IO with an undefined (UD) displaying S-layer discontinuity also at the back (highlighted with white arrows) (EMD-52856). **Scale bars:** a-c, d(ii), e-h: 50nm; d(i): 2 µm.

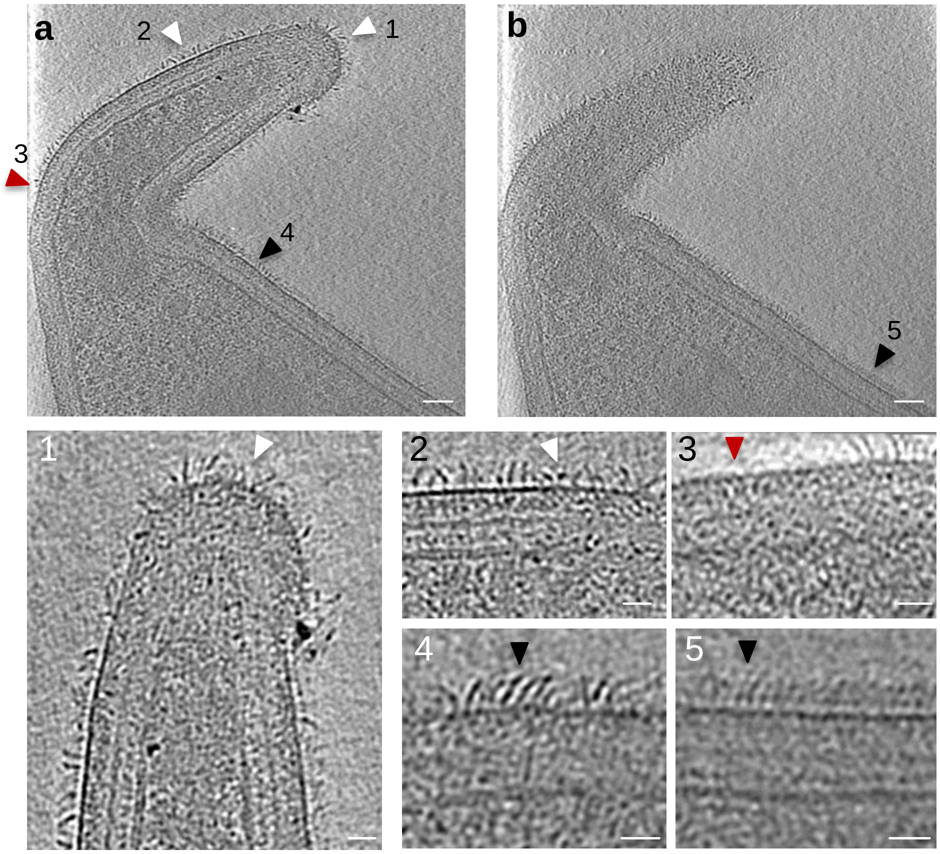


**Supplementary Fig. 7. Potential transitional stage from stage 3 to stage 4. a-g,** Tomographic slices from tomogram EMD-52683. Tomographic slices at **(a,b)** different tilts showing the tip and beginning of the cell body of a mouse-SFB IO (3.3 µm in length) that includes: **(1,2)** disordered hair-like structures (white arrows), **(3)** regions without HLS (red arrow), and **(4,5)** a hair-like layer (black arrows). The regions from the which close ups were taken (1-5) are indicated in panels a and b by arrows labelled with the letter of the corresponding panel. **Scale bars**: a/b: 50 nm; 1-5: 20 nm.

### Supplementary Fig. 8



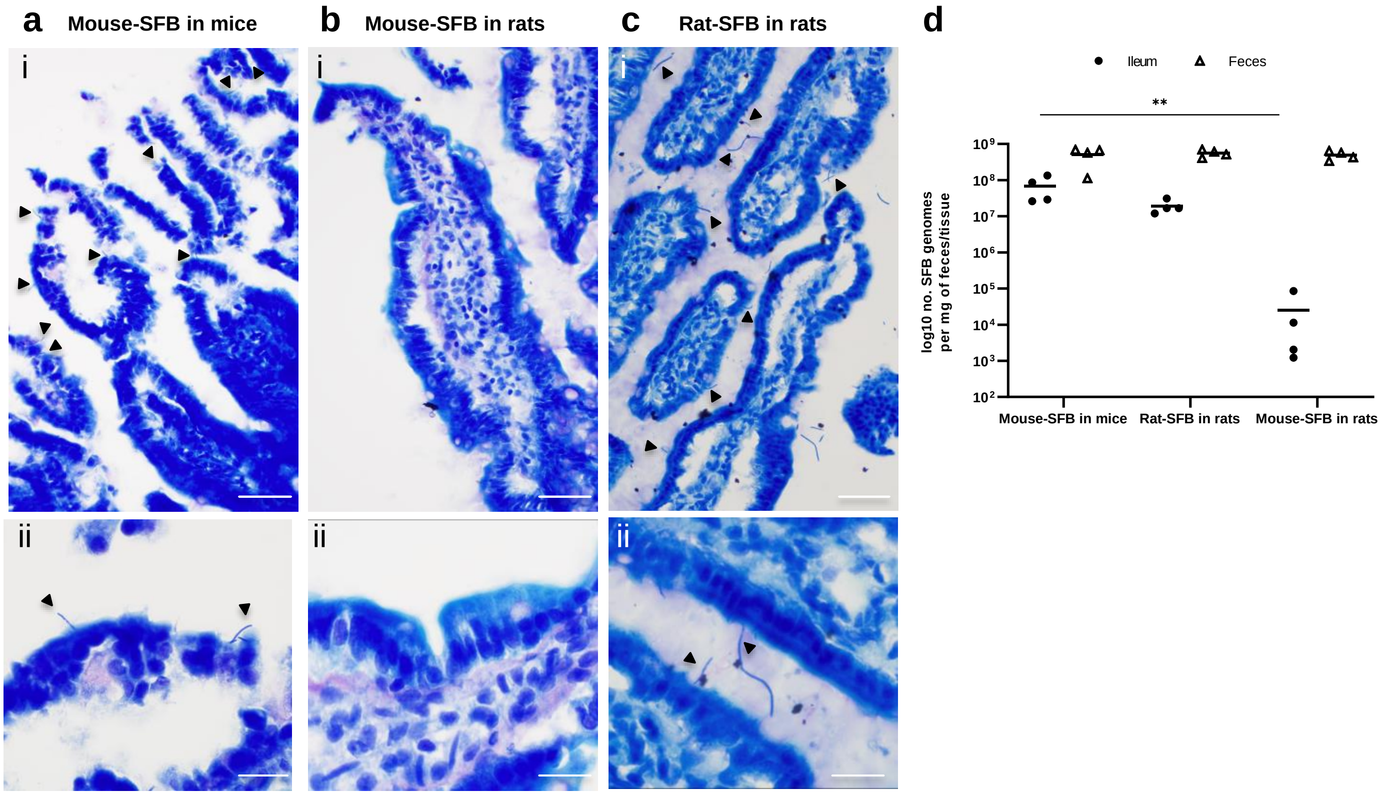


**Supplementary Fig. 8. Assessment of host-specificity of SFB attachment. a-c,** Giemsa stain of the terminal ileum of **(a)** mice colonized by mouse-SFB, **(b)** rats colonized by mouse-SFB, and **(c)** rats colonized by rat-SFB. **a,c(ii),** Close ups of attached SFB. **b(ii)**, Close up of villi where no attached SFB was found. Attached SFB are highlighted with black arrows. **d**, SFB quantification in feces and terminal ileum biopsies of monocolonized mice and rats by qPCR. Data from 4 biological experiments are shown. The statistical significance was assessed using the Kruskal-Wallis test followed by Dunn's test correction for multiple comparisons (ileum mouse-SFB in mice vs ileum mouse-SFB in rats, $p$=0.0098, the remaining comparisons were not significant). **Scale bars:** a-c(i): 2000 µm; a-c(ii): 50 µm.

### Supplementary Fig. 9



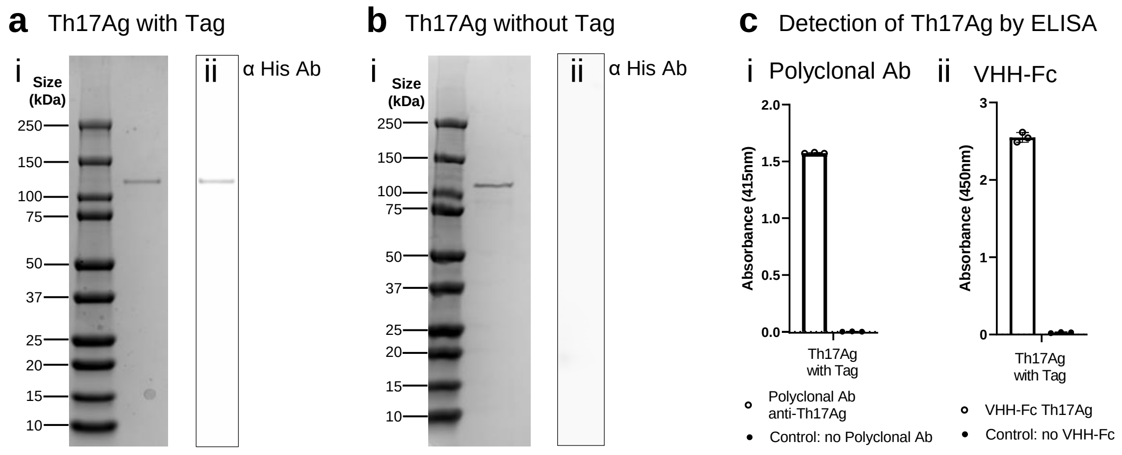


**Supplementary Fig. 9. Purification and detection of a Th17 antigen (Th17Ag). a-b(i),** SDS-PAGE gel stained with Coomassie brilliant blue R-250 showing the purified Th17Ag expressed in *Escherichia coli* BL21 DE3 Star **(a)** before and **(b)** after the cleavage of the N-terminal 6xHis and V5 epitope-containing tag. **a-b(ii)**, Western blot of Th17Ag using an anti-polyhistidine antibody in the conditions described for a-b(i), respectively. A band corresponding to the tagged protein can be seen before the cleavage of the N-terminal tag **(a(ii))** but not after the cleavage of the N-terminal tag **(b(ii))**. Molecular weight of the tagged and untagged protein estimated by Expasy Server[58] is 114 kDa and 110 kDa, respectively. **c**, Detection by enzyme-linked immunosorbent assay (ELISA) of the purified Th17Ag. Binding was detected using: **c(i)** a rabbit polyclonal antibody and a secondary anti-rabbit antibody conjugated with alkaline phosphatase and **c(ii)** a nanobody (VHH) coupled to the Fc of the human IgG1 and a secondary antibody anti-Human IgG Fc fragment conjugated with horseradish peroxidase. The mean and standard deviation of three replicates are shown.

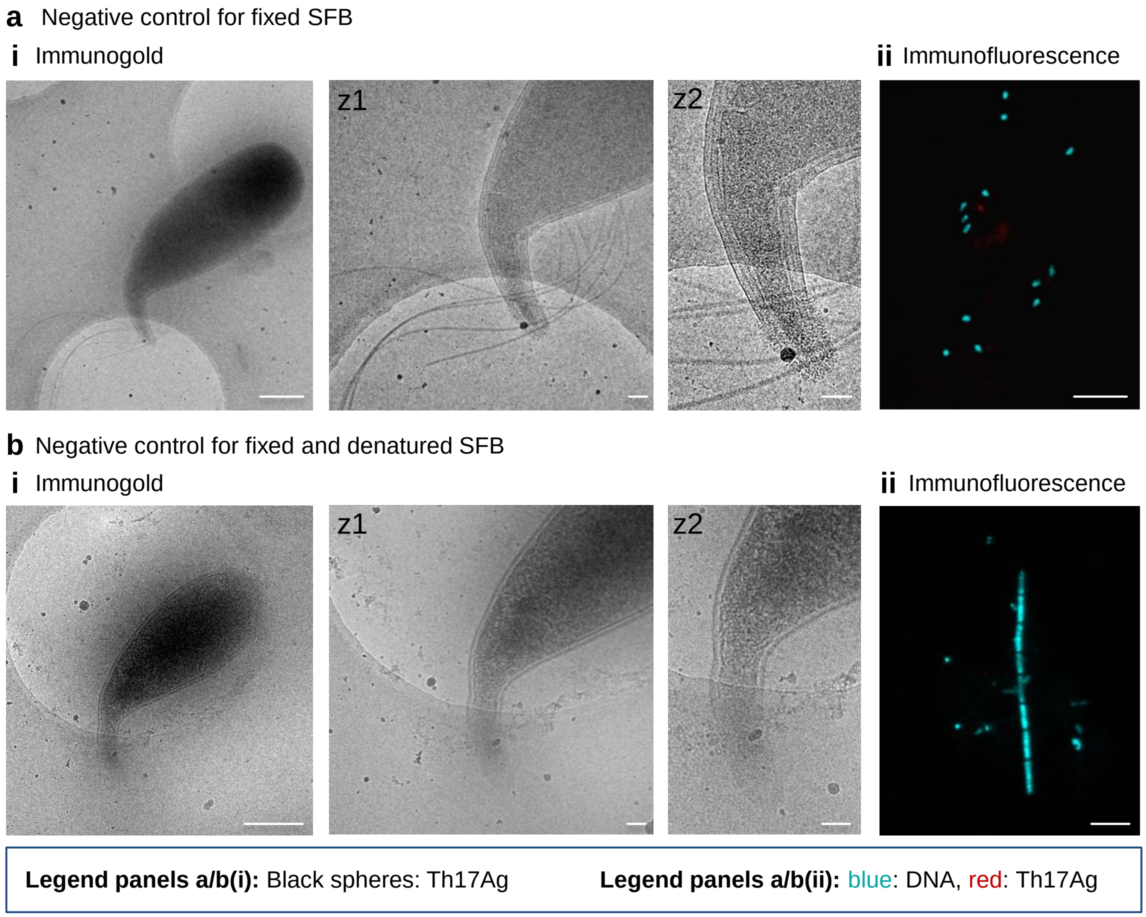


**Supplementary Fig. 10. Negative controls of immunofluorescence and immunogold labelling. a,** Labelling of fixed SFB with **(a(i))** gold-conjugated Protein A or with **(a(ii))** DAPI and secondary antibody anti-Human IgG Fc Fragment conjugated with Alexa 568. **b**, Labelling of fixed and denatured SFB with **(b(i))** Protein A conjugated with 5 nm gold particles or with **(b(ii))** DAPI and secondary antibody anti-rabbit conjugated with Alexa 568. Close ups (z) are shown for the bacteria included in panels a/b. Projection images of immunogold labelled SFB were acquired with a Tecnai F20 electron microscope equipped with a Falcon 2 camera. For all immunogold and immunofluorescence experiments, labelling was performed in two distinct days using two biological replicates for each experiment. **Scale bars:** a-b(i): 500nm; a/b(z): 100 nm; a-b(ii): 5 µm.

### Supplementary Fig. 11



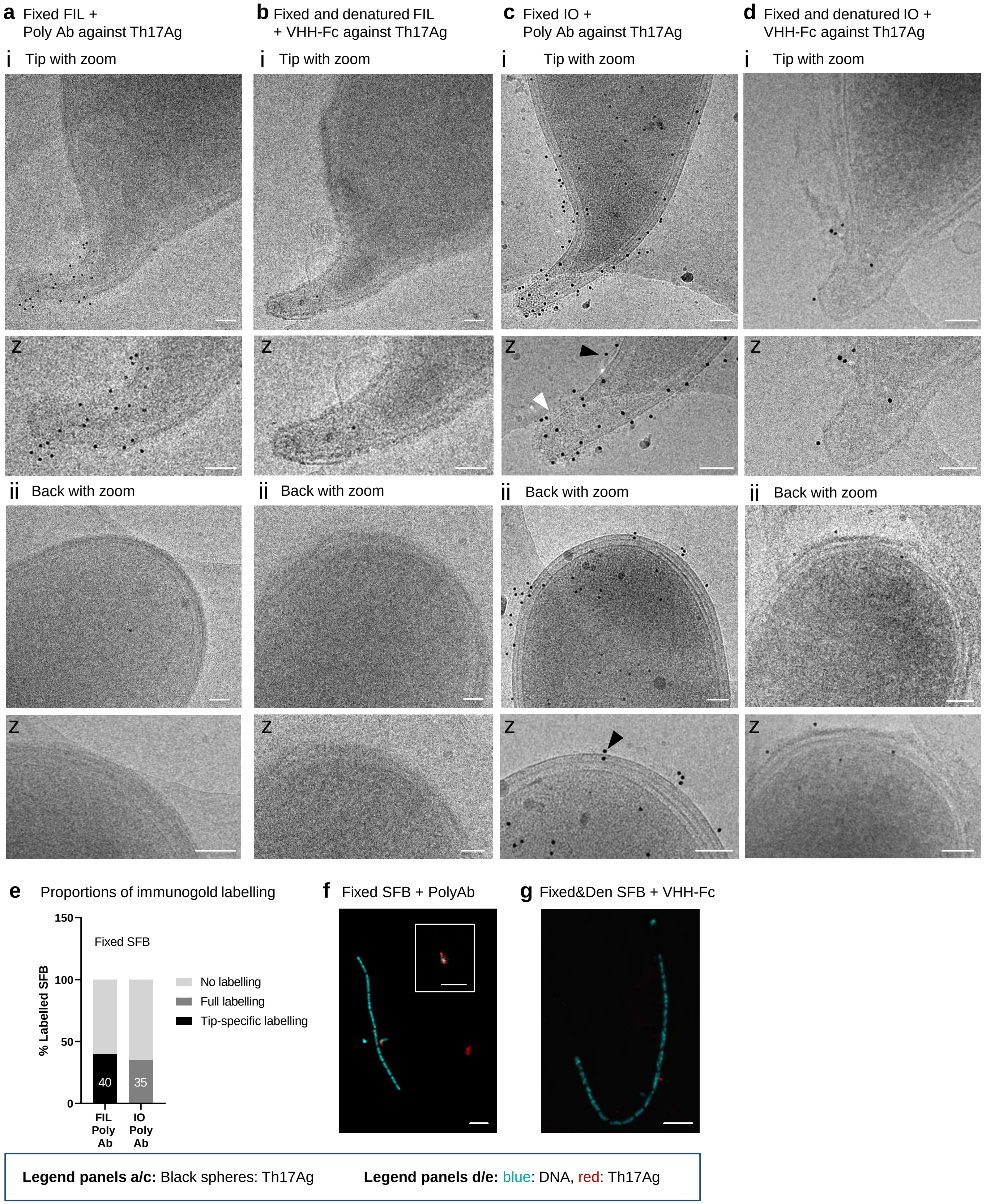

**Supplementary Fig. 11. Additional data for the localization of Th17Ag at the SFB surface. a-d**, Projection images from purified SFB **(a,b)** filaments (FIL) and **(c,d)** IOs stained using immunogold labelling. Fixed and denatured SFB **(a)** filaments and **(c)** IOs were incubated with a VHH-Fc anti-Th17Ag. Fixed SFB **(b)** filaments and **(d)** IOs were incubated with a rabbit polyclonal antibody against Th17Ag. Gold-conjugated Protein A was used for immunogold labelling. Imaging was performed at the SFB **(a-d(i))** tip and **(a-d(ii))** back. Close ups (z) of the SFB tip and back are shown under each panel. Examples of labelled regions both with and without visible hair-like structures are shown by white and black arrows, respectively. Projection images were acquired with a Tecnai F20 electron microscope equipped with a Falcon 2 camera. **e,** Assessment of immunogold labelling for SFB imaged in the condition described for panel d. The percentage of SFB labelled and labelled specifically at the tip are indicated on the corresponding bar. SFB were considered labelled if co-localization with at least 20 gold particles (black spheres) was observed, or at least 10 gold particles if labelling was restricted to the SFB tip. Between 10 and 20 SFB were imaged for each condition**. f,g,** Immunofluorescence images of **(f)** fixed and denatured SFB incubated with a biotinylated VHH-Fc anti-Th17Ag and Streptavidin-Alexa568 and of **(g)** fixed SFB incubated with a rabbit polyclonal antibody (Ab) against Th17Ag and a secondary antibody anti-rabbit conjugated with Alexa 568. All SFB were additionally labelled with DAPI. An extra image of a labelled IO was included as an insert delimited by a white line (f). For all immunogold and immunofluorescence experiments, labelling was performed in two distinct days using two biological replicates for each experiment. **Scale bars**: a-d: 100 nm; f/g: 5 µm.

### Supplementary Fig. 12



**a** Kinetic analysis of Th17Ag binding to monovalent VHH

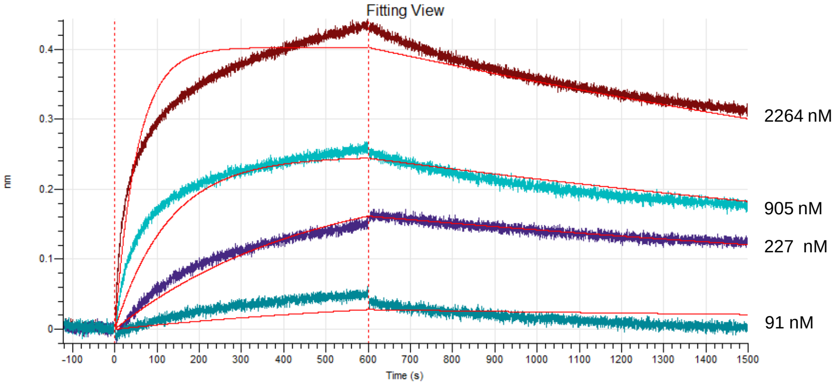


**b** Peptide map of Th17Ag

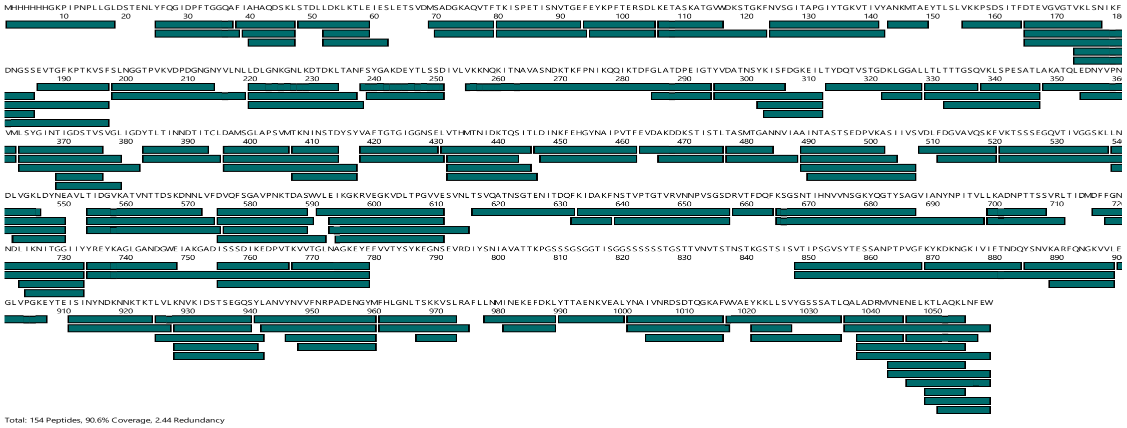


**c** Deuterium uptake profiles and differential uptake profile

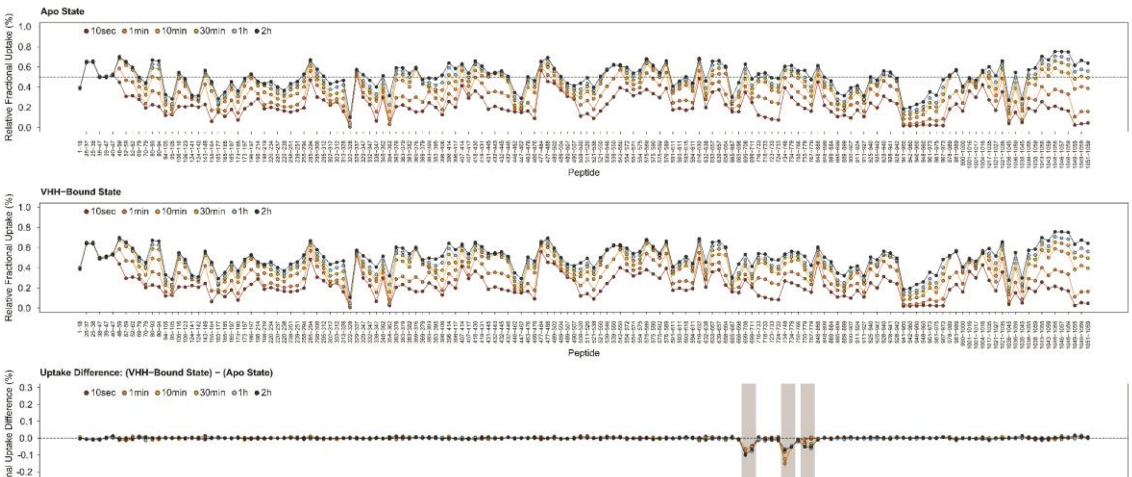


**Supplementary Fig. 12. Kinetic and HDX-MS analysis of Th17Ag binding to monovalent VHH anti-Th17Ag. a,** Kinetic analysis of Th17Ag binding to monovalent anti-TH17Ag VHH. Biolayer Interferometry (BLI) was performed using an Octet HTX system. The resulting sensorgrams showing association and dissociation curves of Th17Ag to immobilized monovalent VHH are shown. The concentrations of Th17Ag used for the binding assays are indicated next to each curve. The fitting curves obtained with a 1:1 Langmuir model are shown in red. **b**, Peptide map of Th17Ag. Each bar corresponds to a unique peptide selected for analysis by Hydrogen/ Deuterium eXchange-Mass Spectrometry (HDX-MS). A total of 154 peptides covering 90.6% of the protein sequence with a 2.44 redundancy were recovered. **c,** Effect of VHH binding on the exchange behaviour of Th17Ag determined by HDX-MS. The deuterium uptake profiles of Th17Ag alone (Apo State) and Th17Ag bound to VHH (VHH-Bound State) were used to calculate the uptake difference between the two protein states, shown by a differential fractional uptake plot. A negative value is indicative of a VHH protective effect.

#### Supplementary Movie Legends

**Supplementary Movie 1. Representative tomogram of a mouse-SFB IO tip (EMD-52655) with the corresponding segmentation showing the S-layer, cell wall, membrane, flagella, intracellular vesicles and tracks. Scale bar:** 50nm

**Supplementary Movie 2. Representative tomogram of a mouse-SFB IO tip (EMD-52667) with the corresponding segmentation showing the S-layer, cell wall, membrane and chemosensory array. Scale bar:** 50nm

**Supplementary Movie 3. Representative tomogram of a mouse-SFB IO tip (EMD-52668) with the corresponding segmentation showing the S-layer, cell wall, membrane and representative clustered filaments. Scale bar:** 50nm

**Supplementary Movie 4. Representative tomogram of a mouse-SFB IO tip (EMD-52669) with the corresponding segmentation showing the S-layer, cell wall, membrane and representative plate-like structures. Scale bar:** 50nm

**Supplementary Movie 5. Representative tomogram of a mouse-SFB IO tip (EMD-52670) with the corresponding segmentation showing the S-layer, cell wall, membrane, undecorated extracellular vesicles, intracellular vesicles and tracks. Scale bar:** 50nm

**Supplementary Movie 6. Representative tomogram of a Stage 1 mouse-SFB IO tip (EMD-52685) with the corresponding segmentation showing the S-layer, cell wall and membrane. Scale bar:** 50nm

**Supplementary Movie 7. Representative tomogram of a Stage 3 mouse-SFB IO tip (EMD-52676) with the corresponding segmentation showing the S-layer, cell wall, membrane and disHLS. Scale bar:** 50nm

**Supplementary Movie 8. Representative tomogram of a Stage 4 mouse-SFB filament tip (EMD-52677) with the corresponding segmentation showing the HLL, cell wall, membrane, disHLS and ordHLS. Scale bar:** 50nm

**Supplementary Movie 9. Representative tomogram of a Stage 5 mouse-SFB filament tip (EMD-52678) with the corresponding segmentation showing the HLL, cell wall, membrane and ordHLS. Scale bar:** 50nm
